# Silybin A from *Silybum marianum* reprograms lipid metabolism to induce a cell fate-dependent class switch from triglycerides to phospholipids

**DOI:** 10.1101/2024.04.09.588052

**Authors:** Solveigh C. Koeberle, Maria Thürmer, Markus Werner, Julia Grander, Laura Hofer, André Gollowitzer, Fengting Su, Loc Le Xuan, Felix J. Benscheid, Ehsan Bonyadi Rad, Armando Zarrelli, Giovanni Di Fabio, Oliver Werz, Valeria Romanucci, Amelie Lupp, Andreas Koeberle

## Abstract

*Silybum marianum* is used to protect against degenerative liver damage. The molecular mechanisms of its bioactive component, silybin, remained enigmatic, although membrane-stabilizing properties, modulation of membrane protein function, and metabolic regulation have been discussed for decades. Here, we show that specifically the stereoisomer silybin A decreases triglyceride levels and lipid droplet content, while enriching major phospholipid classes and maintaining a homeostatic phospholipid composition in human hepatocytes *in vitro* and in mouse liver *in vivo* under normal and pre-disease conditions. Conversely, in cell-based disease models of lipid overload and lipotoxic stress, silybin treatment primarily depletes triglycerides. Mechanistically, silymarin/silybin suppresses phospholipid-degrading enzymes, induces phospholipid biosynthesis to varying degrees depending on the conditions, and down-regulates triglyceride biosynthesis, while inducing complex changes in sterol and fatty acid metabolism. Structure-activity relationship studies highlight the importance of the 1,4-benzodioxane ring configuration of silybin A in triglyceride reduction and the saturated 2,3-bond of the flavanonol moiety in phospholipid accumulation. Enrichment of hepatic phospholipids and intracellular membrane expansion are associated with an heightened biotransformation capacity. In conclusion, our study deciphers the structural features of silybin contributing to hepatic lipid reorganization and offers insights into its liver-protective mechanism, potentially involving a context-dependent lipid class switch from triglycerides to phospholipids.

## 1. Introduction

Hepatic pathologies such as metabolic dysfunction-associated steatotic liver disease (MAFLD; former: non-alcoholic fatty liver disease, NAFLD^1^), metabolic dysfunction-associated steatohepatitis (MASH; former: non-alcoholic steatohepatitis, NASH), fibrosis, and cirrhosis are closely related to the metabolic syndrome and insulin resistance^2-9^. They are driven by high-calorie diets that induce abnormal glucose and lipid metabolism and subsequently cause glucotoxicity, lipotoxicity, oxidative stress, and chronic inflammation^2,6,10-12^. As a consequence, fatty acids are taken up by hepatocytes, and also synthesized *de novo*^13^, incorporated into triglycerides (TGs), and stored in lipid droplets^14^. While the transfer of fatty acids into lipid droplets contributes to the detoxification of excess free fatty acids^14^, a chronic increase in the number and size of lipid droplets induces hepatocyte enlargement and dysfunction^10,15^. This continuous lipid accumulation leads to hepatic steatosis and, as the disease progresses, to cirrhosis and hepatocellular carcinoma^2,16^. As an adaptive strategy to protect hepatocytes from lipid overload, autophagy of lipid droplets (lipophagy) is initiated^17^ and the mobilized fatty acids are subjected to oxidative degradation^18^. Compensatory upregulation of fatty acid oxidation at the onset of MAFLD provides partial relief but is insufficient to reduce hepatic lipids to basal levels. In addition, the increased oxidative breakdown of lipids induces oxidative stress, which can negatively contribute to cell and tissue damage^10,19^. MAFLD is also significantly influenced by genetic factors^20^. Candidate gene variants act in multiple pathways of lipid metabolism^21^, including *de novo* lipogenesis and lipid droplet assembly (LPIN2, ATGL/PNPLA2)^22,23^, phospholipid biosynthesis and remodeling (LPIAT1/MBOAT7, iPLA2/PLA2G6, PNPLA8, PRDX6, PLD1)^24-30^, neutral and phospholipid The flavonolignan silybin favorably redistributes lipids hydrolysis and catabolism (PNPLA3)^31^, sterol metabolism (HSD17B13)^32^, fatty acid compartmentalization (GCKR, TM6SF2), and lipoprotein assembly and secretion (PLA2G7, TM6SF2)^27^. Consequently, both MAFLD and MASH are characterized by extensive changes in hepatic lipid composition, including a decrease in total phosphatidylcholine (PC) and an increase in TG^33-41^.

Milk thistle (*Silybum marianum* L.) is a medicinal plant that is traditionally used for the treatment of liver and biliary tract diseases^42-45^ and a variety of other pathologies, including diabetes^46^ and cancer^47,48^. Organic fruit extracts (silymarin) of *S. marianum* consist of the flavonolignans silybin A and B (∼30%), isosilybin (∼5%), silychristin A (∼7%), and silydianin (∼10%), the flavonoid (+)-taxifolin (∼5%) (Figure 1A), and less defined polyphenols (30%)^47,49^. Minor constituents include silychristin B, isosilychristin, 2,3-dehydrosilybin, quercetin, and kaempferol^47,49,50^. The major biologically active flavonolignan, silybin, also termed as silibinin, exists as a mixture of the two diastereomers silybin A and B^49^. Human and animal studies with silymarin or its main component silybin on liver pathologies such as oxidative or lipotoxic stress-induced alcoholic and non-alcoholic fatty liver disease and steatohepatitis show (pre)clinical efficacy^6,51-57^, whereas studies on xenobiotic-induced liver toxicity produced mixed results^42,44,58^, with only rare cases of side effects^59^. Note that the oral bioavailability of silybin can be substantially boosted by specific formulations, yielding systemic silybin plasma concentrations (C_max_) up to 85 μM in humans^42^. The hepatoprotective activities of silymarin/silybin have been ascribed to antioxidant response inducing, anti-inflammatory^60^, antifibrotic, hepatocyte regeneration-stimulating, and membrane-stabilizing properties^51,52,61^. Several studies have found that administration of silymarin/silybin reduces levels of low-density lipoprotein (LDL), VLDL, cholesterol, and/or TGs, while other studies have not observed substantial changes in the serum lipid profile^62-74^, which is not readily understood but may be related to the dose. Recently, silymarin (but not silybin) has been proposed to decrease lipid accumulation during a high-fat diet by altering the vitamin B12-producing capacity of the gut microbiota^75^. On the other hand, silymarin/silybin has been suggested to induce phospholipid turnover by upregulating choline phosphate cytidylyltransferase^76^. Silymarin/silybin compensated for the decrease of PC and phosphatidylethanolamine (PE) in rat liver upon intoxication^77,78^ and, when given as a silybin- and PC-based food integrator to MASH patients, restored plasma PC and sphingomyelin (SM) levels^62^. Whether silymarin/silybin actively promotes phospholipid enrichment or indirectly increases phospholipid levels by alleviating disease conditions is insufficiently understood, as are the consequences for other membrane phospholipid classes and the knowledge of phospholipidomic profiles. The latter is of great importance because imbalances in the membrane phospholipid composition can cause severe alterations in membrane architecture and function^79^.

**Figure 1.**
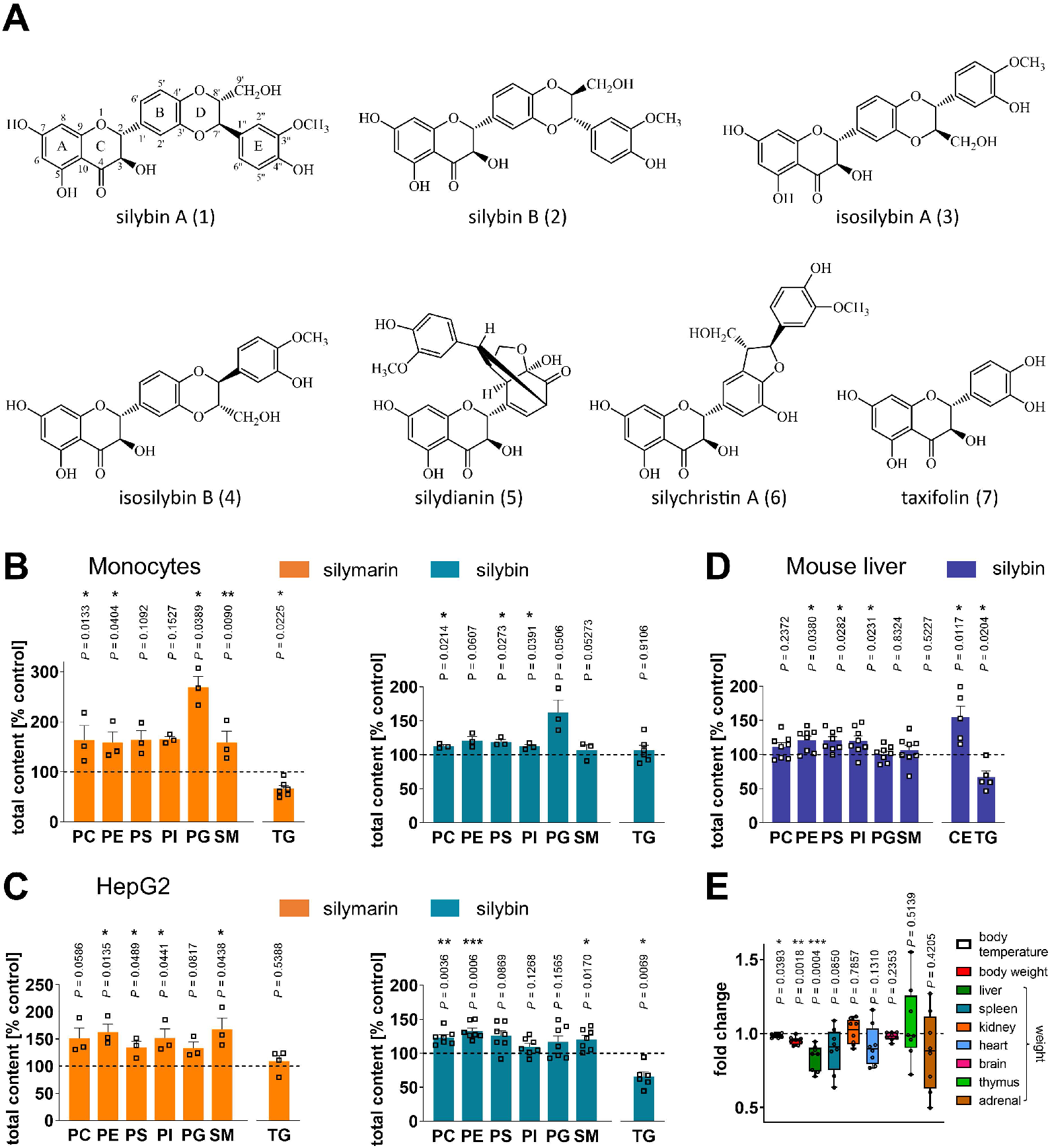
Shift from TGs to phospholipids in human monocytes, hepatocytes, and mouse liver. (A) Main components of silymarin. (B-D) Total amounts of lipid classes were determined by UPLC-MS/MS. (B, C) Human primary monocytes (B) and HepG2 cells (C) were treated with silymarin (50 μg/ml for monocytes and 10 μg/ml for HepG2 cells), silybin (20 μM), or vehicle (ethanol for silymarin, DMSO for silybin) for 24 h. Individual values and mean + SEM; n = 3 (B: except TG, C: silymarin except TG), n = 4 (C: TG silymarin), n = 6 (B: TG silymarin and silybin, C: TG silybin), n = 7 (C: silybin except TG). (D, E) Mice received silybin hemisuccinate (‘silybin’; 200 mg/kg, i.p.) or vehicle (0.9% NaCl) trice at 0, 12, and 24 h and were sacrificed after 37 h. (E) Body temperature, body weight and organ weight of mice upon administration of silybin. Temperature and body weight were measured after 37 h before animals were sacrificed and organs collected. The box-and-whisker plot shows fold-changes upon silybin gavage. The median fold change belonging to each group is shown as bold line. The boxes extend from the 25^th^ to 75^th^ percentiles, and whiskers extend to minimal and maximal values. Individual values and mean + SEM (D) or box plots and individual values (E) from n = 5 (D: CE and TG), n = 7 (body temperature), n = 8 (D: except CE and TG, E: body and organ weights) mice/group. \**P* < 0.05, \*\**P* < 0.01, \*\*\**P* < 0.001 vs. vehicle control. Two-tailed paired (B, C) or unpaired (D, E) Student’s *t*-test.

Here, we demonstrate that silymarin/silybin increases the levels of phospholipids by suppressing their degradation. This effect is partially combined with the induction of phospholipid biosynthetic enzymes, depending on the condition. Simultaneously, it reduces triglycerides by downregulating multiple biosynthetic enzymes in hepatocytes. To some extent, this effect is also observed in extrahepatic cell types. We ascribe this activity to specific structural features of silybin A and find that they prevail in healthy or pre-disease states not yet afflicted with massive lipid overload, whereas TG-lowering mechanisms predominate under the latter severe liver disease conditions. The channeling of fatty acids from triglycerides to phospholipids has the advantage of i) reducing hepatic TG levels and lipid droplet numbers ii) avoiding high lipotoxic levels of free fatty acids are avoided, and iii) expanding intracellular membranes, which may explain the enhanced hepatic biotransformation capacity upon treatment with silybin. Major adverse changes in membrane function are not expected from the balanced upregulation of phospholipid species. Conclusively, our data suggest that the mechanism of silymarin/silybin described here is more effective in protecting against metabolic liver disease rather than reversing advanced disease states

## 2. Results

### 2.1 Silybin induces a switch from hepatic TGs to phospholipids

To investigate the effects of silymarin and silybin on the hepatic lipid composition, we used human HepG2 hepatocarcinoma cells and human primary monocytes (as a surrogate for hepatic phagocytes) and analyzed major phospholipids and neutral lipids by targeted lipidomics. Silymarin (50 μg/ml for monocytes and 10 μg/ml for HepG2 cells) increased the cellular content of major phospholipid classes, i.e., PC, PE, phosphatidylserine (PS), phosphatidylinositol (PI), phosphatidylglycerol (PG), and SM (Figure 1B and C). Similar effects were observed for silybin (20 μM, Figure 1B and C), one of the major bioactive components of silymarin^49^. Instead, TG levels were substantially decreased by both silymarin and silybin treatment, with opposite efficacy in monocytes and hepatocytes. While silymarin specifically reduced TG levels in monocytes, silybin was only effective in hepatocytes (Figure 1B and C). Phospholipid accumulation in HepG2 cells was manifested at ≥ 10 μg/ml silymarin or 20 μM silybin after 24 h (Figure S1), and cytotoxic activities first became evident at ≥ 50-200 μg/ml silymarin and ≥ 100 μM silybin (Figure S2). Silymarin/silybin did not markedly reduce cell proliferation (Figure S2A), membrane integrity (Figure S2B), or mitochondrial dehydrogenase activity in HepG2 cells (Figure S2C) under these experimental conditions. Together, our results suggest that silybin induces a hepatic switch from TGs to phospholipids and point to additional components contained in silymarin that tune the cellular lipid profile.

To investigate whether the decrease in TGs is functionally related to the accumulation of phospholipids, we studied the impact of TG degradation on the cellular PE content, which was robustly upregulated by silybin treatment (Figure 1C). The selective diacylglycerol-*O*-acyltransferase (DGAT)2 inhibitor PF-06424439 (10 μM), which interferes with the final step of TG biosynthesis^80^, decreased TG levels as expected, but failed to increase the amount of PE (Figure S3). Accordingly, inhibition of adipocyte triglyceride lipase (ATGL) using atglistatin neither decreased TG nor significantly elevated PE levels (Figure S3). Thus, our data suggest that TG depletion does not drive phospholipid enrichment, at least under conditions where phospholipid biosynthesis is not upregulated.

Next, we investigated whether silybin counter-regulates phospholipid and TG levels *in vivo*. Mice received silybin (200 mg/kg, i.p.) three times over 37 h, which is expected to produce peak hepatic concentrations >10 nmol/g for the unconjugated drug^81,82^. Silybin increased the hepatic phospholipid content, reaching significance for PE, PS, and PI, and simultaneously lowered TG levels (Figure 1D), as expected from the results for hepatocytes *in vitro*. The shift from TGs to phospholipids was accompanied by a significant loss of liver and body weight (Figure 1E) and a decrease of blood glucose levels (Figure S4), which is of particular interest because fatty liver disease is often associated with insulin resistance that elevates blood glucose levels^3^.

The majority of phospholipids significantly upregulated by silybin in mouse liver contain polyunsaturated fatty acids, either linoleic acid (18:2), arachidonic acid (20:4), or docosahexaenoic acid (22:6) (Figure 2A). Note that an increase in membrane unsaturation has been associated with insulin sensitivity^83^ and may explain the decrease in blood glucose levels with silybin administration (Figure S4). The effect of silymarin/silybin on individual lipid species varies greatly between experimental systems (Figure S5 and S6). While the levels of a broad spectrum of phospholipid species are The flavonolignan silybin favorably redistributes lipids 7 increased, there are also lipids that are regulated in the opposite direction, particularly in mouse liver, where silybin reduces the amount of PC(18:1/18:1) and PE(18:1/18:1), along with other lipids (Figure 2A and Figure S6). The differences between silymarin and silybin lie in the magnitude rather than the direction of the phospholipidomic changes (Figure 2B). In contrast, the levels of TG species are consistently decreased by silymarin in monocytes and by silybin in HepG2 cells (Figure S5).

**Figure 2.**
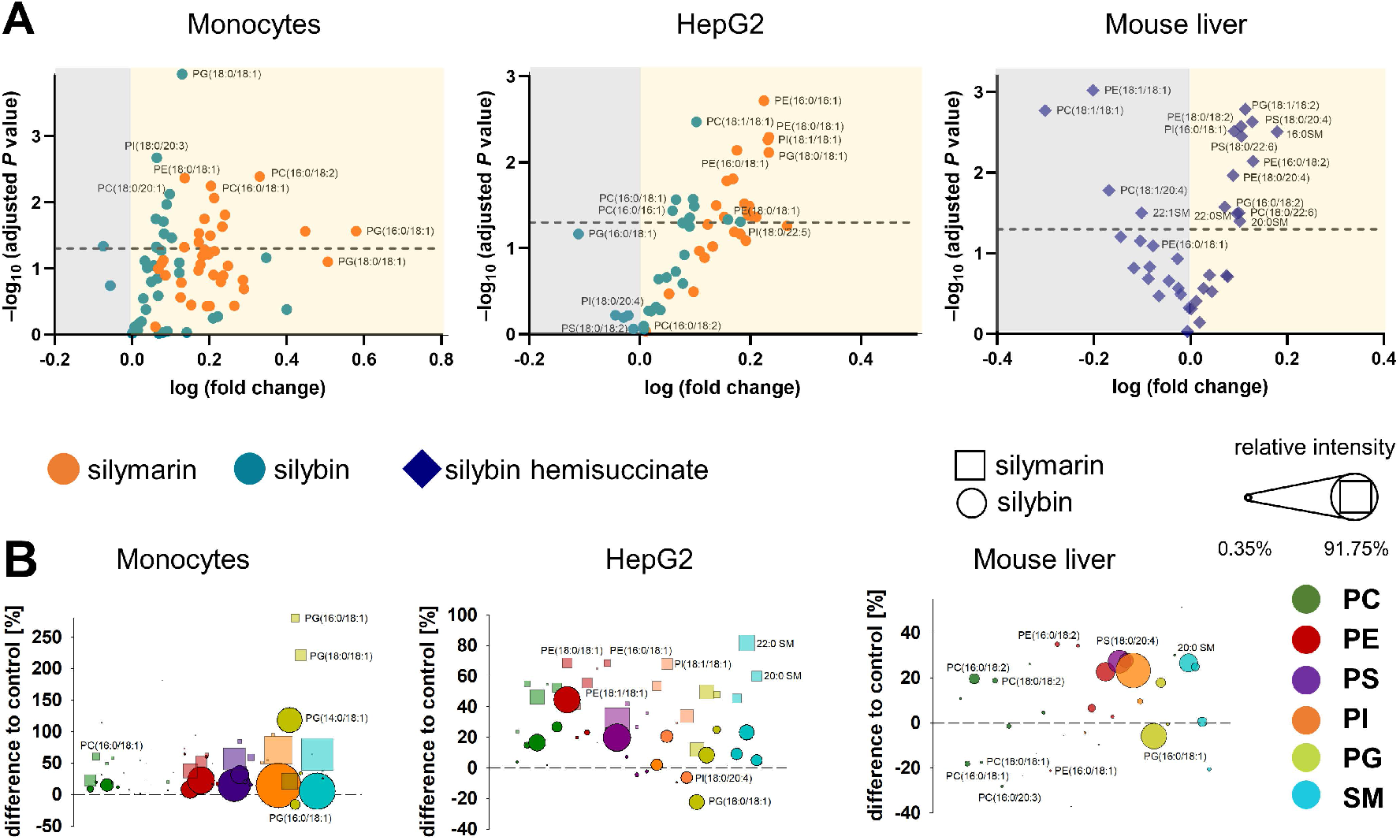
Phospholipid profiling indicates an upregulation of diverse species. Human primary monocytes and HepG2 cells were treated with silymarin (50 μg/ml for monocytes and 10 μg/ml for HepG2 cells), silybin (20 μM) or vehicle (ethanol for silymarin, DMSO for silybin) for 24 h. Mice received silybin hemisuccinate (‘silybin’; 200 mg/kg) or vehicle (0.9% NaCl) trice at 0, 12, and 24 h and were sacrificed after 37 h. (A) Volcano plots showing the cellular proportion of phospholipid species that increase (yellow background) or decrease (grey background) upon treatment with silymarin or silybin. Adjusted P values given vs. vehicle control. The dashed line indicates a P-value of 0.05. (B) Forest plots depicting phospholipid species that are up-(positive values) or down-regulated (negative values) by silymarin (squares) or silybin (circles). Values, calculated as percentage of control, show the difference to 100%, with the dashed line at 0% indicating no difference to control. The dot size describes the mean relative abundance of phospholipid species within the phospholipid subclass (relative intensities). Data and the number of experiments are identical to Figure 1.

To exclude that lipids present in silymarin contribute to changes in the cellular lipid profile, we analyzed the lipid composition of silymarin. Phospholipids with a glycerol backbone (glycerophospholipids) other than PC(16:0/18:2) were not detected in silymarin, and only low-abundance lysophospholipid and SM species were present (Figure S7). Together, the lipids in silymarin do not explain the increase in cellular phospholipids upon treatment.

### 2.2 Accumulated phospholipids are distributed across intracellular membranes

Phospholipids are organized in plasma and intracellular membranes and, to a lesser extent, in lipid droplets and the cytosol^14,84^. It can be excluded that the silymarin/silybin-induced increase in cellular phospholipids is related to the plasma membrane, as the diameter of both monocytes and HepG2 cells was not altered by treatment (Figure 3A). To define the membrane compartment where the additional phospholipids are deposited, we assessed their size and morphology using organelle-specific protein markers. We expected that the 1.2-to 1.5-fold increase in total intracellular phospholipids would be visible as a gain in size or morphological change if the additional phospholipids were preferentially incorporated into a specific membrane compartment. If, instead, the phospholipids are evenly distributed throughout the intracellular membranes, even the 1.5-fold increase in spherical surface area (formed by membrane phospholipids) would result in only a 1.2-fold increase in diameter, and this factor is further reduced for tubular systems such as ER and Golgi with strongly increased surface areas as compared to spherical structures. Apparently visible effects on organelle size and structure are unlikely to be achieved in this case. We focused on large intracellular membrane compartments, i.e., nucleus, ER, and Golgi, which were stained with DAPI and specific antibodies against the integral ER protein calnexin and the *cis*-Golgi marker GM130, respectively. Silymarin/silybin did not markedly affect the intensity or distribution of the immunofluorescence signal (Figure 3B and C), as suggested by visual inspection and confirmed by quantitative analysis of the fluorescence signal. Thus, phospholipids seem to be enriched at intracellular sites but not preferentially incorporated into a major membrane compartment such as the ER, Golgi, or nucleus.

**Figure 3.**
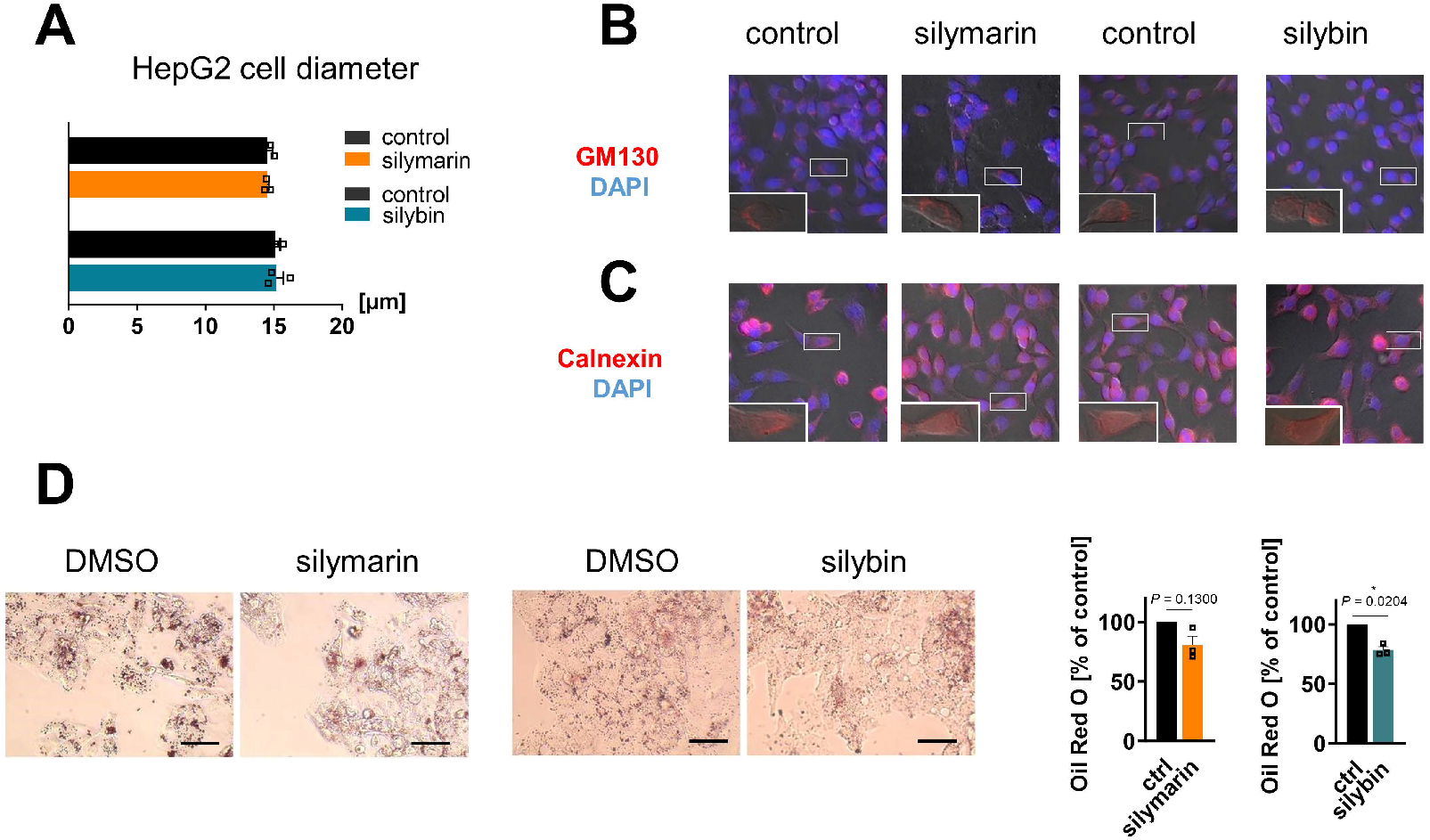
Impact on intracellular membrane compartments and lipid droplets. HepG2 cells were incubated with silymarin (10 μg/ml), silybin (20 μM) or vehicle control (ethanol for silymarin, DMSO for silybin) for 24 h. (A) Cell diameter (μm) of HepG2 cells determined with a Vi-CellTM XR cell counting system. Individual values and mean + SEM; n = 3. (B and C) Immunofluorescence labeling of Golgi (B) and ER (C): anti-GM130 (red) was used as Golgi marker and anti-calnexin (red) as ER marker; nuclei were stained with DAPI (blue). (D) Left panel: Staining of lipid droplets with Oil Red O. Scale bar 50 μM. Right panel: Relative lipid droplet content. Individual values and mean + SEM; n = 3. *P < 0.05 vs. vehicle control. Two-tailed paired Student’s t-test (A, D).

### 2.3 Silybin causes a decrease in lipid droplets

Lipid droplets are universal storage organelles for neutral lipids such as TG and cholesteryl esters (CE) and represent dynamic cellular organelles with an important role in lipid and membrane homeostasis^14^. We treated HepG2 cells with silymarin or silybin and stained lipid droplets with either Oil Red O or 4,4-difluoro-1,3,5,7,8-pentamethyl-4-bora-3a,4a-diaza-*s*-indacene (BODIPY 493/503). Lipid droplet content (spectroscopically quantified based on incorporated Oil Red O) (Figure 3D) and lipid droplet number (assessed from images of BODIPY493/503-stained cells) (Figure S8) were decreased by silybin, as expected from the declining TG levels (Figure 1C) and in accordance with previous *in vitro* and *in vivo* studies using silymarin or silybin^85-89^. Silymarin was considerably less efficient in reducing both TG levels (Figure 1C) and lipid droplet content and number in HepG2 cells (Figure 3D and Figure S8).

### 2.4 Stereochemical requirements of silybin for targeting lipid metabolism

Natural silybin is a mixture of the diastereoisomers silybin A and B^49^. To elucidate the active isomer and explore crucial structural features, we applied an efficient preparative HPLC method to obtain the two isomers A and B in pure form^90^. Starting from these isomers, the corresponding 2,3-dehydrosilybin enantiomers and the hemiacetal product, in which the 2,3-dihydro-chromane is replaced by 2*H*-benzofuran-3-one, were synthesized^91^ (Figure 4). Lipidomic analysis revealed that silybin A increased phospholipid and decreased TG levels in HepG2 cells, whereas silybin B was considerably less effective (Figure 4). Introduction of a double bond into the flavanon-3-ol moiety of silybin yielded 2,3-dehydrosilybin, which (as *7’R,8’R* isomer A) decreased TG levels comparably to silybin but was no longer active on phospholipids (Figure 4). These findings indicate that both, the 2,3-dihydrochromane and the 1,4-benzodioxan scaffold of silybin A contribute to the phospholipid-accumulating activity, whereas modifications of the 2,3-dihydrochromane ring are compatible with TG-lowering properties. Hence, silybin seems to modulate TG and phospholipid metabolism through independent mechanisms.

**Figure 4.**
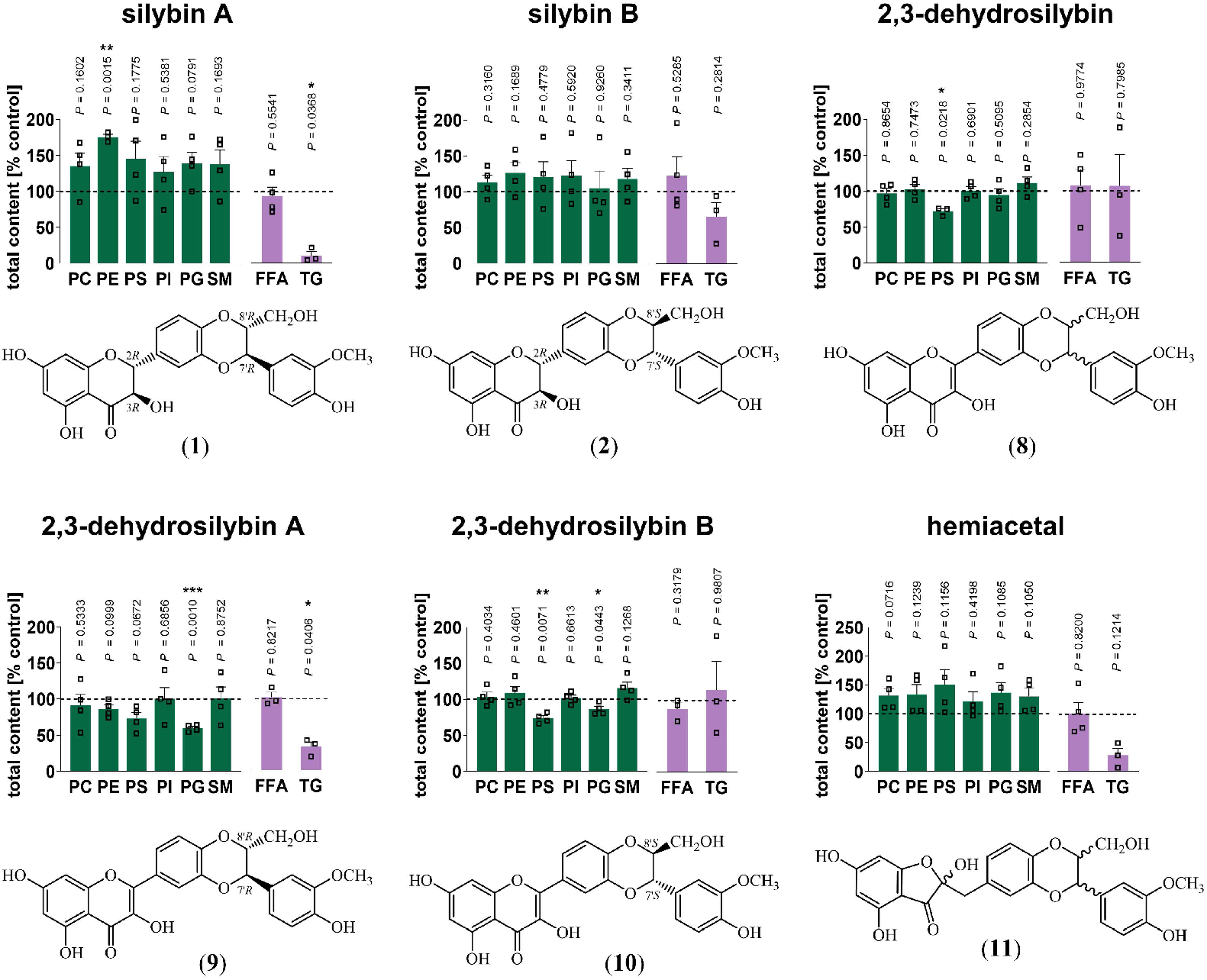
Silybin A is the active isomer that causes the switch from TGs to phospholipids. HepG2 cells were treated with the indicated compounds (20 μM) or vehicle (DMSO) for 24 h. Total amounts of lipid classes were determined by UPLC-MS/MS. Individual values and mean + SEM; n = 3 (PE silybin A, PS dehydrosilybin, TGs, free fatty acids (FFA) 2,3-dehydrosilybin A and B) n = 4 (except PE silybin A, PS dehydrosilybin, TGs, free fatty acids FFA 2,3-dehydrosilybin A and B). *P < 0.05, **P < 0.01, ***P < 0.001 vs. vehicle control (DMSO). Two-tailed paired Student’s t-test.

2,3-Dehydrosilybin and its isomers A and B (Figure 4) selectively decreased the abundance of anionic phospholipids. On the one hand, 2,3-dehydrosilybin lowered the cellular PS content, which we ascribed to isomer B. On the other hand, both isomers, but surprisingly not the stereomeric mixture, induced a drop of PG (2,3-dehydrosilybin A > 2,3-dehydrosilybin B), the precursor of cardiolipins^79^. The hemiacetal (Figure 4) increased phospholipid and decreased TG levels by trend, being slightly less efficient than silybin A (Figure 4) but more active than the stereomeric mixture of silybin (Figure 1C). Neither cell number nor membrane integrity were substantially reduced by any of the silybin derivatives up to 20 μM (Figure S9). Together, the effects of silybin on the cellular lipid profile are mediated by only one isomer, and small changes in its structure allow to dissect the activities on phospholipids and TGs.

### 2.5 Silybin-induced phospholipid accumulation does not rely on G2/M cell cycle arrest or ER stress

Besides being hepatoprotective, silymarin and silybin exhibit anticancer activity *in vitro*^47^ and upon oral gavage *in vivo*^92^. The decrease in cancer cell viability and induction of apoptosis by silybin (≥ 50 μM) has been attributed to, among others, disruption of Ca^2+^-homeostasis and induction of ER stress^93,94^. We asked whether silymarin/silybin causes a mild G2/M arrest, which could prevent cell division, or induces the unfolded protein response (UPR), an adaptive stress pathway aimed at augmenting the capacity of the ER, for example by incorporating proteins, such as biotransformation enzymes^95^. Both processes would increase phospholipid levels along with either cell or ER size, which we did not observe for either silymarin or silybin under our experimental conditions (Figure 3A and C). To further exclude a major role of G2/M cell cycle arrest in silymarin/silybin-induced phospholipid accumulation, we stained the DNA with propidium iodide and monitored cell cycle progression by flow cytometry. Neither silymarin (10 μg/ml) nor silybin (20 μM) induced a G2/M cell cycle arrest in HepG2 cells (Figure 5A). Silymarin slightly increased the fraction of cells in sub-G1 and S phase and silybin tended to increase the fraction in sub-G1 and G1 phase (Figure 5A). The anti-microtubule agent vinblastine, used as control, instead efficiently blocked the G2/M transition, as expected. Together, silymarin and silybin do not induce substantial cell cycle arrest at concentrations that are effective on lipid metabolism.

**Figure 5.**
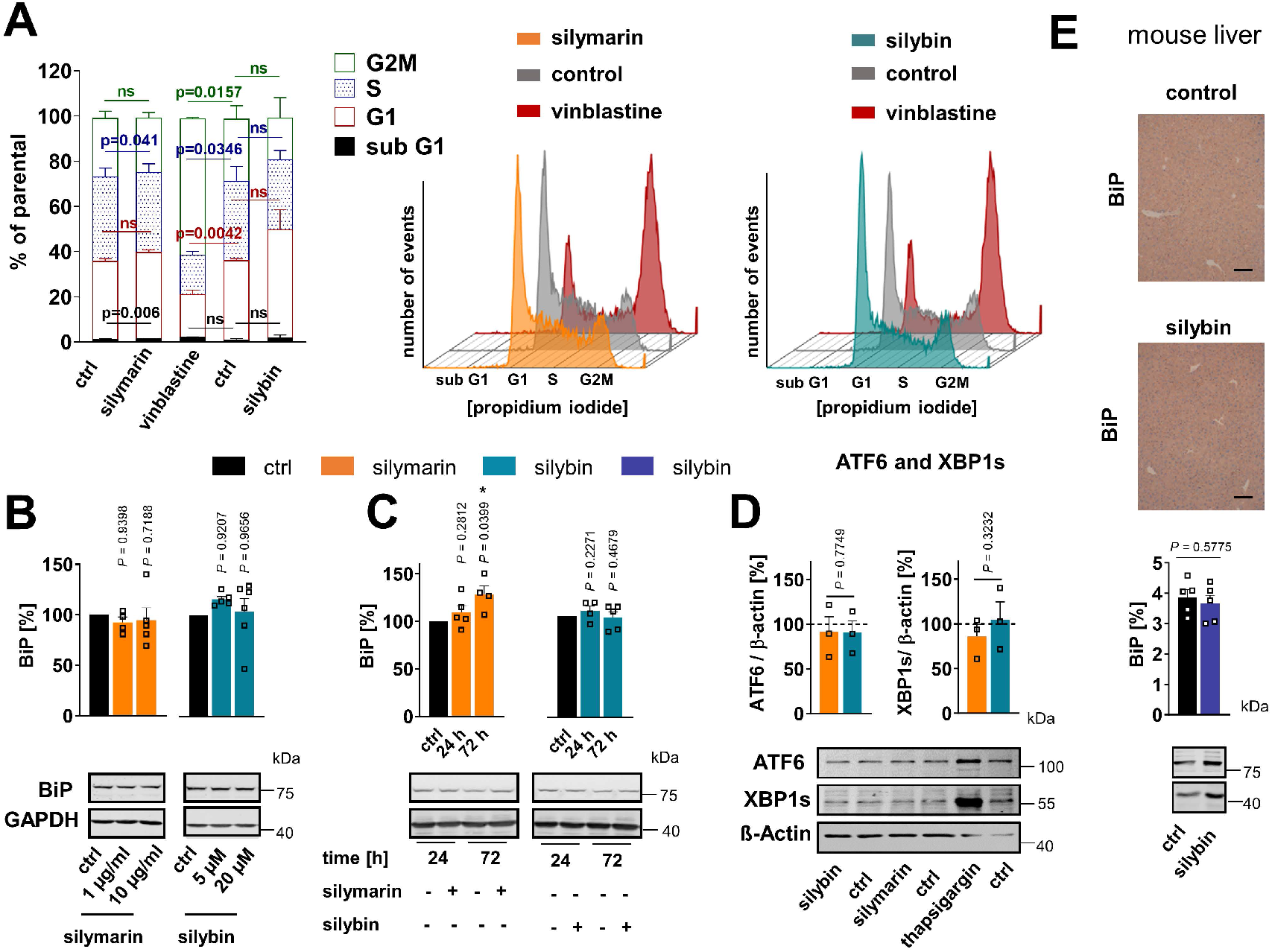
Cell cycle arrest and ER stress are limited to treatment with silymarin. (A-C) HepG2 cells were incubated with silymarin (10 μg/ml), silybin (20 μM), vinblastine (100 nM), or vehicle (DMSO for silybin and vinblastine, ethanol for silymarin) for 24 h unless indicated otherwise. (A) Propidium iodide staining and cell cycle analysis by flow cytometry. The relative distribution of cells in subG1, G1, S and G2/M phase is shown as mean + SEM; n = 3. Histograms are representative of three independent experiments. (B, C) Concentration-(B) and time-dependent (C) expression of the ER stress marker BiP. Individual values and mean + SEM; n = 4 (B: 1 μg/ml silymarin, C: silymarin 72 h, silybin 24 h), n = 5 (B: 5 μM silybin and 10 μg/ml silymarin, C: silymarin 24 h, silybin 72 h) or n = 6 (B: 20 μM silybin). (D) Expression levels of the ER stress marker ATF6 and XBP1s. Individual values and mean + SEM; n = 3. (E) Mice received silybin hemisuccinate (‘silybin’; 200 mg/kg, i.p.) or vehicle (0.9% NaCl) trice at 0, 12, and 24 h and were sacrificed after 37 h. Upper panel: Immunohistochemical analysis of BiP in mouse liver; scale bar, 100 μm. Lower panel: BiP protein expression in mouse liver homogenates determined by Western blotting. Individual values and mean + SEM; n = 5 mice/group. *P < 0.05 vs. vehicle controls. Repeated measures one-way ANOVA + Tukey HSD post hoc tests (A-C) or two-tailed paired (D) or unpaired (E) Student’s t-test.

To assess the effect of silymarin and silybin on ER stress and UPR induction, we analyzed the expression of the marker protein BiP (GRP78), which is an ER chaperone and a major regulator of ER stress signaling leading to UPR activation^96^. Silymarin and silybin only marginally enhanced BiP expression in HepG2 cells within 24 h (Figure 5B and C), and comparable results were obtained for two additional markers of ER stress, spliced X-box binding protein 1 (XBP1s) and activating transcription factor (ATF)6 (Figure 5D). Conversely, the expression of these markers was significantly induced by thapsigargin, a known elicitor of ER stress (Figure 5D). Prolonged exposure of HepG2 cells (72 h) to silymarin slightly increased BiP levels, but this did not seem to be related to silybin (Figure 5C). Accordingly, silybin (200 mg/kg) failed to induce BiP expression in mouse liver upon i.p. administration, as determined by immunoblotting and immunohistochemistry (Figure 5E). In conclusion, silymarin and silybin do not significantly impact ER stress under experimental conditions that effectively increase phospholipid levels.

### 2.6 Targeting lipid metabolism

To elucidate the molecular mechanisms by which silymarin/silybin induces a lipid class switch from TGs to phospholipids, we reanalyzed previously published transcriptomic datasets from hepatocytes (*in vitro* and *in vivo*) and acquired the transcriptome of an exemplary extrahepatic cell line. We focused on genes from the category “Lipid Metabolism” of the Reactome Pathway Database^97^ and studied their expression in four experimental systems *in vitro* and *in vivo*: i) human HepG2 hepatocarcinoma cells treated with silymarin (12 μg/ml) for 24 h^98^, ii) human Huh7.5.1 hepatocarcinoma cells treated with silymarin (40 μg/ml) for 4, 8, and 24 h^99^, human Caco-2 colon carcinoma cells treated with either silybin (30 μM) or silymarin (30 μg/ml) for 24 h, and iv) hepatocytes isolated from chronically hepatitis C virus (HCV)-infected mice receiving daily intravenous injections of silybin (265-469 mg/kg) for 3 or 14 d^100^. Silybin/silymarin affects the expression of a wide range of enzymes and factors involved in lipid metabolism, but the effects are moderate and, with one exception, do not reach significance after global correction for false discovery (Figure S10A-D). Only the cytochrome P_450_ (CYP) monooxygenase *CYP1A1*, which accepts various endogenous substrates, including steroids and polyunsaturated fatty acids^101-105^, is highly significantly upregulated in Caco-2 cells (Figure S10C).

#### 2.6.1 Phospholipases and phospholipid biosynthesis

Given the detected changes in the HepG2 lipidome (Figure 1B-D), we extended our study to genes that were differentially regulated according to non-adjusted *P*-values and for which the respective pathway was significantly regulated in the same direction for at least two independent model systems. We found that silybin/silymarin i) decreased the expression of several hydrolases involved in phospholipid degradation (Figure 6A-F), including phospholipases A1 (*PLA1A*, Figure 6B), phospholipases A_2_ (*PLA2G1B, PLA2G6*, Figure 6A and D, and Figure S11), and phospholipase D (*PLD1, PLD6*, Figure 6A, B, and D, and Figure 11), specifically in primary hepatocytes and hepatocyte-derived cell lines. In addition, silybin/silymarin ii) upregulates factors that deplete phospholipases (*PLA2R1*, Figure 6B), iii) downregulates enzymes that degrade intermediates in phospholipid biosynthesis (*TECR, MGLL, ACP6, GDPD3, PNPLA7*, Figure 6B, D and E), and iv) less consistently induces the expression of phospholipid biosynthetic enzymes and other factors (*GNPAT, CHKA, SLC44A1, AGPS, AGPAT2, MBOAT2, LPGAT1, DEGS1, CERS6*, Figure 6C and D and Figure S11). Compensatory mechanisms seem to exist that decrease phospholipid biosynthesis (*via PCYT1A, ETNK2, PEMT, GPAM, SPTLC3, CERS2*, Figure 6B, C, D and E) or enhance phospholipid degradation (*PLA2G4C, DDHD1, ACER3, PLD6*, Figure 6B, C and D), possibly buffering the accumulation of phospholipids or rearranging phospholipid profiles through different substrate specificities.

**Figure 6.**
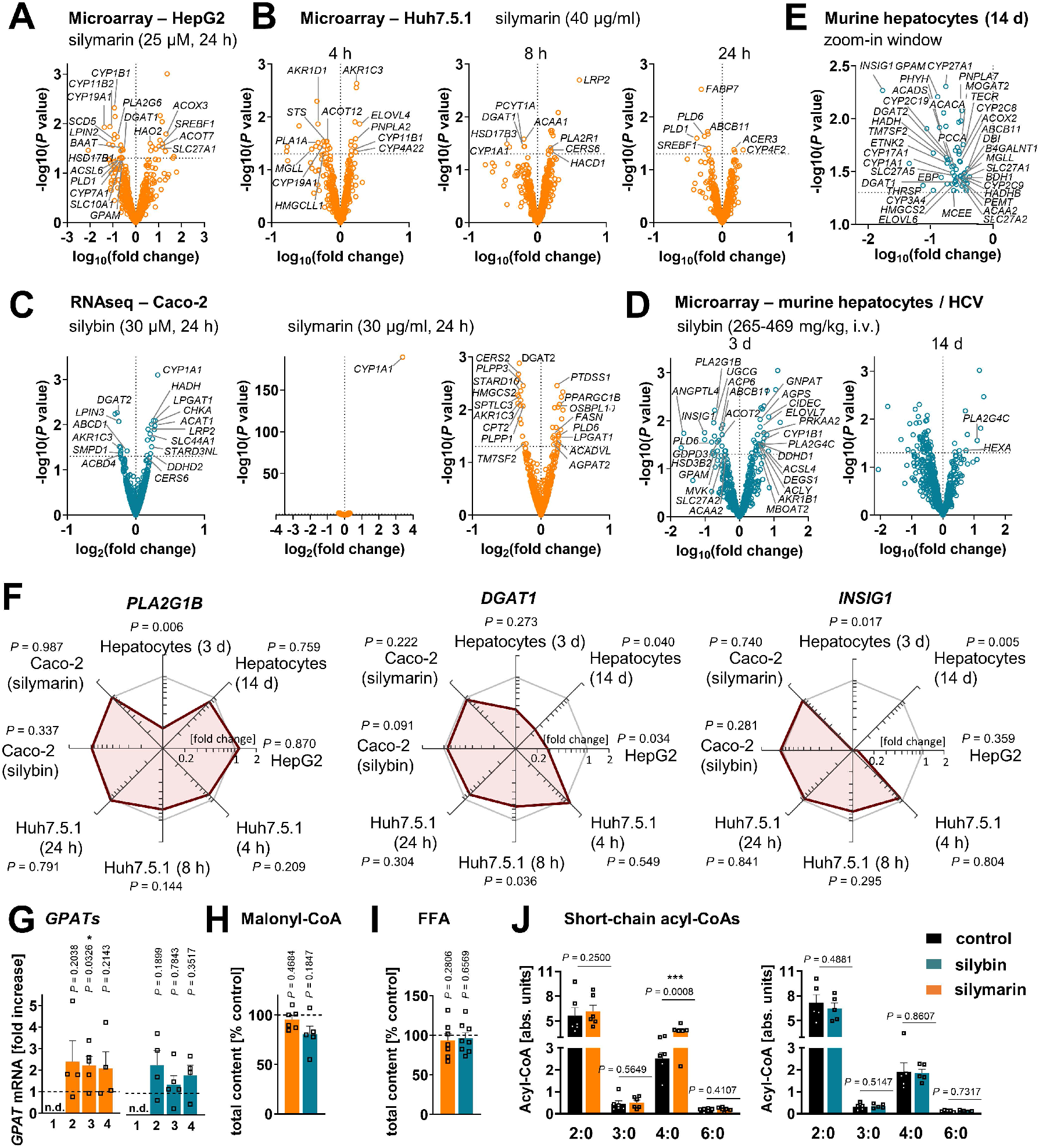
Silymarin/silybin induces global changes in phospholipid, TG and sterol metabolism. (A-F) Comparative analysis of transcriptome data from silymarin-treated HepG2 (A) and Huh7.5.1 hepatocarcinoma cells (B), silymarin- and silybin-treated Caco-2 colon carcinoma cells (C), and hepatocytes derived from HCV-infected mice receiving silybin (D, E). Volcano plots compare the expression of lipid metabolic genes upon silymarin (A-C) or silybin (C-E) treatment vs. vehicle control. Differentially expressed genes are defined as those that show consistent regulation in the same direction in at least two independent model systems at a significance level of P < 0.05 (without adjustment for multiple comparisons) and are annotated in the corresponding plots. The dashed line indicates a P-value of 0.05; multiple two-tailed unpaired Student’s t-tests. (F) Radar plots indicating the fold change in PLA2G1, DGAT1, and INSIG1 expression by silymarin (HepG2, Huh7.5.1, Caco-2) or silybin (hepatocytes, Caco-2) relative to vehicle control. Non-adjusted P values given vs. vehicle control; multiple two-tailed unpaired Student’s t-tests (G-J). HepG2 cells were incubated with silymarin (10 μg/ml), silybin (20 μM) or vehicle (ethanol for silymarin, DMSO for silybin) for 24 h. (G) mRNA levels of GPAT2-4 normalized to β-actin. Individual values and mean + SEM as fold-change of control; n = 4 (GPAT2 and GPAT4), n = 5 (GPAT3). (H) Cellular levels of malonyl-CoA. Individual values and mean + SEM, n = 5 (silybin) n = 6 (silymarin). (I) Cellular levels of free fatty acids (FFA). Individual values and mean + SEM; n = 7 (silymarin) or n = 8 (silybin). (J) Effects of silymarin and silybin on the cellular ratio of short-chain acyl-CoAs, normalized to the internal standard [13C3]-malonyl-CoA. Individual values and mean + SEM; n = 5 (silybin) and n = 6 (silymarin). *P < 0.05, ***P < 0.001 vs. vehicle controls; two-tailed paired Student’s t-tests.

To investigate whether silymarin/silybin elevates phospholipid levels via de novo phospholipid biosynthesis under our experimental conditions, we treated HepG2 cells with silymarin or silybin for 24 h and determined the mRNA expression of glycerophosphate acyltransferase (GPAT) isoenzymes and lysophosphatidic acid acyltransferase (LPAAT)/lysophospholipid acyl-transferase (LPLAT) isoenzymes at the mRNA level. GPATs and LPAATs successively transfer acyl-chains from acyl-CoA to the sn-1 and sn-2 positions of glycerol-3-phosphate to form phosphatidic acid, the common precursor of glycerophospholipids and TGs^79^. Silymarin and silybin increased the mRNA levels of GPAT isoenzymes 2 to 4, reaching significance for the silymarin-mediated induction of GPAT3 (Figure 6G), which is consistent with a previous report showing enhanced Gpat3 mRNA expression in the liver of silybin-treated mice on a methionine- and choline-deficient diet85. In contrast, the expression of LPAAT/LPLAT isoenzymes was not markedly affected (Figure S12A). Together, the moderate but versatile induction of phospholipid biosynthesis and inhibition of phospholipid degradation by silymarin/silybin likely accounts for the accumulation of phospholipids in hepatocytes.

#### 2.6.2 TG metabolism

The decrease in TG levels is driven by the repression of genes associated with the generation of DAGs from either phosphatidate (*LPIN2, LPIN3, PLPP1, PLPP3*, Figure 6A and C, and Figure S11) or monoacylglycerols (*MOGAT2*, Figure 6E) and their acylation to TGs (*DGAT1, DGAT2*, Figure 6A, B, C and E, and Figure S11), as suggested by comparative transcriptomics. The concrete mode of action seems to be context-dependent and possibly under kinetic control, as suggested by the failure of silybin and silymarin to reduce *DGAT1* and *DGAT2* protein expression in HepG2 cells 24 h after treatment (Figure S12B). TGs are a major component of the hydrophobic core of lipid droplets, which form contact sites with essentially all other cellular organelles and are at the nexus of lipid and energy metabolism^14,106,107^. Interestingly, selective inhibition of DGAT1 (by A-922500) or DGAT2 (by PF-06424439) and antagonism of the DGAT-inducing transcription factor peroxisome proliferator activated receptor (PPARγ)^108^ (by GW9662) moderately reduced lipid droplet staining in palmitate (PA, 16:0)-loaded human HepaRG hepatocytes (Figure 7A), but only the combined inhibition of DGAT1 and DGAT2 reached the efficacy of the silybin isomer A (Figure 7B). Since lipolysis of TGs in lipid droplets is initiated by ATGL/PNPLA2^109^, we investigated the effect of silymarin/silybin on the protein expression of this enzyme, but again found no substantial regulation (Figure S12B), consistent with the transcriptomics data (Figure 6A-E). Note that selective inhibition of ATGL (by atglistatin) also failed to increase lipid droplet signals in stressed HepaRG cells (Figure 7A).

**Figure 7.**
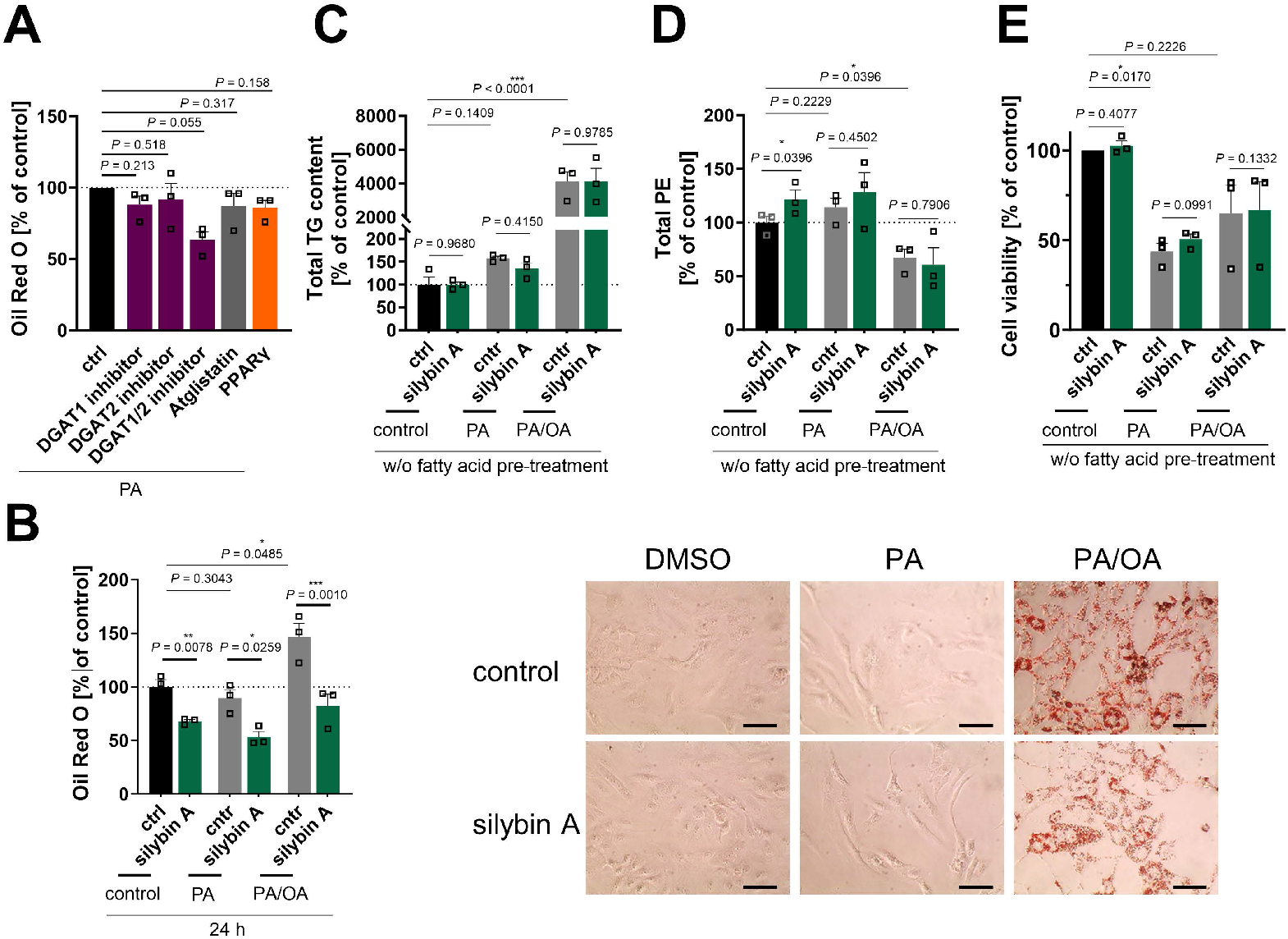
The efficacy of silybin in inducing a lipid class switch differs between hepatocyte pre-disease and disease models. (A,B) HepaRG cells were treated with 0.1 mM palmitate (PA) or a mixture of PA/oleate (OA) in a 1:2 ratio (in total 1 mM) together with vehicle (DMSO, 0.5%), silybin A (20 μM), the ATGL inhibitor atglistatin (50 μM), the DGAT1 inhibitor A 922500 (5 μM), the DGAT2 inhibitor PF-06424439 (10 μM), a combination of DGAT1 (5 μM) and DGAT2 inhibitors (10 μM), or the PPARγ antagonist GW9662 (5 μM) for 24 h. (A) Relative lipid droplet content. Individual values and mean + SEM, n = 3. (B) Left panel: Relative lipid droplet content. Individual values and mean + SEM, n = 3. Right panel: Representative images of HepaRG cells stained for lipid droplets using Oil Red O; scale bar, 50 μm. (C, D) HepaRG cells were co-treated directly with with 0.1 mM palmitate (PA) or a mixture of PA/oleate (OA) in a 1:2 ratio (in total 1 mM) and vehicle (DMSO, 0.5%) or silybin A (20 μM) for 24 h. Total levels of TG (C) and PE (D) determined by UPLC-MS/MS. Individual values and mean + SEM, n = 3. (E) Cell viability measured by MTT assay. Individual values and mean + SEM, n = 3. *P < 0.05, **P < 0.01, ***P < 0.001 vs. control; two-tailed paired (A, B, E) or unpaired (C, D) Student’s t-test.

#### 2.6.3 Fatty acid biosynthesis

Both phospholipid and TG biosynthesis depend on the availability of activated fatty acids^110^. Their biosynthesis from acetyl-CoA is an energy- and NADPH-consuming process, which is initiated by the rate-limiting enzyme acetyl-CoA carboxylase (ACC, *ACACA*)^111^. The product of this reaction, malonyl-CoA, is subsequently transferred to fatty acid synthase (*FASN*), which produces long-chain fatty acids that are activated as CoA esters by acyl-CoA synthetases before further metabolism^112,113^. As expected from the multiple roles of acyl-CoAs in lipogenesis, silymarin/silybin ambiguously regulates genes related to fatty acid metabolism, with expression changes either promoting or inhibiting *de novo* fatty acid biogenesis (*ACACA, FASN, SCD5*, Figure 6A, C, and E), fatty acid uptake respectively activation (*SLC27A1, SLC27A2, SLC27A5, ACSL4, ACSL6*, Figure 6A, D, E), fatty acid elongation (*ELOVL4, ELOVL6, ELOVL7, TECR*, Figure 6B, D and E), and the intracellular transport of acyl-CoAs (*ACBD4, DBI, HACD1*, Figure 6B, C and E). In HepG2 cells, silymarin/silybin slightly increased ACC/*ACACA* (but not FASN) protein expression, which was significant for silybin (Figure S12B), while ACC phosphorylation, which inactivates ACC^111^, tend to be decreased (Figure S12B). This weak stimulatory regulation of ACC by silymarin/silybin was not translated into increased cellular concentrations of i) malonyl-CoA (ACC product, Figure 6H), ii) long-chain fatty acids (FASN products, Figure 6I), or iii) long-chain acyl-CoAs (acyl-CoA synthetase products, Figure S12C).

Conclusively, silybin and silymarin induce changes in fatty acid anabolism that may contribute to, but do not appear to be essential for, the lipid class switch from TGs to phospholipids.

#### 2.6.3 Fatty acid degradation

Since the intracellular concentration of long-chain fatty acids is not markedly altered by silymarin/silybin (Figure S12C), while the fatty acid storage capacity in TGs is compromised (Figure 1B-D), we addressed the fate of fatty acids. On the one hand, they seem to be channeled towards phospholipid biosynthesis, as supported by our data (Figure 1B-D). On the other hand, they might be subjected to fatty acid oxidation *via* mitochondrial or peroxisomal pathways to sustain the energy demand for phospholipid biosynthesis^33,114-116^. In support of this hypothesis, oral administration of silybin increased the mRNA expression of carnitine palmitoyl-transferase 1α (*Cpt1a*) in mouse liver, suggesting an efficient transfer of acyl-CoAs into mitochondria for β-oxidation^85^. Transcriptomic analysis underlines that mitochondrial (*HADH, ACAT1, ACADVL*, Figure 6C, Figure S11) and peroxisomal β-oxidation (*ACOX3, HAO2*, Figure 6A) are enhanced for specific settings, and we confirmed in cultured HepG2 cells that silymarin increased the levels of the β-oxidation intermediate butyryl-CoA in cultured hepatocytes (Figure 6J). However, the effect does not seem to be mediated by silybin, which failed to enrich β-oxidation intermediates (Figure 6J). Since extensive fatty acid oxidation depletes fatty acid concentrations and thus competes with efficient phospholipid biosynthesis, we would expect fatty acid degradation to be kept in check. Consistent with these considerations, silymarin/silybin decreased the mitochondrial degradation of straight-chain, odd-chain, and branched fatty acids (*CPT2, ACAA1, ACAA2, HADH, ACADS, HADHB, PCCA, MCEE*, Figure 6B, C, D, E, Figure S11) as well as peroxisomal oxidation (*ABCD1, ACOX2, PHYH*, Figure 6C and E, Figure S11) and ketogenesis (*HMGCS2, BDH1, HMGCLL1*, Figure 6B, C, E), especially in mouse liver *in vivo* and Huh7.5.1 hepatoma cells *in vitro*. Fatty acid oxidation by CYP enzymes is also subject to intense regulation. Among the various CYP enzymes repressed by silymarin/silybin are those involved in the epoxidation and hydroxylation of polyunsaturated fatty acids (*CYP2C8, CYP2C9, CYP2C19, CYP3A4*, Figure 6E, Figure S11). ω-Oxidases are instead upregulated (*CYP4F2, CYP4A22*, Figure 6B and Figure S11), and results for *CYP1A1* are mixed (Figure 6B, C and E). A detailed description of the impact of silymarin/silybin on cholesterol and CE metabolism is given in Supplementary Note 1.

Together, silymarin/silybin induce a lipid class switch from TGs to phospholipids by limiting TG biosynthesis and suppressing phospholipid degradation in both hepatocytes and extrahepatic cells, partly combined with enhanced phospholipid biosynthesis. These central adaptations are accompanied by pronounced changes in cholesterol and fatty acid metabolism.

### 2.7 Efficacy of silybin in in vitro models of MAFLD and lipotoxicity

The predominant fatty acids present in TGs of the liver, both in healthy individuals and in MAFLD patients, are palmitic acid (PA, 16:0) and oleic acid (OA, 18:1)^117^. Following previously published procedures^118^, we established *in vitro* models of MAFLD and acute lipotoxicity by overloading human HepaRG cells (as a surrogate for normal hepatocytes^119^) with a balanced saturated/unsaturated fatty acid mixture (PA:OA = 1:2) or by challenging them with the saturated fatty acid PA^118^. We monitored the (time-dependent) increase in lipid droplets (Figure 7B and Figure S13A), TG levels (Figure 7C and Figure S13B), and phospholipid content, specifically PE (Figure 7D and Figure S13C) and PC levels (Figure S13D), and determined the consequences on cellular dehydrogenase activity (as a measure of cell viability) (Figure 7E and Figure S13E), viable cell number (Figure S13F), and membrane integrity (Figure S13G). PA/OA strongly increased lipid droplet staining (Figure 7B), elevated TG levels (Figure 7C), and caused a shift from PE (Figure 7D) to PC (Figure S13D) within 24 h. PA was less efficient in increasing TG levels and did not enhance the lipid droplet signal (Figure 7B), but raised the levels of both phospholipid subclasses investigated (Figure 7D and Figure S13D), as expected from the associated induction of ER stress and the UPR^120,121^. The effects were less pronounced or even disappeared at longer incubation times (48 h) (Figure S13A-D). While OA/PA did not or hardly impair the metabolic activity of the cells (Figure 7E and Figure S13E), PA was cytotoxic within 24 h (EC_50_ = 70 μM) (Figure 7E, and Figure S13E), but did not yet disrupt membrane integrity (Figure S13G) or reduce the number of viable cells (Figure S13F).

To further validate the experimental model, we first investigated whether silybin A is able to induce a lipid class switch in unchallenged HepaRG cells, as expected from our studies in HepG2 cells (Figure 4). Indeed, silybin A (although the effects were less pronounced) induced a lipid class switch in HepaRG cells, but apparently from neutral lipids in lipid droplets to phospholipids (Figure 7B and Figure S13D), with little effect on total cellular TG levels (Figure 7C). We then investigated the effect of silybin A in the two disease models (PA or PA/OA treatment): silybin A still reduced the lipid droplet (but not TG) content (Figure 7B and C), but became less efficient in upregulating phospholipid levels (Figure 7D, and Figure S13C and D) and did not attenuate the lipotoxic drop in cell viability (Figure 7E). Our data indicate that silybin A preferentially redirects lipid metabolism from TGs to phospholipids in healthy hepatocytes and extrahepatic cells, and that this metabolic switch becomes less efficient under severe lipid overload, which might provide a mechanistic basis for the mixed results in clinical trials both, under disease and non-disease conditions^74,122-126^.

### 2.8 Activation of hepatic phase I and II metabolism in healthy mice

Silybin induces the expression of phase II enzymes (including glutathione S-transferase, GST) in mouse liver and other tissues^82,127-129^. Instead, the consequences on CYP monooxygenases (phase I enzymes) are mixed^82,129-131^, possibly due to superimposed direct enzyme inhibition, differences between healthy and diseased states, and different kinetics^132-136^.

To gain an overview about the global regulation of drug-metabolizing enzymes by silymarin/silybin, we analyzed the transcriptome data from the experimental systems described in section 2.6 for changes in the expression of genes of the Reactome Pathway Database^97^ categories ‘metabolism - oxidation’, ‘phase I metabolism (compound functionalization)’, and ‘phase II metabolism (compound conjugation)’. Hepatocytes from silybin-treated HCV-infected mice showed a clear kinetic trend: genes of drug-metabolizing enzymes are initially upregulated (day 3) and then downregulated with prolonged treatment (day 14) (Fig. 8A). Instead, the mRNA expression of *CYP* enzymes was differentially regulated in cell-based systems, with individual isoenzymes being up- or downregulated (Fig. S14A-C), following independent kinetics (Fig. S14B). With few exceptions (*CYP3A5, CYP26A1*, Figure 8A and B, Figure S14B), silymarin/silybin decreased the mRNA expression of those CYP enzymes that are prominently involved in drug metabolism (*CYP2B6, CYP2C8, CYP2C9, CYP2C19, CYP3A4*, Figure 8A and B, Figure S14B and D) and of amino oxidases (*AOC3, MAOA, MAOB*) (Figure 8A and B, Figure S14A and B), which oxidatively deaminate xenobiotic amines, in cell-based systems and *in vivo* after prolonged administration.

**Figure 8.**
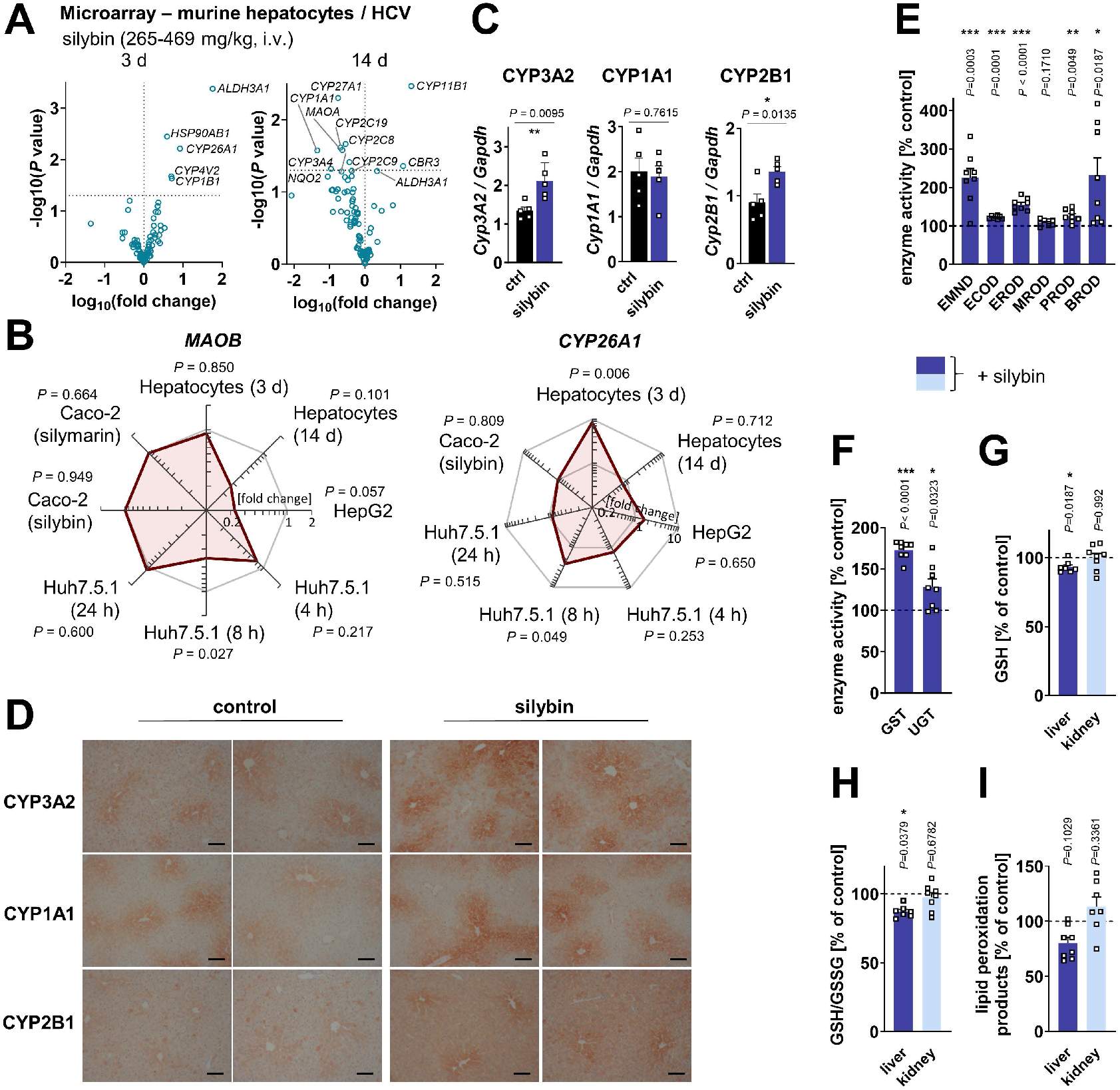
Elevated CYP enzyme expression and activity in mouse liver. (A) Comparative analysis of transcriptome data from hepatocytes derived from silybin-treated HCV-infected mice. Volcano plots compare the expression of lipid metabolic genes upon silybin treatment vs. vehicle control at day 3 and day 14. The dashed line indicates a non-adjusted P-value of 0.05; multiple two-tailed unpaired Student’s t-tests. (B) Radar plots indicating the fold change in MAOB and CYP26A1 expression by silybin relative to vehicle control. Non-adjusted P values given vs. vehicle control; multiple two-tailed unpaired Student’s t-tests. (C-I) Mice received silybin hemisuccinate (‘silybin’; 200 mg/kg, i.p.) or vehicle (0.9% NaCl) trice at 0, 12, and 24 h and were sacrificed after 37 h. (C) Protein expression of CYP3A2, CYP1A1, and CYP2B1 in mouse liver homogenates. Representative Western blots are shown in Figure S15. Individual values and mean + SEM; n = 5 mice/group. (D) Immunohistological analysis of CYP3A2, CYP1A1 and CYP2B1 expression in mouse liver; scale bar, 100 μm. n = 5 mice/group. (E) Cyp activity measured by detecting the oxidative demethylation product formaldehyde (EMND) or the conversion of fluorogenic substrates in mouse liver homogenates. EMND: N-ethylmorphine N-demethylation (CYP3A), ECOD: 7-ethoxycoumarin O-deethylation (CYP1A and CYP2A-C), EROD: 7-ethoxyresorufin O-deethylation (CYP1A), MROD: 7-methoxyresorufin O-demethylation (CYP1A), PROD: 7-pentoxyresorufin O-depentylation (CYP2B), BROD: 7-benzyloxyresorufin O-debenzylation (CYP2A-C and CYP3A). Indicative CYP enzymes are listed in brackets. Individual values and mean + SEM; n = 8 mice/group. (F) Enzyme activity of GST and UGT in mouse liver homogenates. GST: gluthathione S-transferase; UGT: UDP-glucuronosyltransferase. Individual values and mean + SEM; n = 8 mice/group. (G-I) GSH levels (G), GSH/GSSG ratio (H) and lipid peroxidation (I) in mouse liver and kidney homogenates. Individual values and mean + SEM; n = 7 (liver GSH and GSH/GSSG, kidney lipid peroxidation) or n = 8 (kidney GSH and GSH/GSSG, liver lipid peroxidation) mice/group. *P < 0.05, **P < 0.01, ***P < 0.001 vs. control; two-tailed unpaired Student’s t-test.

Based on these data, we speculated that, in healthy mice receiving silybin for a short period of time (24 h), the increase in intracellular membranes is functionally coupled to membrane protein biosynthesis and accompanied by an increased availability of membrane-bound phase I and II isoenzymes that metabolize and detoxify xenobiotics^137,138^. In fact, the protein levels of CYP3A2 and CYP2B1 (but not CYP1A1) were markedly enhanced in the liver of mice receiving silybin hemisuccinate, as shown by immunoblotting (Figure 8C and S15A) and visibly confirmed for all CYP isoforms tested by immunohistochemical analysis (Figure 8D). By contrast, the total amount of hepatic proteins decreased (Figure S15B). CYP enzyme expression was mainly concentrated around the endothelial cells of the central veins. Accordingly, the biotransformation activity of CYP enzymes (Figure 8E), GST and UDP-glucuronosyltransferase (UGT) increased strongly (Figure 8F), as determined by the conversion of indicative substrates, possibly to support silymarin glucuronidation and excretion^42^. Likely as a consequence of the increased GST turnover, the hepatic glutathione (GSH) pool decreased, with both GSH levels and the ratio to glutathione disulfide (GSSG) being significantly reduced (Figure 8G and H). Since GSH, as an essential co-substrate of glutathione peroxidase (GPX)4, contributes to the reduction of lipid hydroperoxides and prevents degenerative cell death^139^, we speculated that the decrease in GSH might enhance lipid peroxidation, which was, however, not the case (Figure 8I). Silybin actually attenuated the formation of lipid peroxidation products by trend in the liver but not in the kidney (Figure 8I), consistent with previous studies on silymarin/silybin^86,140,141^.

Together, the silybin-mediated accumulation of hepatic phospholipids (Figure 1B-D) is associated with an upregulation of membrane-bound detoxifying enzymes as well as GST isoenzymes that are present in different subcellular membrane compartments, including cytosol, mitochondria, ER, plasma membrane and nucleus^142^.

For information on the effects of silymarin/silybin on vitamin A metabolism, see Supplementary Note 2.

## 3. Discussion

The efficacy of silymarin and its major active component, silybin, in alleviating toxic liver injury and metabolic diseases^59^ has been ascribed to hepatoprotective, anti-inflammatory, anti-oxidative response-inducing and membrane-stabilizing properties as well as to lipid-(TG and cholesterol)-lowering effects^44,66,143^. Here, we report that silybin induces a metabolic switch in hepatocytes and extrahepatic cells, especially under non-or pre-disease conditions, linking hepatic TG metabolism with membrane biogenesis and potentially biotransformation activity (Figure 9), with the latter potentially contributing to the liver protective function.

**Figure 9.**
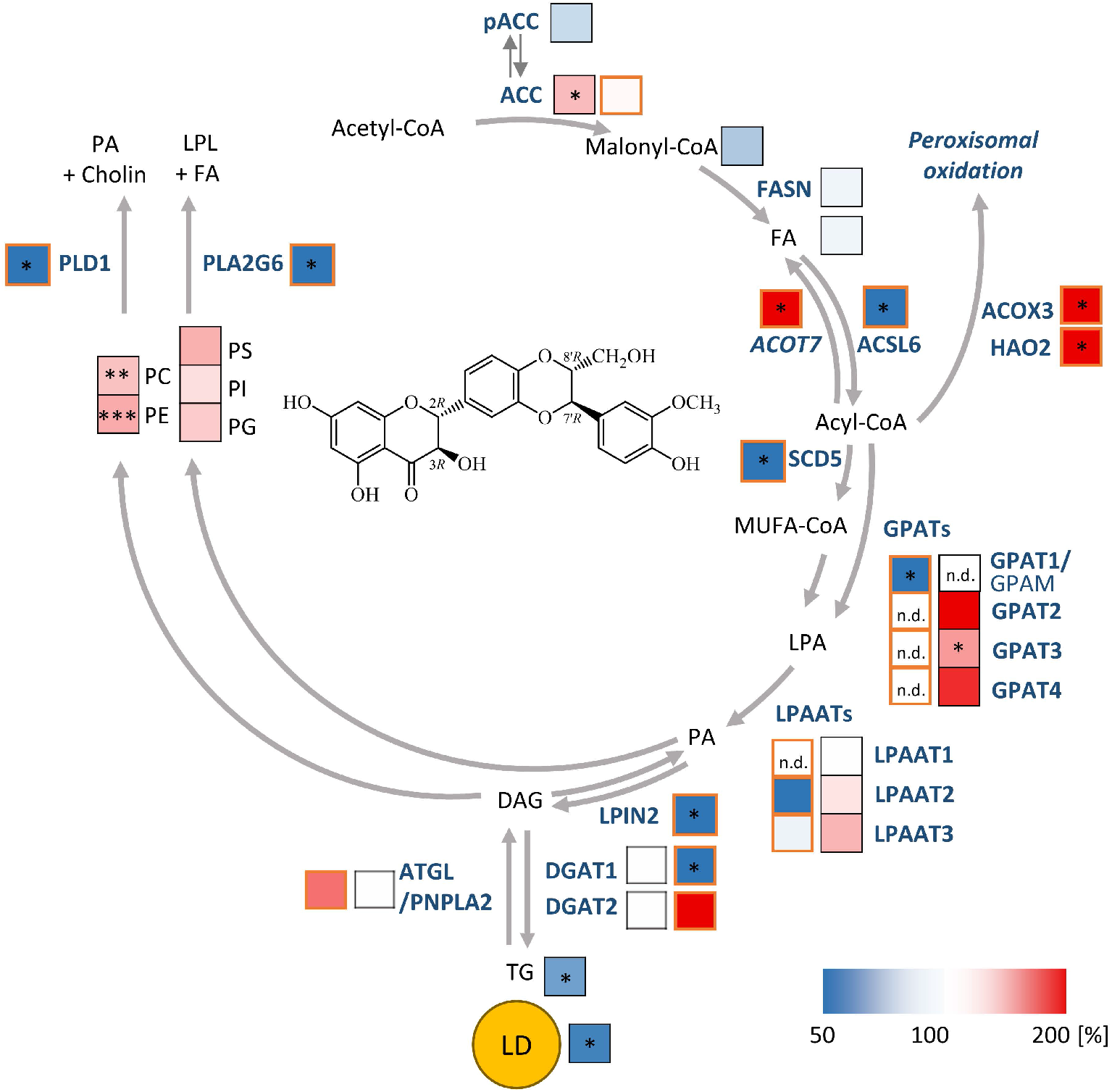
Proposed mechanisms of silymarin and its bioactive constituent silybin A in hepatocytes. Acetyl-CoA-carboxylase (ACC/ACACA) converts acetyl-CoA to malonyl-CoA, which is elongated to long-chain fatty acids by fatty acid synthase (FASN). Acyl-CoA esters are formed from free fatty acids (FAs) by acyl-CoA synthetases (ACSLs), which also activate exogenous fatty acids for further metabolism. Saturated acyl-CoAs are converted into monounsaturated acyl-CoAs (MUFA-CoA) preferentially by Δ9-desaturases, such as the stearoyl-CoA desaturase (SCD) isoenzyme 5. Acyl-CoA thioesterases (ACOTs) catalyze the opposite reaction, hydrolyzing acyl-CoAs to free fatty acids (FAs). Acyl-CoAs are used by glycerol-3-phosphate acyltransferases (GPATs) and lysophospholipid (LPL) acyltransferases/lysophosphatidic acid acyltransferases (LPLATs/LPAATs) to introduce fatty acyl-chains into the sn-1 and sn-2 positions of glycerol-3-phosphate and lysophosphatidic acid (LPA), respectively. The resulting PA is either converted to CDP-DAG for PI, PG, and PS biosynthesis or dephosphorylated to DAG for TG, PC, and PE biosynthesis by lipins (LPINs) and other PA phosphatases. LPIN2 also plays an important role in the regulation of fatty acid metabolism as nuclear transcriptional coactivator. Acylation of DAG by DGATs yields TGs, which are stored in lipid droplets and mobilized by ATGL/PNPLA2 and other triglyceride lipases, providing DAG and FAs. Phospholipid degradation is driven by a large number of phospholipases with different specificities. PLA2G6 releases saturated and unsaturated long-chain fatty acids from the sn-1 or sn-2 position of phospholipids, such as PC, PE and PA, whereas PLD1 specifically cleaves PC to PA and choline. By targeting multiple nodes, silymarin/silybin triggers a switch from TGs to phospholipids, thereby enriching intracellular membranes with phospholipids that have a balanced fatty acid composition. The increase in intracellular membranes is associated with enhanced membrane-associated biotransformation capacities. Mechanistically, silymarin/silybin inhibits phospholipid degradation, while moderately activating de novo phospholipid biosynthesis and stimulating TG catabolism in lipid droplets (LD), which in combination results in an effective channeling of TG-derived DAG and FAs into membrane biogenesis. In addition, silymarin induces the expression of genes involved in peroxisomal fatty acid degradation (HAO2, ACOX3), upregulates ACOT7, which hydrolyzes acyl-CoAs into FAs and CoA, and decreases the expression of ACSLs, that activate long-chain fatty acids. The color scale in the pathway diagram indicates the percentage changes in metabolite levels, lipid droplet counts, and enzyme expression by silybin relative to vehicle control in HepG2 cells (black bordered boxes) or by silymarin relative to vehicle control in HepG2 cells (orange bordered boxes). GPAM, glycerol-3-phosphate acyltransferase, mitochondrial.

Specifically, silybin treatment lowers TG levels, while limiting phospholipid degradation in hepatocytes and, under certain settings, additionally stimulates phospholipid biosynthesis, reflecting a net transfer of fatty acids from TGs to phospholipids (Figure 9). Context-dependent adaptations of fatty acid biosynthesis, intracellular transport, mitochondrial and peroxisomal degradation, cholesterol biosynthesis, and sterol metabolism further add to the class switch from TGs to phospholipids. Consequently, the number of lipid droplet decreases while the content of membrane phospholipids increases. At the same time, intracellular membranes are formed that, when coordinated with an upregulation of phase I and II membrane-(associated) enzymes, may enhance the biotransformation capacity of subcellular compartments, such as the ER.

A decrease of hepatic lipid droplets and TG levels is generally considered beneficial in metabolic diseases^13,144^. In support of this principle, increased ATGL/PNPLA2 expression protects against hepatic steatosis^145^, whereas ATGL/PNPLA2 repression promotes the development of MAFLD^146,147^. On the other hand, lipolysis is also associated with elevated levels of free fatty acids, which in excess may be lipotoxic to hepatocytes or cause oxidative stress when being degraded by mitochondrial or peroxisomal β-oxidation^5,148^. Thus, suppression of TG biosynthesis by selective inhibition of DGAT2 improves steatohepatitis and insulin sensitivity, but at the same time exacerbates liver damage in a methionine and choline deficient (MCD) mouse model of NASH^5^. However, in alternative animal models (such as those using diets high in fructose, saturated fat, and cholesterol^149^, or Western diets^150^), DGAT2 inhibition reduced steatosis without affecting inflammation or fibrosis in the latter. Conclusively, the reduction of lipid droplets and TGs alone does not fully explain the hepatoprotective function of silymarin and silybin, but requires an efficient channeling of the degradation products into non-toxic metabolites, i.e., phospholipids, as suggested by our results. A similar redistribution was observed for the inhibition of DGAT2 (PF-06424439), although it was restricted to the ER and PE species^151^. This metabolic switch to phospholipids is likely to be of biomedical relevance in toxic liver injury and MAFLD/MASH, where either the hepatic content of total phospholipids or specific phospholipid subclasses is decreased^41,61,62,77^. Consistent with this metabolic dysregulation, many key regulatory factors of MAFLD involve enzymes that are central to phospholipid and TG metabolism, including PNPLA3^31^, LPIAT1/MBOAT7^20,30,31,122,152^, ATGL/PNPLA2^23^, iPLA2/PLA2G6^24,25^, PLA2G7^153^, PLA2 activity of PRDX6^28^, PLD1^29^, and LPIN2^22^. Altogether, genetic variations or changes in protein expression of these enzymes define the risk of developing MAFLD^21,154^ and, together with other regulatory mechanisms, may shape the aberrant lipid composition of the diseased liver, with decreasing phospholipids and increasing TGs^33-41^. Silymarin/silybin regulates a significant number of these lipid metabolic genes, including *iPLA2/PLA2G6, ATGL/PNPLA2, PLD1, LPIN2*, and by trend *PNPLA3* (which is a major genetic risk factor for MAFLD^122^) and *HSD17B13*, counteracting the observed dysregulation in liver diseases. In line with our findings, a recently published randomized controlled trial showed that silybin treatment The flavonolignan silybin favorably redistributes lipids 20 improved MAFLD parameters only in patients without a genetic predisposition, while it was ineffective in patients carrying either one or a combination of mutations responsible for genetically inherited forms of MAFLD^122^. Given that the metabolic and genetic components of MAFLD differ fundamentally^155^, these findings suggest that silymarin/silybin may be particularly effective against the metabolic, but not against the genetic, form of MAFLD.

In addition, we show here that silymarin/silybin represses the potentially disease-promoting oxidative metabolism of fatty acids (via mitochondrial and peroxisomal pathways but also CYP enzymes) in many settings, including primary mouse hepatocytes, although opposite regulations were also observed at the transcriptome level.

Diverse mechanisms have been discussed for the TG- and cholesterol-lowering activity of silymarin/silybin: i) reduced lipid resorption, ii) upregulated cholesterol efflux *via* (ABC) transporters that excrete cholesterol from the liver to the bile^66,85,156^, and iii) adjustments in lipid biosynthesis, transport, and degradation by targeting major transcription factors in lipid metabolism^72,76,85,157-160^, such as PPARα/PPARγ, LXR; ChREBP and SREBP-1c^67,159-163^. Lipid-metabolic proteins proposed to be affected by silymarin/silybin include enzymes involved in fatty acid biosynthesis (ACC/*ACACA*, FASN, SCD-1), uptake (FABP5), and degradation (CPT1α, ACOX), phospholipid biosynthesis (GPATs), lipid transport (MTTP), and phosphatidic acid/TG turnover (PNPLA3)^72,76,85,157-160,164^. While these studies focused on the ability of silymarin/silybin to restore expression levels under pathophysiological conditions, we first addressed non-stressed cells, healthy mice, and pre-disease conditions. Comparative transcriptomic analyses in four different model systems confirmed that silymarin/silybin differentially regulates pathways contributing to TG and phospholipid metabolism. Specifically, our analysis revealed that silybin broadly manipulates phospholipid metabolism, although the exact mechanism varies between model systems and experimental settings. Overall, we show that silymarin/silybin reduces phospholipid degradation by repressing various phospholipases (*PLA2G1B, PLA2G6, PLD1*, and/or *PLD6*) or inducing the expression of phospholipase-suppressing factors (e.g., *PLA2R1*). However, the specific enzymes targeted can vary between different experimental models, and in some cases alternative enzymes may be upregulated, potentially acting as compensatory mechanisms. Less consistently, silymarin/silybin also upregulates the expression of factors involved in acyl-CoA supply (e.g., *FASN, SLC27A1, ELOVL7, ACSL4)* and phospholipid biosynthesis (e.g., *MBOAT2, LPGAT1*), *via* both *de novo* and remodeling pathways. On the other hand, silymarin/silybin suppresses TG biosynthesis mainly by repressing enzymes involved in the generation of DAGs and their acylation to TGs (e.g., via *DGAT1* and *DGAT2*).

The important role of silymarin/silybin in modulating lipid metabolism has been recognized before, and effects on ACC^67,89,165^, FASN ^67,89,156,160,165-167^, SCD-1^15,85,160,168^, GPATs^85^, PNPLA3^85,164,166^, FABP5^85,160,168^, CPT1a^67,85,89,160,167,169^, MTTP^85^, ACOX^85^, PPARα/PPARγ^67,85,160,164,166,169,170^ and SREBP-1c^67,85,164-166,169,170^ have been reported independently, either under disease^67,75,76,85,141,158,159,164,165^ or non-disease conditions^72,156,166,167^, largely without considering their interplay. By converging transcriptomics, metabololipidomics, and functional studies, we put these individual findings into context. Thus, our data strongly suggest that silybin, by coordinating multiple enzymes involved in lipid metabolism, facilitates the efficient channeling of fatty acids from TGs into phospholipids unless cells experience extensive lipid overload, with potential implications for disease prevention. The liver and body weight of the treated mice were reduced accordingly. We conclude that silybin buffers excessive hepatic TG accumulation, a hallmark of MAFLD, and redirects fatty acids by limiting phospholipid degradation and stimulating (energy-consuming) phospholipid biosynthesis and possibly membrane biogenesis.

Gavage of silymarin/silybin to mice reduced pathological changes in liver and serum lipid composition^62,77,78,171,172^, and its beneficial effects were anticipated to depend on either a decreased cholesterol/phospholipid ratio^77^, a reduced proportion of SM relative to PC^77,171^, or increased PE levels^78^. Our lipidomic analysis essentially confirmed an efficient upregulation of PE and other glycerophospholipids (rather than sphingolipids) in mouse liver by silybin. The influence on membrane properties is difficult to assess, but the homogeneous upregulation of phospholipid classes suggests that there are no major changes. It should be noted that we did not analyze free cholesterol, a major membrane component that affects rigidity and fluidity^173^.

Reported cytostatic, growth-inhibitory, or cytotoxic effects of silymarin/silybin on cancer cells are usually observed at much higher concentrations^174-176^ than those that we found to induce the switch from TG to phospholipids in hepatocytes. Thus, silymarin/silybin did not induce G2/M cell cycle arrest, trigger ER-stress, or reduce hepatocyte viability (except for an increase in the UPR marker BiP after 72 h treatment with silymarin) at the concentrations and time points used here. These results rather exclude that non-specific stress (adaptive) responses drive the observed changes in hepatocyte lipid metabolism.

Structure-activity relationship studies underscore that silybin functionally intervenes at two (or more) different sites in lipid metabolism to accomplish the shift from TGs to phospholipids. While the saturation of the flavonoid scaffold at the 2,3-position is essential for phospholipid accumulation (but has little effect on the amount of cellular TG), changes in the stereochemistry at the dioxan ring reduced both the effect on phospholipid and TG levels. The introduction of a 2,3-double bond yielding 2,3-dehydrosilybin even resulted in a decrease of the PS and PG content. The biosynthesis of these acidic phospholipids requires the conversion of DAG to phosphatidic acid, whereas PC and PE can be synthesized directly from DAG (Figure 9)^177^. Interestingly, the ring rearrangement in the hemiacetal did not substantially hamper the activity on phospholipids or TGs when compared to the diastereomeric silybin mixture. Together, silybin modulates phospholipid and TG metabolism through independent pathways, with the magnitude and directionality of the effect strongly dependent on the stereochemistry and saturation of the flavonolignan. Consistent with the hypothesis that silybin upregulates the intracellular phospholipid content also independently of the decrease in TGs, pharmacological inhibition of specific isoenzymes involved in lipid droplet degradation (ATGL) or lipid droplet biogenesis (DGAT2) did not markedly alter the cellular phospholipid content.

We also show that silymarin/silybin increases the content of phospholipids in hepatocytes, thereby forming intracellular membranes that are likely to host enzymes involved in biotransformation. On the one hand, phase I and phase II enzymes provide protection against multiple xenobiotics and diminish drug-induced hepatotoxicity^178-180^. On the other hand, phase I CYP enzymes convert various xenobiotics, e.g., the analgesic drug acetaminophen (paracetamol), into toxic metabolites^181^. Silymarin has been proposed to protect against toxic liver injury i) by inhibiting CYP enzymes and suppressing deleterious metabolism, ii) by inducing the expression of phase II enzymes such as UGT and GST, and iii) by upregulating membrane transporters that enhance the excretion of xenobiotics^127,128^. While our data confirm an upregulation of phase II enzyme activities by silybin, we found that not only the expression but also the activity of multiple CYP isoenzymes was increased rather than decreased under short-term treatment. We suggest that the mixed outcomes of studies investigating the effect of silymarin/silybin on CYP enzymes originate from kinetic regulation and the competition between CYP expression and inhibition, which seems to be sensitive to the dosage, route of application, formulation, and/or duration of treatment^82,127,129,182-184^. Thus, the elevated CYP enzyme activity in our experimental design is likely due to an increased CYP protein expression that masks the inhibitory effect of silybin on CYP activity. In support of this hypothesis, silymarin administration to rats increased the hepatic cytochrome P450 levels with defined kinetics^185^, as further validated here at the transcriptome level in HCV-infected mice treated with silybin. Overall, silymarin/silybin induced a rapid upregulation of drug-metabolizing enzymes, followed by a decrease in expression with prolonged treatment. Another factor that may contribute to the variable study results on CYP enzymes is that healthy and diseased tissues seem to respond differently to silybin. In fact, silymarin/silybin partially restores CYP enzyme homeostasis under pathophysiological conditions^131,132,134^, whereas effects in healthy individuals are more diverse^66,130,135,136,186,187^. It should be noted that, with a few exceptions, most *in vivo* animal and human studies have failed to confirm that silymarin/silybin substantially interferes with the pharmacokinetics of various drugs that are metabolized by CYP enzymes^42^. However, one of the few studies showing significant effects in healthy volunteers found, consistent with our results in mice, that silymarin (140 mg, daily) increased the clearance and decreased the C_max_ values of metronidazole, a drug that is metabolized by CYP3A4 and CYP2C9^188^. Whether the modulation of CYP enzyme activity by silymarin is of clinical relevance in patients with MAFLD remains elusive and needs to be systematically evaluated for specific formulations, dosages and CYP isoenzymes in future kinetic studies.

Given the mixed results of silymarin/silybin on lipid metabolism in health and disease^74,122-126^, we investigated the ability of silybin to redirect lipid metabolism in *in vitro* models of MAFLD (achieved by massive fatty acid overload) and acute lipotoxicity (induced by excess saturated fatty acids). On the one hand, silybin A consistently reduced lipid droplet content below basal levels in unstressed and fatty acid-challenged hepatocytes, notably superior to selective inhibitors of DGAT1 or DGAT2 or antagonist of PPARγ. On the other hand, silybin A was considerably less effective in redirecting fatty acids from lipid droplets to phospholipids in stressed as compared to non-stressed cells and did not protect against lipotoxicity. Our results suggest that the silymarin/silybin-induced lipid class switch from TGs to phospholipids is particularly effective in protecting against adverse dysregulation of lipid metabolism in normal and pre-disease conditions, whereas the beneficial effects appear to be largely limited to reducing TGs in liver diseases associated with severe TG accumulation, such as MAFLD. Further studies are needed to elucidate the general relevance of this dual mechanism by directly comparing healthy and diseased states in clinical trials.

Silymarin and silybin modulate lipid metabolism in hepatocytes in a comparable manner, but there are also substantial differences, and some effects are seen only for silymarin. First, silymarin significantly decreased the number of cells in S-phase while increasing the subG1 fraction, and induced ER stress after prolonged treatment for 72 h. These effects were not observed for silybin. Second, only silymarin elevated the levels of butyryl-CoA, an intermediate of β-oxidation (1.5-fold), implying that fatty acid degradation is stimulated by components of silymarin other than silybin. Whether such polypharmacological modulation by silymarin has advantages, is poorly understood. On the one hand, we discussed above that excessive β-oxidation under stress conditions could be detrimental due to the production of reactive oxygen species (ROS)^15^. On the other hand, the induction of fatty acid catabolism by silymarin is moderate and could help to meet the energy demands for stress-adaptive, regenerative pathways (including lipid remodeling), especially since the antioxidants in silymarin already counteract ROS accumulation^189^. In addition to modulating lipid metabolism, silymarin is known to have beneficially effects on other pathogenic mechanisms associated with MAFLD, such as inflammation and glucose metabolism^190^, the latter of which is also supported by our data

## 4. Conclusion

The milk thistle extract silymarin and its bioactive component silybin have unique lipid-modulating properties. In hepatic pre-disease states, silybin decreases TG biosynthesis, while attenuating phospholipid degradation and stimulating phospholipid biosynthesis. In combination with the parallel reprogramming of phase I and II metabolism, this lipid class switch seems to expand functional intracellular membranes and redirects the hepatic biotransformation capacity.

Considering that the selective reduction of TG stores actually enhances liver damage, as suggested by preclinical studies with DGAT2 antisense oligonucleotide treatment^5^, we propose that the lipid metabolic switch from TGs to phospholipids in hepatocytes and potentially other liver cells (i.e., Kupffer cells) and extrahepatic cells critically contributes to the liver-protective (rather than disease-alleviating) function of silybin. The beneficial reprogramming of lipid metabolism is based on the absolute configuration of silybin as well as the saturation of ring C in the flavonoid scaffold at the 2,3-position. These structural aspects contribute differentially to the TG-lowering and phospholipid-accumulating activities of silybin, and structural modifications realized in minor components of milk thistle allow dissection of both activities.

In conclusion, our results shed light on the mechanisms underlying liver protection by silymarin/silybin under physiological and pathophysiological conditions and may help to place the pleiotropic but mechanistically diffuse facets of their action on lipid metabolism in a broader context. Future studies are needed to explore the relevance of the silybin A-induced lipid metabolic switch in humans, both for disease prevention and under pathophysiological conditions.

## 5 Materials and Methods

### 5.1 Materials

Silybin, staurosporine, and atglistatin were obtained from Merck (Darmstadt, Germany), silybin-C-2’,3-bis(hydrogen succinate) disodium salt (Legalon^®^ SIL) was from Madaus GmbH (Köln, Germany), vinblastine, the PPARγ antagonist GW9662, and the DGAT1 inhibitor A-922500 were purchased from Cayman Chemicals (Ann Arbor, MI), the DGAT2 inhibitor PF-06424439 was bought from Bio-Techne (Abingdon, United Kingdom), thapsigargin was from Enzo Life Sciences (Farmingdale, NY), and silymarin (Silimarit®) was a kind gift from Bionorica SE (Neumarkt, Germany). Silybin, its derivatives and other compounds were dissolved in DMSO, stored in the dark at -20°C under argon, and freezing/thawing cycles were kept to a minimum. Silymarin was freshly dissolved in ethanol at the day of experiment. Phospholipid standards were purchased from Otto Nordwald GmbH (Hamburg, Germany) or Merck Millipore (Darmstadt, Germany), were dissolved in chloroform, aliquoted and stored under argon protected from light at -80°C. BODIPY 493/503 and ProLong™ Diamond Antifade Mountant with DAPI were purchased from Thermo Fisher Scientific (Waltham, MA). Rabbit anti-β-actin (13E5; #4970), mouse anti-β-actin (8H10D10; #3700), rabbit anti-acetyl-CoA carboxylase (C83B10; #3676), rabbit anti-ATF-6 (D4Z8V, #65880), rabbit anti-ATGL (#2138), rabbit anti-BiP (C50B12, #3177), rabbit anti-phospho-acetyl-CoA carboxylase (Ser79; D7D11; #11818), rabbit anti-GAPDH (D16H11; #5174), mouse anti-GAPDH (D4C6R; #97166), rabbit anti-FAS (#3189), and rabbit anti-XBP-1s (D2C1F, #12782S) were obtained from Cell Signaling (Danvers, MA). Mouse anti-calnexin (C8.B6; #MAB3126) was from Merck Millipore (Darmstadt, Germany) and mouse anti-GM130 (#610822) from BD Bioscience (San Jose, CA, USA). Goat anti-rat CYP1A1 (#219207), goat anti-rat CYP3A2, (#210167), and goat anti-rat CYP2B1, (#219207) were obtained from Daiichi Pure Chemicals Co. LTD (Tokyo, Japan). Rabbit anti-DGAT1 (NB110-41487SS) and rabbit anti-DGAT2 (NBP1-71701SS) were from Novus Biologicals (Abingdon, UK). Mouse anti-GRP78/BiP (A-10, #sc-376768) was purchased from Santa Cruz Biotechnology (Dallas, TX). Alexa Fluor 555 goat anti-mouse IgG (H+L) and Alexa Fluor 488 goat anti-rabbit IgG (H+L) were purchased from Life Technologies (MA, USA). Secondary antibodies for Western blot studies were from LI-COR Biosciences (Bad-Homburg, Germany) and Thermo Fisher Scientific. Peroxidase-conjugated avidin and the secondary biotinylated antibodies rabbit anti-mouse IgG and rabbit fblanti-goat used in immunohistochemical studies were from VECTASTAIN^®^ Elite ABC-Kit (Vector Laboratories, Burlingame, CA)

### 5.2 Synthesis of silybin derivatives

Silybin A and B were separated from the diastereomeric mixture silybin (Merck) by preparative HPLC as described^90^. Starting from the purified silybin A and B, the two enantiomers of 2,3-dehydrosilybin (A and B) were synthesized in good yields and optically pure by base-catalyzed oxidation under microwave heating^91^. The hemiacetal **11**, was obtained in good yield by the microwave conversion of silybin in pyridine at 110°C^91^. All products were fully characterized by NMR (^1^H, ^13^C), CD, [α]_D_, and ESI MS analyses. The purities of the products were higher than 98%.

### 5.3 Cell culture, primary monocytes and cell treatment

Cultured cell lines: Human HepG2 liver carcinoma cells (1×10^5^ cells/cm^2^, Leibniz Institute DSMZ-German Collection of Microorganisms and Cell Cultures, Braunschweig, Germany) were grown in RPMI 1640 medium containing 10% heat-inactivated fetal calf serum (FCS, GE Healthcare, Freiburg, Germany or Merck) at 37°C and 5% CO_2_. Human HepaRG hepatoma cells (1.5-2×10^5^ cells/cm^2^, Biopredic International, Rennes, France) were cultured in William’s E medium (Merck) supplemented with 10% heat-inactivated FCS, 2 mM *L*-glutamine (Merck), 5 μg/ml human insulin (Merck), and 50 μM hydrocortisone (Cayman) at 37°C and 5% CO_2_. Human Caco-2 colorectal adenocarcinoma cells (1.7×10^5^ cells/cm^2^) were cultured in DMEM medium (Merck) containing 10% FCS at 37°C and 5% CO_2_. Cells were detached by trypsin/EDTA and reseeded every 3-4 days before reaching confluence.

Primary cells: Collection of venous blood in heparinized tubes (16 I.E. heparin/mL blood) was performed by the Institute for Transfusion Medicine of the University Hospital Jena (Germany) with informed consent of registered male and female healthy adult volunteers (18 to 65 years). Blood donors were fasted for at least 12 h, had not taken antibiotics or anti-inflammatory drugs prior to blood donation (> 10 days), and were free of apparent infections, inflammatory disorders, or acute allergic reactions. The volunteers regularly donated blood (every 8 to 12 weeks) and were physically inspected by a clinician. Leukocyte concentrates were prepared, erythrocytes removed by dextran sedimentation, and peripheral blood mononuclear cells (PBMC) were isolated by density gradient centrifugation on lymphocyte separation medium (LSM 1077, GE Healthcare) as previously described^191^. The fraction of PBMC was cultivated in RPMI 1640 medium containing 10% FCS in 12-well plates (37°C, 5% CO_2_) at a density of 2×10^7^/ml for 1 to 1.5 h to separate adherent monocytes. The cell population used for further studies consisted of more than 85% monocytes according to forward and side scatter properties and CD14 surface expression (BD FACS Calibur flow cytometer, BD Biosciences, Heidelberg, Germany). Experiments were approved by the ethical commission of the Friedrich-Schiller-University Jena.

Cell treatment: HepG2 cells (1×10^5^ cells/cm^2^) and monocytes (6×10^5^/cm^2^) were seeded and directly exposed to vehicle (0.1% DMSO or 0.05% ethanol), silymarin (50 μg/ml for monocytes and 10 μg/ml for HepG2 cells), silybin A/B (20 μM), STS (1 μM), or vinblastine (100 nM). Adherent cells were harvested with trypsin/EDTA (Merck or Promega, Madison, WI). For lipid droplet staining with Oil Red O, HepG2 cells were instead seeded in 96-well plates at 20,000 cells per well and incubated for 24 h before treatment with vehicle (0.5% DMSO or 0.5% ethanol), silymarin (10 μg/ml), or silybin A/B (20 μM) for an additional 24 h. Treatment of HepaRG cells is described in sections 5.5. For transcriptome analysis, Caco-2 cells (1.7×10^5^ cells/cm^2^) were seeded and directly exposed to vehicle (0.5% DMSO), silymarin (30 μg/ml), and silybin (30 μM) for 24 h. Adherent cells were harvested with trypsin/EDTA.

### 5.4 Complexation of fatty acids to BSA

BSA (1%, Carl Roth, Karlsruhe, Germany) was dissolved in Williams E medium, sterile filtered (Rotilabo^®^-syringe filter, PVDF, 0.22 μm, Carl Roth), mixed with PA (50 mM) or OA (50 mM), sonicated at 60°C for 30 min using a USC100TH sonicator (VWR, Vienna, Austria, 60 W, 45 kHz), and stored at -20°C. Solutions were mixed vigorously immediately before use.

### 5.5 Cell-based models of MAFLD and lipotoxic stress

HepaRG cells (10,000 / well, 96-well plate) or 2.5×10^6^ cells/25 cm^2^ were cultured at 37°C and 5% CO_2_ for 24 h. The cell culture medium was replaced with fresh medium supplemented with i) vehicle (1% BSA in Williams E medium), ii) BSA-complexed PA/16:0 (0.1 mM, Merck) and OA/18:1 (Cayman) in a 1:2 ratio (in total 1 mM) to induce massive lipid accumulation (mimicking MAFLD), or iii) BSA-complexed PA (0.1 mM) to induce lipotoxic stress. For lipidomic analysis, cells were either co-treated directly with vehicle (DMSO, 0.5%) or silybin A (20 μM), and the incubation was prolonged for another 24 h. Alternatively, treatment was started 24 h after fatty acid challenge and incubation was prolonged for a further 24 h. For lipid droplet analysis, cells were co-treated with vehicle (DMSO, 0.5%), silybin A (20 μM), the ATGL inhibitor atglistatin (50 μM), the DGAT1 inhibitor A 922500 (5 μM), the DGAT2 inhibitor PF-06424439 (10 μM), a combination of DGAT1 (5 μM) and DGAT2 inhibitors (10 μM), or the PPARγ antagonist GW9662 (5 μM) and the incubation was prolonged for another 24 h or 48 h, respectively. Lipid droplet signals, the number of viable cells and membrane integrity, cellular metabolic activity, and phospholipid and TG levels were determined as described in sections 5.6, 5.7, 5.8, 5.10, and 5.11, respectively.

### 5.6 Quantitation of lipid droplets in hepatocytes

HepaRG cells were washed twice with 100 μl PBS pH 7.4 and fixed with paraformaldehyde solution (4% in PBS pH 7.4, Merck) for 40 min at room temperature. After removal of the fixative, the cells were washed twice with 100 μl of water, incubated with aqueous isopropanol (60%, 100 μl, 5 min) to remove polar lipids and reduce background signals, and stained with Oil Red O solution (50 μl) for 25 min at room temperature. The latter was prepared by diluting 0.5% Oil Red O in isopropanol (Merck) 1.7-fold in water, sterile-filtered (Rotilabo^®^-syringe filter, PVDF, 0.22 μm, Carl Roth), and allowed to stand for 10 min before staining. Cells were washed three times with water, and microscopic images were taken using a 40× objective (Motic, Barcelona, Spain) on a Motic AE31E microscope (Motic) equipped with a Motic camera. Alternatively, lipid droplets in HepG2 cells were stained with BODIPY 493/503 and manually counted as described in section 5.15. For photometric quantitation of the stained lipid droplets, Oil Red O was extracted with 60% isopropanol in water (100 μl) for 10 min at room temperature, and the absorbance of the extracted solution was measured at 510 nm using a multi-mode microplate reader (SpectraMax iD3, Molecular Devices).

### 5.7 Cell number, viability, morphology, and cell diameter

Cell number, cell viability, and cell diameters were determined after trypan blue staining using a Vi-CELL Series Cell Counter (Beckmann Coulter GmbH, Krefeld, DE). Morphological analysis of the cells was carried out on an Axiovert 200 M microscope with a Plan Neofluar × 100/1.30 Oil (DIC III) objective (Carl Zeiss, Jena, Germany). Images were obtained using an AxioCam MR3 camera (Carl Zeiss).

### 5.8 Cell viability based on cellular dehydrogenase activity

Cytotoxic effects of silymarin and silybin were determined as described^192^. Briefly, HepG2 cells (1×10^5^/well of a 96-well plate) or HepaRG cells were cultured as described in sections 5.3. and 5.5. Cells were treated with silymarin, silybin, or vehicle (0.5% DMSO or 0.25% ethanol) at 37°C and 5% CO_2_. The pan-kinase inhibitor staurosporine (1 μM) was used as reference compound. After 24 h, 3-(4,5-dimethylthiazol-2-yl)-2,5-diphenyltetrazolium bromide (MTT, 20 μl, 5 mg/ml, Merck) was added to each well, and cells were incubated for another 3 h (HepG2) or 2.5 h (HepaRG) at 37°C and 5% CO_2_ before being lysed in SDS buffer (10% in 20 mM HCl, pH 4.5) overnight. The absorption of the solubilized formazan product was measured at 570 nm (Multiskan Spectrum, Thermo Fisher Scientific or SpectraMax iD3, Molecular Devices).

### 5.9 Cell cycle analysis

HepG2 cells were fixed by dropwise addition of the cell suspension (in 1 ml PBS pH 7.4) to ice-cold ethanol (70%, 9 ml) under shaking. Fixed cells were stored overnight at 4°C, washed with ice-cold PBS pH 7.4, and pelleted (700×g, 5 min). Cell pellets were stained with 50 μg/ml propidium iodide (Thermo Fisher Scientific) in PBS pH 7.4, 0.1% Triton X-100, and 100 μg/ml DNase-free RNase A. After incubation at room temperature in the dark for 20 min at 37°C, cells were subjected to flow cytometric analysis (BD LSR Fortessa flow cytometer, BD Biosciences, Franklin Lakes, NJ). Data analysis was accomplished with Flowlogic software (Miltenyi Biotech, Bergisch Gladbach, Germany).

### 5.10 Extraction and analysis of phospholipids, neutral lipids, and fatty acids

To extract lipids from cell pellets (HepG2 cells, HepaRG cells, and monocytes) or supernatants of liver homogenates after centrifugation (9,000×g, 10 min, 4°C), PBS pH 7.4, methanol, chloroform, and saline (final ratio: 14:34:35:17) were added in succession^193,194^. Phospholipids, TGs and fatty acids in the lower organic phase were evaporated to dryness, dissolved in methanol, and analyzed by UPLC-MS/MS. Internal standards: 1,2-dimyristoyl-*sn*-glycero-3-phosphatidylcholine, 1,2-dimyristoyl-*sn*-glycero-3-phosphatidylethanolamine, 1,2-di-heptadecanoyl-*sn*-glycero-3-phosphatidylglycerol, 1,2-diheptadecanoyl-*sn*-glycero-3-phosphoserine, 1,2-dioctanoyl-*sn*-glycero-3-phospho-(1’-myo-inositol), and/or (15,15,16,16,17,17,18,18,18-d9)oleic acid. Alternatively, 1-pentadecanoyl-2-oleoyl(d7)-*sn*-glycero-3-phosphoethanolamine, 1-pentadecanoyl-2-oleoyl(d7)-*sn*-glycero-3-phosphocholine, and/or 1,3-dipentadecanoyl-2-oleyol(d7)-glycerol were used for lipidomic analysis related to Fig. 7 and Fig. S13.

Phospholipids, TGs, and free fatty acids were separated on an Acquity™ UPLC BEH C8 column (1.7 μm, 2.1×100 mm, Waters, Milford, MA, USA) using an Acquity™ Ultraperformance LC system (Waters) as described before^195-197^. Alternatively, phospholipids and TGs were separated by an ExionLC™ AD UHPLC (Sciex, Framingham, MA, USA) ^198-200^. In brief, phospholipids were analyzed at a flow rate of 0.75 ml/min at 45°C using acetonitrile/water (95/5) and 2 mM ammonium acetate as mobile phase A and water/acetonitrile (90/10) and 2 mM ammonium acetate as mobile phase B. Mobile phase A was ramped from 75 to 85% within 5 min, followed by an increase to 100% within 2 min and isocratic elution for another 2 min. For the separation of TGs, mobile phase B was replaced by isopropanol, and the initial composition of mobile phase A was lowered from 90 to 70% within 6 min, which was succeeded by isocratic elution for 4 min.

Glycerophospholipids were detected by multiple reaction monitoring in the negative ion mode based on their fatty acid anion fragments using a QTRAP 5500^196^ or QTRAP 6500^+^ Mass Spectrometer (Sciex), which were equipped with electrospray ionization (ESI) sources. For the analysis of PE and PC using the QTRAP 6500^+^ Mass Spectrometer (Figure 7, Figure S3, and Figure S13), the curtain gas was set to 40 psi, the collision gas was set to medium, the ion spray voltage was set to -4500 V, the heated capillary temperature was set to 650°C (PE) or to 350 °C (PC), the sheath gas pressure was set to 55 psi, the auxiliary gas pressure was set to 75 psi, the declustering potential was set to -50 V, the entrance potential was set to -10 V, the collision energy was set to -38 eV, and the collision cell exit potential was set to -12 V^200^.

TGs were identified and quantified in the positive ion mode as NH_4_^+^ adduct ions that undergo neutral loss of either of the acyl groups^197^. When using the QTRAP 6500^+^ Mass spectrometer (Figure 7, Figure S3, and Figure S13), the curtain gas was set to 40 psi, the collision gas to low, the ion spray voltage to 5500 V, the heated capillary temperature to 400°C, the sheath gas pressure to 60 psi, the auxiliary gas pressure to 70 psi, the declustering potential to 120 V, the entrance potential to 10 V, the collision energy to 35 eV, and the collision cell exit potential to 26 V^200^.

Free fatty acids were analyzed by single ion monitoring in the negative ion mode^193^ and SM by multiple reaction monitoring in the positive ion mode based on the detection of the choline headgroup (m/z = 184)^193^.

Absolute lipid quantities were normalized to internal standards and either cell number or tissue weight. Relative intensities represent the percentage of individual lipid species relative to all lipid signals determined within the respective lipid class (= 100%). The most intensive or specific transition was used for quantitation. Analyst 1.6 or Analyst 1.7 (Sciex) were used to acquire and process mass spectra.

### 5.11 Extraction and analysis of acyl-CoAs

HepG2 cells were suspended in methanol/water (70/30) and placed at -20°C for 1 h. After vigorous mixing, the methanol/water ratio was adjusted to 50/50 and the samples were incubated for another hour at -20°C. Protein precipitates were removed by centrifugation (20,000×g, 5 min, 4°C), and the supernatant was evaporated to dryness. The residue was extracted with methanol/water (50/50) and the extract subjected to UPLC-MS/MS analysis. [^13^C_3_]-Malonyl-CoA (1 nmol; Merck) was used as internal standard.

Acyl-CoAs were separated on an Acquity™ UPLC BEH C18 column (1.7 μM, 2.1×50 mm) with an Acquity™ Ultra Performance LC system^201^ and analyzed by multiple reaction monitoring in the positive ion mode following electrospray ionization (QTRAP 5500 mass spectrometer). Fragments formed by neutral loss of 2’-phospho-ADP ([M+H-507]^+^) were detected for quantitation. The ion spray voltage was set to 3,000 V, the heated capillary temperature to 600°C, the curtain gas pressure to 30 psi, the sheath gas pressure to 45 psi, the auxiliary gas pressure to 55 psi, the declustering potential to 60 V, the entrance potential to 10 V, and the collision energy to 45 eV (malonyl-CoA) or 30 eV (other acyl-CoAs). Absolute lipid amounts are calculated with respect to the internal standard of the subclass and are normalized to cell number, protein content or tissue weight. Relative lipid proportions are expressed as a percentage of the total sum of all species detected within the corresponding subclass (equal to 100%). Mass spectra were acquired and analyzed using Analyst 1.6 or 1.7 (Sciex).

### 5.12 Transcriptome analysis

Caco-2 cells (1.7×105 cells/cm2) were treated with vehicle (0.5% DMSO), 30 μg/ml silymarin or 30 μM silybin for 24 h (n = 3 biological replicates). Total RNA was isolated using a RNeasy Mini Kit (Qiagen) and potential DNA contamination was digested with DNase I (Qiagen) during RNA purification according to the manufacturer’s protocol. RNA concentration and quality were assessed using a SpectraMax iD3 microplate reader (Molecular Devices), a bioanalyzer (Agilent) and Qubit (Thermo Fisher Scientific) before being submitted to the MultiOmics Core Facility, Medical University of Innsbruck, for sequencing. The RNA integrity (RIN) of all samples was > 9.5 (out of 10) and no genomic DNA contamination was detected in any of the samples prior to RNA sequencing. Libraries were prepared using Lexogen’s Quant Seq 3’mRNA Seq Library Kit FWD with UMI protocol (Lexogen GmbH, Vienna, Austria). Quality validated libraries were multiplexed and sequenced at 150 bp read length using Illumina NovaSeq technology and the generated paired-end raw sequence data reads were quality controlled using FastQC and MultiQC202^202^.

Sequencing adapters and reads shorter than 50 base pairs were removed using Trim Galore (Galaxy version 0.6.7) to improve mapping quality, and reads were mapped to the GRCh38 human reference genome (December 2013) using the RNAStar aligner (Galaxy version 2.7.10b)^203^. Final transcript count data were generated with HTSeq framework (Galaxy version 2.0.5)^204^ for high-throughput sequencing data based on the Ensemble release Homo_sapiens.GRCh38.107 gene annotation with default settings. All analyses were performed on a public instance of Galaxy at usegalaxy.eu. Differential gene expression analysis was performed using DESeq2 package version 1.26^205^ with an adjusted *P*-value < 0.05 (5% FDR).

In addition, we re-analyzed microarray-based transcriptome datasets: i) HepG2 cells treated with vehicle (0.0125% DMSO) or 12 μg/ml silymarin (Merck) for 24 h (n = 3 biological replicates)^98^; ii) Huh7.5.1 cells treated with vehicle (0.32% DMSO) or 40 μg/ml silymarin (Madaus Group, Cologne, Germany) for 4, 8, or 24 h (pooled triplicates in three [silymarin, 8 h; silymarin, 24 h], four [vehicle and silymarin, 4 h], or five [vehicle, 8 and 24 h] technical replicates)^99^; iii) primary human hepatocytes from chronically HCV-infected chimeric mice with humanized livers either untreated or receiving 469 mg/kg silybin-C-2’,3-bis(hydrogen succinate) disodium salt (Legalon^®^ SIL, in saline, all three mice on day 3 and two mice on day 14) or 265 mg/kg Legalon^®^ SIL (in saline, one mouse on day 14) intravenously daily for 3 or 14 days (n = 3 mice/group)^100^. Data are accessible at NCBI GEO database^206^, accessions GSE67504, GSE50994, and GSE79103. Differentially regulated genes were identified by pairwise comparison of treatment and control groups using the GEO2R interactive web tool (https://www.ncbi.nlm.nih.gov/geo/geo2r/)^206^. *P* values were calculated by multiple *t*-tests, either with or without correction for multiple comparisons according to Benjamini and Hochberg (false discovery rate 5%) and auto-detection for log-transformation.

### 5.13 Sample preparation, SDS-PAGE, and Western blotting

Pelleted and washed monocytes and HepG2 cells were lysed in ice-cold 20 mM Tris-HCl (pH 7.4), 150 mM NaCl, 2 mM EDTA, 1% Triton X-100, 5 mM sodium fluoride, 10 μg/ml leupeptin, 60 μg/ml soybean trypsin inhibitor, 1 mM sodium vanadate, 2.5 mM sodium pyrophosphate, and 1 mM phenylmethanesulphonyl fluoride, and sonicated on ice (2 × 5 s, Q125 Sonicator, QSonica, Newtown, CT, 125 W, 35% amplitude). After centrifugation (cell lysates: 12,000×g, 5 min, 4°C; liver homogenates: 9,000×g, 10 min, 4°C), the protein concentration of the supernatants was determined using a DC protein assay kit (Bio-Rad Laboratories, CA). Samples (10-15 μg total protein) were combined with loading buffer (1×; 125 mM Tris-HCl pH 6.5, 25% sucrose, 5% SDS, 0.25% bromophenol blue, and 5% β-mercaptoethanol) and heated for 5 min at 95 °C. Proteins were separated by 8-10% SDS-PAGE and transferred to a Hybond ECL nitrocellulose membrane (GE Healthcare) or Amersham Protran 0.45 μm NC nitrocellulose membranes (Carl Roth, Karlsruhe, Germany). Membranes were blocked with 5% bovine serum albumin (BSA) or skim milk for 1 h at room temperature and incubated with primary antibodies overnight at 4°C. IRDye 800CW-labeled anti-rabbit IgG (1:10,000, 92632211, LI-COR Biosciences, Lincoln, NE), IRDye 800CW-labeled anti-mouse IgG (1:10,000, 926-32210, LI-COR Biosciences, Lincoln, NE), IRDye 680LT-labeled anti-rabbit IgG (1:80,000, 926-68021, LI-COR Biosciences, Lincoln, NE), IRDye 680LT-labeled anti-mouse IgG (1:80,000, 926-68020, LI-COR Biosciences, Lincoln, NE), DyLight^®^ 680 goat anti-rabbit IgG (1:10,000, # 35569, Thermo Fisher Scientific), and/or DyLight^®^ 800 goat anti-mouse IgG (1:10,000, # SA5-10176, Thermo Fisher Scientific) were used as secondary antibodies. Fluorescent, immunoreactive bands were visualized using an Odyssey infrared imager (LI-COR) or a Fusion FX7 Edge Imaging System (spectra light capsules: C680, C780; emission filters: F-750, F-850; VILBER Lourmat, Collegien, France) ^198^. Acquired data from densitometric analysis were linearly adjusted and background-corrected using Odyssey Infrared Imaging System Application Software Version 3.0 (LI-COR Biosciences) or Evolution-Capt Edge Software Version 18.06 (VILBER Lourmat) and Bio-1D imaging software Version 15.08c (Vilber Lourmat), and protein levels were normalized to GAPDH or β-actin.

### 5.14 Real-time quantitative PCR

HepG2 cells were incubated with silymarin (10 μg/ml), silybin (20 μM), or vehicle (ethanol for silymarin, DMSO for silybin) for 24 h. Total RNA of HepG2 cells was isolated with the E.Z.N.A Total RNA Kit (Omega Bio-tek, Norcross, GA).

SuperScript III First-Strand Synthesis SuperMix (Thermo Fisher Scientific) was used for transcription into cDNA. An aliquot of the cDNA preparation (1.25 μl) was combined with 1× Maxima SYBR Green/ROX qPCR Master Mix (Fermentas, Darmstadt, Germany) and forward and reverse primer (0.5 μM; TIB MOLBIOL, Berlin, Germany) in Mx3000P 96-well plates. Primer sequences are given in Table 1. β-Actin and GAPDH were used as reference. PCR was performed on a StraTGene Mx 3005P qPCR system (Agilent Technologies, Santa Clara, CA). The PCR program heats to 95°C for 10 min and conducts 45 cycles of 15 s at 95°C, 30 s at 61°C, and 30 s at 72°C. Threshold cycle values were determined by MxPro Software (Mx3005P/version 4.10, Agilent Technologies) and normalized to the amount of total RNA.

**Table 1.**
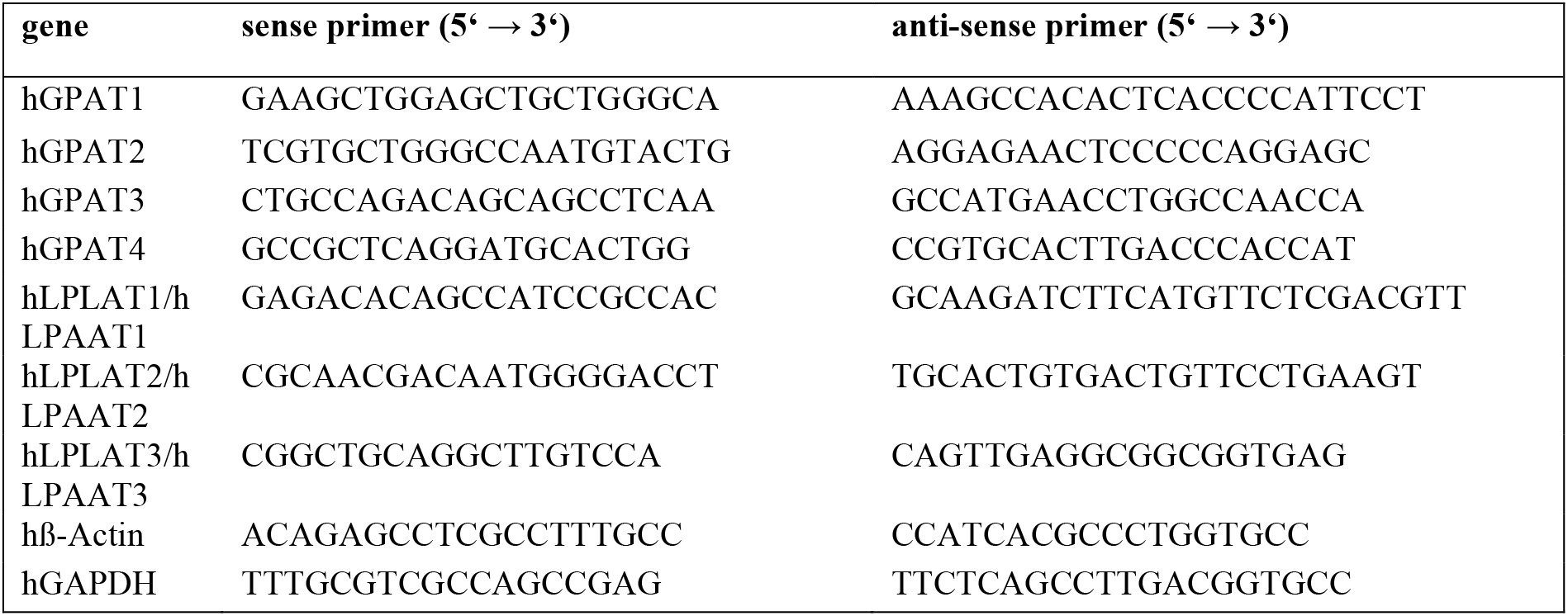
Primer sequences used in real-time quantitative PCR experiments.

### 5.15 Immunofluorescence microscopy

HepG2 cells (2.5×10^4^/3.9 cm^2^) were seeded onto poly-*D*-lysine-coated glass cover slips (Neuvitro Corp., Vancouver, WA) and cultured for 24 h. Vehicle, silymarin, or silybin were added, and cells were incubated for another 24 h. Hepatocytes were fixed with 4% paraformaldehyde for 20 min at room temperature and permeabilized with 0.25% Triton X-100 for 10 min at 4°C. After blocking with 5% normal goat serum (Thermo Fisher Scientific), samples were sequentially incubated with primary antibodies (anti-calnexin, 1:200; anti-GM130, 1:1000) at 4°C for 16 h and fluorescent-labeled secondary antibodies (Alexa Fluor 488 goat anti-rabbit IgG, 1:1000 or Alexa Fluor 555 goat anti-mouse IgG, 1:1000, Thermo Fisher Scientific) for 30 min at room temperature. For staining of lipid droplets, fixed cells were incubated with BODIPY 493/503 (0.1 μg/ml in 150 mM NaCl) for 5 min at room temperature under light exclusion. ProLong Diamond Antifade Mountant with DAPI (Thermo Fisher Scientific) was used to stain nuclear DNA. Fluorescent labeled organelles were visualized by an Axiovert 200 M microscope (Carl Zeiss), which was equipped with a Plan Neofluar×100/1.30 Oil (DIC III) objective and an AxioCam MR3 camera. Images were processed with AxioVision 4.8 software (Carl Zeiss).

### 5.16 Immunohistochemistry (IHC)

Liver samples were immediately fixed in neutral buffered 4% paraformaldehyde for at least 24 h and then dehydrated in increasing alcohol concentrations, embedded in paraffin, and sliced into 4 μm sections as described before^207^. The sections were deparaffinized with xylene and rehydrated using an inverse series of aqueous alcohol concentrations. Hydrogen peroxide (0.3% in methanol) was applied for 45 min to block endogenous peroxidase activity.

Sections were microwaved in citric acid (10 mM, pH 6.0) for 12 min at 600 W and then incubated with primary antibodies (mouse anti-GRP78, 1:5000; goat anti-rat CYP1A1, 1:5000; goat anti-rat CYP3A2, 1:5000; goat anti-rat CYP2B1, 1:5000) in PBS pH 7.4 and 5% BSA overnight at 4°C, followed by treatment with secondary biotinylated rabbit anti-goat IgG or rabbit anti-mouse IgG (30 min, room temperature) and peroxidase-conjugated avidin (VECTASTAIN^®^ Elite ABC-Kit; Vector Laboratories, Burlingame, CA; another 30 min). The chromogen 3-amino-9-ethylcarbazole (AEC Substrate Pack; BioGenex, San Ramon, CA) was applied twice for 15 min to visualize immunoreactive sites. Sections were mounted in Vectamount™ mounting medium (Vector Laboratories, Burlingame, CA) and analyzed using an Axio Imager A1 microscope equipped with a 20× objective and a ProgRes C5 camera (Jenoptik, Jena, Germany).

### 5.17 Animal housing and treatment of mice with silybin hemisuccinate

Male C57BL/6 mice (12-weeks-old, body weight 25– 30 g; Charles River, Sulzfeld, Germany) were housed under standardized conditions with a day-night cycle of 12 h/12 h at 22 ± 1°C and 50 ± 10% environmental humidity. Standard diet and water were provided *ad libitum*. Animals were adapted to laboratory conditions before the experiment for at least 2 days. Silybin hemisuccinate (200 mg/kg) or vehicle (0.9% NaCl) were intraperitoneally administered trice (at 0, 12, and 24 h). Mice were anesthetized by isoflurane and sacrificed by isoflurane overdose after 37 h, and organs were removed, weighed and either fixed in 10% buffered formaldehyde or snap-frozen in liquid nitrogen for biochemical analysis. All experiments were performed in accordance with the German legislation on protection of animals and with approval of the Thuringian Animal Protection Committee.

### 5.18 GSH and GSSG levels

The tissue content of glutathione in its reduced (GSH) and oxidized (GSSG) form was analyzed by homogenizing the liver and kidney samples with eleven volumes of 0.2 M sodium phosphate buffer (5 mM EDTA; pH 8.0) and four volumes of 25% metaphosphoric acid. After centrifugation (12,000×g, 4°C, 30 min), the GSH content was measured in the supernatants using a colorimetric assay as previously described^208^. The GSSG concentration was assessed fluorometrically^209^.

### 5.19 Lipid peroxidation

To determine the tissue content of lipid peroxides as thiobarbituric acid reactive substances (TBARS), liver and kidney samples were homogenized in 19 volumes of ice-cold saline and analyzed fluorometrically^210^.

### 5.20 Biotransformation capacity

To obtain 9,000×g supernatants, the livers were homogenized in 0.1 M sodium phosphate buffer (pH 7.4) (1:2 w/v) and subsequently centrifuged at 9,000×g for 20 minutes at 4°C. Activities of all biotransformation reactions were assessed in these 9,000×g supernatants and referred to the protein content of this fraction which was determined with a modified Biuret method^211^. For assessment of CYP enzyme activities, the following model reactions were performed: benzyloxyresurofin-*O*-debenzylation (BROD)^212^, ethoxycoumarin-*O*-deethylation (ECOD)^213^, ethoxyresorufin-*O*-deethylation (EROD)^214^, ethylmorphine-*N*-demethylation (EMND)^215^, methoxyresorufin-*O*-demethylation (MROD)^214^, pentoxyresorufin-*O*-depentylation (PROD)^214^. Gst activities were determined using *o*-dinitrobenzene as a substrate. The resulting dinitrobenzene-glutathione conjugate was measured photometrically^216^. For the determination of Ugt activities, 4-methylumbelliferone was used as a substrate and the respective glucuronide was measured fluorometrically^217,218^.

### 5.12 Blood glucose levels

Blood glucose levels were determined using a commercially available blood glucose meter and respective test strips (BG star1, Sanofi-Aventis, Frankfurt, Germany).

### 5.13 Data analysis and statistics

Data are given as individual values and/or means ± SEM of *n* independent experiments. Statistical analysis was performed with GraphPad Prism 8.3 or 9.0 (GraphPad Software Inc, San Diego, CA, USA) using non-transformed or logarithmized data. Ordinary or repeated-measures one-way ANOVAs followed by Tukey *post-hoc* tests were applied for multiple comparison, and two-tailed Student’s *t*-tests were used for paired and unpaired observations (two-sided α levels of 0.05). Statistical significance was defined as *P* < 0.05 (*), *P* < 0.01 (**), and *P* < 0.001 (***). Outliers were determined by Grubb’s test. Figures were created with Graphpad Prism 8.3 or 9.0 (GraphPad Software Inc), Excel 2016 or 2020 (Microsoft, Redmond, WA), or Sigma Plot 13.0 (Systate Software GmbH, San Jose, CA).

## Abbreviations

ACC: acetyl-CoA carboxylase
ACOX3: acyl-CoA oxidase 3, pristanoyl
ACOT7: acyl-CoA thioesterase 7
ACSL6: acyl-CoA synthetase long chain family member 6
ATF6: activating transcription factor
ATGL: adipocyte triglyceride lipase
BSA: bovine serum albumin
CE: cholesteryl esters
CL: chemiluminescence
CPT1a: carnitine palmitoyl-transferase 1α
CYP: cytochrome P_450_
DAG: diacylglycerol
DGAT: diacylglycerol-O-acyltransferase
ER: endoplasmic reticulum
FASN: fatty acid synthase
GPAM: glycerol-3-phosphate acyltransferase, mitochondrial
GPAT: glycerophosphate acyltransferase
GPX: glutathione peroxidase
GSH: glutathione
GSSG: glutathione disulfide
GST: glutathione S-transferase
HAO2: hydroxyacid oxidase 2
HCV: hepatitis C virus
HSD17B13: 17-beta-hydroxysteroid dehydrogenase 13
LPIAT1: lysophosphatidylinositol acyltransferase 1
LPIN2: Lipin 2
LPAAT: lysophosphatidic acid acyltransferase
LPLAT: lysophospholipid acyltransferase
MBOAT7: membrane-bound O-acyltransferase domain-containing 7
MAFLD: metabolic dysfunction-associated steatotic liver disease
MASH: metabolic dysfunction-associated steatohepatitis
NAFLD: non-alcoholic fatty liver disease
NASH: non-alcoholic steatohepatitis
PC: phosphatidylcholine
PE: phosphatidylethanolamine
PG: phosphatidylglycerol
PI: phosphatidylinositol
PNPLA: patatin-like phospholipase domain-containing
PLA1A: phospholipases A1
PLA2: phospholipase A2
iPLA2/PLA2G6: calcium-independent phospholipase A2 / phospholipase A2 Group VI
PLA2G7: phospholipase A2 Group VII
PLD1: phospholipase D1
PRDX6: peroxiredoxin 6
PS: phosphatidylserine
ROS: reactive oxygen species
SCD5: stearoyl-CoA desaturase 5
SM: sphingomyelin
SREBF: sterol regulatory element binding transcription factor 1
TG: triglyceride; UGT; UDP glucuronosyltransferase
UPR: unfolded protein response
XBP1s: spliced form of X-box-binding protein 1.

## Author contributions

Solveigh Koeberle: Conceptualization, Data Curation, Formal analysis, Supervision,Visualization, Writing - Original Draft, Writing - Review & Editing. Maria Thürmer: Investigation, Methodology, Visualization, Writing - Review & Editing. Markus Werner: Investigation, Writing - Review & Editing. Julia Grander: Investigation, Visualization. Laura Hofer: Investigation. André Gollowitzer: Investigation, Writing - Review & Editing. Fengting Su: Investigation, Visualization. Loc Le Xuan: Investigation. Felix Benscheidt: Investigation. Ehsan Bonyadirad: Investigation. Armando Zarrelli: Methodology, Writing - Review & Editing. Giovanni Di Fabio: Methodology, Writing - Review & Editing. Oliver Werz: Writing - Review & Editing. Valeria Romanucci: Funding acquisition, Investigation, Methodology, Writing - Review & Editing. Amelie Lupp: Investigation, Methodology, Writing - Review & Editing. Andreas Koeberle: Conceptualization, Data Curation, Formal analysis, Funding acquisition, Project administration, Supervision, Visualization, Writing - Original Draft, Writing - Review & Editing.

## Declaration of Generative AI and AI-assisted technologies in the writing process

During the preparation of this work the authors used ChatGPT3.5 and DeepLWrite in order to improve readability and language. After using these tools, the authors reviewed and edited the content as needed and take full responsibility for the content of the publication.

## References

1 Rinella, M. E. et al. A multisociety Delphi consensus statement on new fatty liver disease nomenclature. Hepatology 78, 1966–1986, doi:10.1097/HEP.0000000000000520 (2023).

2 Loomba, R., Friedman, S. L. & Shulman, G. I. Mechanisms and disease consequences of nonalcoholic fatty liver disease. Cell 184, 2537–2564, doi:10.1016/j.cell.2021.04.015 (2021).

3 Meex, R. C. R. & Watt, M. J. Hepatokines: linking nonalcoholic fatty liver disease and insulin resistance. Nat Rev Endocrinol 13, 509–520, doi:10.1038/nrendo.2017.56 (2017).

4 Perry, R. J., Samuel, V. T., Petersen, K. F. & Shulman, G. I. The role of hepatic lipids in hepatic insulin resistance and type 2 diabetes. Nature 510, 84–91, doi:10.1038/nature13478 (2014).

5 Yamaguchi, K. et al. Inhibiting triglyceride synthesis improves hepatic steatosis but exacerbates liver damage and fibrosis in obese mice with nonalcoholic steatohepatitis. Hepatology 45, 1366–1374, doi:10.1002/hep.21655 (2007).

6 Tajmohammadi, A., Razavi, B. M. & Hosseinzadeh, H. Silybum marianum (milk thistle) and its main constituent, silymarin, as a potential therapeutic plant in metabolic syndrome: A review. Phytother Res 32, 1933–1949, doi:10.1002/ptr.6153 (2018).

7 Cheng, K. C. et al. Silymarin induces insulin resistance through an increase of phosphatase and tensin homolog in Wistar rats. PLoS One 9, e84550, doi:10.1371/journal.pone.0084550 (2014).

8 Ipsen, D. H., Lykkesfeldt, J. & Tveden-Nyborg, P. Molecular mechanisms of hepatic lipid accumulation in non-alcoholic fatty liver disease. Cell Mol Life Sci 75, 3313–3327, doi:10.1007/s00018-018-2860-6 (2018).

9 Younossi, Z. M. et al. Global epidemiology of nonalcoholic fatty liver disease-Meta-analytic assessment of prevalence, incidence, and outcomes. Hepatology 64, 73–84, doi:10.1002/hep.28431 (2016).

10 Baldini, F. et al. Biomechanics of cultured hepatic cells during different steatogenic hits. J Mech Behav Biomed Mater 97, 296–305, doi:10.1016/j.jmbbm.2019.05.036 (2019).

11 Feng, B. et al. Silymarin alleviates hepatic oxidative stress and protects against metabolic disorders in high-fat diet-fed mice. Free Radic Res 50, 314–327, doi:10.3109/10715762.2015.1116689 (2016).

12 Hellerbrand, C., Schattenberg, J. M., Peterburs, P., Lechner, A. & Brignoli, R. The potential of silymarin for the treatment of hepatic disorders. Clin Phytosci 2, doi:10.1186/s40816-016-0019-2 (2017).

13 Seebacher, F., Zeigerer, A., Kory, N. & Krahmer, N. Hepatic lipid droplet homeostasis and fatty liver disease. Semin Cell Dev Biol 108, 72–81, doi:10.1016/j.semcdb.2020.04.011 (2020).

14 Olzmann, J. A. & Carvalho, P. Dynamics and functions of lipid droplets. Nat Rev Mol Cell Biol 20, 137–155, doi:10.1038/s41580-018-0085-z (2019).

15 Baldini, F. et al. Aquaporin-9 is involved in the lipid-lowering activity of the nutraceutical silybin on hepatocytes through modulation of autophagy and lipid droplets composition. Biochim Biophys Acta Mol Cell Biol Lipids 1865, 158586, doi:10.1016/j.bbalip.2019.158586 (2020).

16 Su, F. & Koeberle, A. Regulation and targeting of SREBP-1 in hepatocellular carcinoma. Cancer Metastasis Rev, doi:10.1007/s10555-023-10156-5 (2023).

17 Wang, C. W. Lipid droplets, lipophagy, and beyond. Biochim Biophys Acta 1861, 793–805, doi:10.1016/j.bbalip.2015.12.010 (2016).

18 Vergani, L. Fatty Acids and Effects on In Vitro and In Vivo Models of Liver Steatosis. Curr Med Chem 26, 3439–3456, doi:10.2174/0929867324666170518101334 (2019).

19 Malhi, H. & Gores, G. J. Molecular mechanisms of lipotoxicity in nonalcoholic fatty liver disease. Semin Liver Dis 28, 360–369, doi:10.1055/s-0028-1091980 (2008).

20 Eslam, M., Valenti, L. & Romeo, S. Genetics and epigenetics of NAFLD and NASH: Clinical impact. J Hepatol 68, 268–279, doi:10.1016/j.jhep.2017.09.003 (2018).

21 Sookoian, S., Pirola, C. J., Valenti, L. & Davidson, N. O. Genetic Pathways in Nonalcoholic Fatty Liver Disease: Insights From Systems Biology. Hepatology 72, 330–346, doi:10.1002/hep.31229 (2020).

22 Chen, F. et al. Integrated Analysis of Key Genes and Pathways Involved in Nonalcoholic Steatohepatitis Improvement After Roux-en-Y Gastric Bypass Surgery. Front Endocrinol (Lausanne) 11, 611213, doi:10.3389/fendo.2020.611213 (2020).

23 Fuchs, C. D. et al. Hepatocyte-specific deletion of adipose triglyceride lipase (adipose triglyceride lipase/patatin-like phospholipase domain containing 2) ameliorates dietary induced steatohepatitis in mice. Hepatology 75, 125–139, doi:10.1002/hep.32112 (2022).

24 Teslovich, T. M. et al. Biological, clinical and population relevance of 95 loci for blood lipids. Nature 466, 707–713, doi:10.1038/nature09270 (2010).

25 Chamulitrat, W., Jansakun, C., Li, H. & Liebisch, G. Rescue of Hepatic Phospholipid Remodeling Defectin iPLA2beta-Null Mice Attenuates Obese but Not Non-Obese Fatty Liver. Biomolecules 10, doi:10.3390/biom10091332 (2020).

26 Kienesberger, P. C., Oberer, M., Lass, A. & Zechner, R. Mammalian patatin domain containing proteins: a family with diverse lipolytic activities involved in multiple biological functions. J Lipid Res 50 Suppl, S63–68, doi:10.1194/jlr.R800082-JLR200 (2009).

27 Kim, K. Y. et al. SREBP-2/PNPLA8 axis improves non-alcoholic fatty liver disease through activation of autophagy. Sci Rep 6, 35732, doi:10.1038/srep35732 (2016).

28 Shen, W. et al. PRDX6 Promotes Fatty Acid Oxidation via PLA2-Dependent PPARalpha Activation in Rats Fed High-Fat Diet. Antioxid Redox Signal 38, 1184–1200, doi:10.1089/ars.2022.0065 (2023).

29 Wang, H. et al. Inhibition of phospholipase D1 ameliorates hepatocyte steatosis and non-alcoholic fatty liver disease. JHEP Rep 5, 100726, doi:10.1016/j.jhepr.2023.100726 (2023).

30 Tanaka, Y. et al. LPIAT1/MBOAT7 depletion increases triglyceride synthesis fueled by high phosphatidylinositol turnover. Gut 70, 180–193, doi:10.1136/gutjnl-2020-320646 (2021).

31 Trepo, E. & Valenti, L. Update on NAFLD genetics: From new variants to the clinic. J Hepatol 72, 1196–1209, doi:10.1016/j.jhep.2020.02.020 (2020).

32 Abul-Husn, N. S. et al. A Protein-Truncating HSD17B13 Variant and Protection from Chronic Liver Disease. N Engl J Med 378, 1096–1106, doi:10.1056/NEJMoa1712191 (2018).

33 Paul, B., Lewinska, M. & Andersen, J. B. Lipid alterations in chronic liver disease and liver cancer. JHEP Rep 4, 100479, doi:10.1016/j.jhepr.2022.100479 (2022).

34 Ooi, G. J. et al. Hepatic lipidomic remodeling in severe obesity manifests with steatosis and does not evolve with non-alcoholic steatohepatitis. J Hepatol 75, 524–535, doi:10.1016/j.jhep.2021.04.013 (2021).

35 Masoodi, M. et al. Metabolomics and lipidomics in NAFLD: biomarkers and non-invasive diagnostic tests. Nat Rev Gastroenterol Hepatol 18, 835–856, doi:10.1038/s41575-021-00502-9 (2021).

36 Haberl, E. M. et al. Hepatic lipid profile in mice fed a choline-deficient, low-methionine diet resembles human non-alcoholic fatty liver disease. Lipids Health Dis 19, 250, doi:10.1186/s12944-020-01425-1 (2020).

37 Mannisto, V. et al. Total liver phosphatidylcholine content associates with non-alcoholic steatohepatitis and glycine N-methyltransferase expression. Liver Int 39, 1895–1905, doi:10.1111/liv.14174 (2019).

38 Deng, X. et al. iPLA2beta deficiency attenuates obesity and hepatic steatosis in ob/ob mice through hepatic fatty-acyl phospholipid remodeling. Biochim Biophys Acta 1861, 449–461, doi:10.1016/j.bbalip.2016.02.004 (2016).

39 Sanyal, A. J. & Pacana, T. A Lipidomic Readout of Disease Progression in A Diet-Induced Mouse Model of Nonalcoholic Fatty Liver Disease. Trans Am Clin Climatol Assoc 126, 271–288 (2015).

40 Eisinger, K. et al. Lipidomic analysis of the liver from high-fat diet induced obese mice identifies changes in multiple lipid classes. Exp Mol Pathol 97, 37–43, doi:10.1016/j.yexmp.2014.05.002 (2014).

41 Puri, P. et al. A lipidomic analysis of nonalcoholic fatty liver disease. Hepatology 46, 1081–1090, doi:10.1002/hep.21763 (2007).

42 Tvrdy, V. et al. Systematic review of pharmacokinetics and potential pharmacokinetic interactions of flavonolignans from silymarin. Med Res Rev 41, 2195–2246, doi:10.1002/med.21791 (2021).

43 Rainone, F. Milk thistle. Am Fam Physician 72, 1285–1288 (2005).

44 Abenavoli, L., Capasso, R., Milic, N. & Capasso, F. Milk thistle in liver diseases: past, present, future. Phytother Res 24, 1423–1432, doi:10.1002/ptr.3207 (2010).

45 Yan, T. et al. Herbal drug discovery for the treatment of nonalcoholic fatty liver disease. Acta Pharm Sin B 10, 3–18, doi:10.1016/j.apsb.2019.11.017 (2020).

46 Stolf, A. M., Cardoso, C. C. & Acco, A. Effects of Silymarin on Diabetes Mellitus Complications: A Review. Phytother Res 31, 366–374, doi:10.1002/ptr.5768 (2017).

47 Kren, V. Chirality Matters: Biological Activity of Optically Pure Silybin and Its Congeners. Int J Mol Sci 22, doi:10.3390/ijms22157885 (2021).

48 Milic, N., Milosevic, N., Suvajdzic, L., Zarkov, M. & Abenavoli, L. New therapeutic potentials of milk thistle (Silybum marianum). Nat Prod Commun 8, 1801–1810, doi:10.1177/1934578X1300801236 (2013).

49 Biedermann, D., Vavrikova, E., Cvak, L. & Kren, V. Chemistry of silybin. Nat Prod Rep 31, 1138–1157, doi:10.1039/c3np70122k (2014).

50 Fenclova, M. et al. Liquid chromatography-drift tube ion mobility-mass spectrometry as a new challenging tool for the separation and characterization of silymarin flavonolignans. Anal Bioanal Chem 412, 819–832, doi:10.1007/s00216-019-02274-3 (2020).

51 Feher, J. & Lengyel, G. Silymarin in the prevention and treatment of liver diseases and primary liver cancer. Curr Pharm Biotechnol 13, 210–217, doi:10.2174/138920112798868818 (2012).

52 Federico, A., Dallio, M. & Loguercio, C. Silymarin/Silybin and Chronic Liver Disease: A Marriage of Many Years. Molecules 22, doi:10.3390/molecules22020191 (2017).

53 Kalopitas, G. et al. Impact of Silymarin in individuals with nonalcoholic fatty liver disease: A systematic review and meta-analysis. Nutrition 83, 111092, doi:10.1016/j.nut.2020.111092 (2021).

54 Derakhshandeh-Rishehri, S. M., Heidari-Beni, M. & Eftekhari, M. H. The Effects of Realsil (Silybin-Phospholipid-Vitamin E Complex) on Liver Enzymes in Patients with Non-Alcoholic Fatty Liver Disease (Nafld) or Non-Alcoholic Steato-Hepatitis (Nash): A Systematic Review and Meta-Analysis of Rcts. Acta Endocrinol (Buchar) 16, 223–231, doi:10.4183/aeb.2020.223 (2020).

55 Anushiravani, A., Haddadi, N., Pourfarmanbar, M. & Mohammadkarimi, V. Treatment options for nonalcoholic fatty liver disease: a double-blinded randomized placebo-controlled trial. Eur J Gastroenterol Hepatol 31, 613–617, doi:10.1097/MEG.0000000000001369 (2019).

56 Ou, Q. et al. Silybin Alleviates Hepatic Steatosis and Fibrosis in NASH Mice by Inhibiting Oxidative Stress and Involvement with the Nf-kappaB Pathway. Dig Dis Sci 63, 3398–3408, doi:10.1007/s10620-018-5268-0 (2018).

57 Mahli, A. et al. Hepatoprotective effect of oral application of a silymarin extract in carbon tetrachloride-induced hepatotoxicity in rats. Clinical Phytoscience 1, 5, doi:10.1186/s40816-015-0006-z (2015).

58 Niu, H. et al. Prevention and management of idiosyncratic drug-induced liver injury: Systematic review and meta-analysis of randomised clinical trials. Pharmacol Res 164, 105404, doi:10.1016/j.phrs.2020.105404 (2021).

59 Saller, R., Brignoli, R., Melzer, J. & Meier, R. An updated systematic review with meta-analysis for the clinical evidence of silymarin. Forsch Komplementmed 15, 9–20, doi:10.1159/000113648 (2008).

60 Bahmani, M., Shirzad, H., Rafieian, S. & Rafieian-Kopaei, M. Silybum marianum: Beyond Hepatoprotection. J Evid Based Complementary Altern Med 20, 292–301, doi:10.1177/2156587215571116 (2015).

61 Fraschini, F., Demartini, G. & Esposti, D. Pharmacology of Silymarin. Clin. Drug Investig. 22, 51–65, doi:10.2165/00044011-200222010-00007 (2002).

62 Stiuso, P. et al. Serum oxidative stress markers and lipidomic profile to detect NASH patients responsive to an antioxidant treatment: a pilot study. Oxid Med Cell Longev 2014, 169216, doi:10.1155/2014/169216 (2014).

63 Bhattacharya, S. Phytotherapeutic properties of milk thistle seeds: An overview. Journal of Advanced Pharmacy Education & Research 1, 69–79 (2011).

64 Raina, K., Serkova, N. J. & Agarwal, R. Silibinin feeding alters the metabolic profile in TRAMP prostatic tumors: 1H-NMRS-based metabolomics study. Cancer Res 69, 3731–3735, doi:10.1158/0008-5472.CAN-09-0096 (2009).

65 Skottova, N. et al. Phenolics-rich extracts from Silybum marianum and Prunella vulgaris reduce a high-sucrose diet induced oxidative stress in hereditary hypertriglyceridemic rats. Pharmacol Res 50, 123–130, doi:10.1016/j.phrs.2003.12.013 (2004).

66 Poruba, M. et al. Improvement bioavailability of silymarin ameliorates severe dyslipidemia associated with metabolic syndrome. Xenobiotica 45, 751–756, doi:10.3109/00498254.2015.1010633 (2015).

67 Ni, X. & Wang, H. Silymarin attenuated hepatic steatosis through regulation of lipid metabolism and oxidative stress in a mouse model of nonalcoholic fatty liver disease (NAFLD). Am J Transl Res 8, 1073–1081 (2016).

68 Skottova, N. et al. Effect of silymarin on serum cholesterol levels in rats. Acta Univ Palacki Olomuc Fac Med 141, 87–89 (1998).

69 Huseini, H. F. et al. The efficacy of Silybum marianum (L.) Gaertn. (silymarin) in the treatment of type II diabetes: a randomized, double-blind, placebo-controlled, clinical trial. Phytother Res 20, 1036–1039, doi:10.1002/ptr.1988 (2006).

70 Cicero, A. F. G. et al. Lipid-lowering nutraceuticals in clinical practice: position paper from an International Lipid Expert Panel. Nutr Rev 75, 731–767, doi:10.1093/nutrit/nux047 (2017).

71 Sobolova, L., Skottova, N., Vecera, R. & Urbanek, K. Effect of silymarin and its polyphenolic fraction on cholesterol absorption in rats. Pharmacol Res 53, 104–112, doi:10.1016/j.phrs.2005.09.004 (2006).

72 Nassuato, G. et al. Effect of Silibinin on biliary lipid composition. Experimental and clinical study. J Hepatol 12, 290–295, doi:10.1016/0168-8278(91)90829-z (1991).

73 Voroneanu, L., Nistor, I., Dumea, R., Apetrii, M. & Covic, A. Silymarin in Type 2 Diabetes Mellitus: A Systematic Review and Meta-Analysis of Randomized Controlled Trials. J Diabetes Res 2016, 5147468, doi:10.1155/2016/5147468 (2016).

74 Lirussi, F. et al. Silybin-beta-cyclodextrin in the treatment of patients with diabetes mellitus and alcoholic liver disease. Efficacy study of a new preparation of an anti-oxidant agent. Diabetes Nutr Metab 15, 222–231 (2002).

75 Sun, W. L. et al. Microbially produced vitamin B12 contributes to the lipid-lowering effect of silymarin. Nat Commun 14, 477, doi:10.1038/s41467-023-36079-x (2023).

76 Schriewer, H. & Weinhold, F. The influence of silybin from Silybum marianum (L.) Gaertn. on in vitro phosphatidyl choline biosynthesis in rat livers. Arzneimittelforschung 29, 791–792 (1979).

77 Muriel, P. & Mourelle, M. Prevention by silymarin of membrane alterations in acute CCl4 liver damage. J Appl Toxicol 10, 275–279, doi:10.1002/jat.2550100408 (1990).

78 Schriewer, H., Lohmann, J. & Rauen, H. M. [The effect of silybin-dihemisuccinate on regulation disorders in phospholipid metabolism in acute galactosamine intoxication in the rat]. Arzneimittelforschung 25, 1582–1585 (1975).

79 Hermansson, M., Hokynar, K. & Somerharju, P. Mechanisms of glycerophospholipid homeostasis in mammalian cells. Prog Lipid Res 50, 240–257, doi:10.1016/j.plipres.2011.02.004 (2011).

80 Bhatt-Wessel, B., Jordan, T. W., Miller, J. H. & Peng, L. Role of DGAT enzymes in triacylglycerol metabolism. Arch Biochem Biophys 655, 1–11, doi:10.1016/j.abb.2018.08.001 (2018).

81 Christodoulou, E. et al. Serum and tissue pharmacokinetics of silibinin after per os and i.v. administration to mice as a HP-beta-CD lyophilized product. Int J Pharm 493, 366–373, doi:10.1016/j.ijpharm.2015.07.060 (2015).

82 Zhao, J. & Agarwal, R. Tissue distribution of silibinin, the major active constituent of silymarin, in mice and its association with enhancement of phase II enzymes: implications in cancer chemoprevention. Carcinogenesis 20, 2101–2108, doi:10.1093/carcin/20.11.2101 (1999).

83 Perona, J. S. Membrane lipid alterations in the metabolic syndrome and the role of dietary oils. Biochim Biophys Acta Biomembr 1859, 1690–1703, doi:10.1016/j.bbamem.2017.04.015 (2017).

84 van Meer, G., Voelker, D. R. & Feigenson, G. W. Membrane lipids: where they are and how they behave. Nat Rev Mol Cell Biol 9, 112–124, doi:10.1038/nrm2330 (2008).

85 Liu, Y., Xu, W., Zhai, T., You, J. & Chen, Y. Silibinin ameliorates hepatic lipid accumulation and oxidative stress in mice with non-alcoholic steatohepatitis by regulating CFLAR-JNK pathway. Acta Pharm Sin B 9, 745–757, doi:10.1016/j.apsb.2019.02.006 (2019).

86 Vecchione, G. et al. The Nutraceutic Silybin Counteracts Excess Lipid Accumulation and Ongoing Oxidative Stress in an In Vitro Model of Non-Alcoholic Fatty Liver Disease Progression. Front Nutr 4, 42, doi:10.3389/fnut.2017.00042 (2017).

87 Feng, R. et al. A combination of Pueraria lobata and Silybum marianum protects against alcoholic liver disease in mice. Phytomedicine 58, 152824, doi:10.1016/j.phymed.2019.152824 (2019).

88 Sun, R. et al. Silybin ameliorates hepatic lipid accumulation and modulates global metabolism in an NAFLD mouse model. Biomed Pharmacother 123, 109721, doi:10.1016/j.biopha.2019.109721 (2020).

89 Yang, L. et al. Silibinin improves nonalcoholic fatty liver by regulating the expression of miR122: An in vitro and in vivo study. Mol Med Rep 23, doi:10.3892/mmr.2021.11974 (2021).

90 Di Fabio, G., Romanucci, V., Di Marino, C., De Napoli, L. & Zarrelli, A. A rapid and simple chromatographic separation of diastereomers of silibinin and their oxidation to produce 2,3-dehydrosilybin enantiomers in an optically pure form. Planta Med 79, 1077–1080, doi:10.1055/s-0032-1328703 (2013).

91 Di Fabio, G. R. V.; De Nisco, M.; Pedatella, S.; Di Marino, C.; Zarrelli, A. Microwave-assisted oxidation of silibinin: a simple and preparative method for the synthesis of improved radical scavengers. Tetrahedron Lett. 54, 6279–6282, doi:10.1016/j.tetlet.2013.09.035 (2013).

92 Fallah, M. et al. Silymarin (milk thistle extract) as a therapeutic agent in gastrointestinal cancer. Biomed Pharmacother 142, 112024, doi:10.1016/j.biopha.2021.112024 (2021).

93 Yang, W. S. et al. Peroxidation of polyunsaturated fatty acids by lipoxygenases drives ferroptosis. Proc Natl Acad Sci U S A 113, E4966–4975, doi:10.1073/pnas.1603244113 (2016).

94 Wang, C. et al. Autophagy activated by silibinin contributes to glioma cell death via induction of oxidative stress-mediated BNIP3-dependent nuclear translocation of AIF. Cell Death Dis 11, 630, doi:10.1038/s41419-020-02866-3 (2020).

95 Lindholm, D., Korhonen, L., Eriksson, O. & Koks, S. Recent Insights into the Role of Unfolded Protein Response in ER Stress in Health and Disease. Front Cell Dev Biol 5, 48, doi:10.3389/fcell.2017.00048 (2017).

96 Malhi, H. & Kaufman, R. J. Endoplasmic reticulum stress in liver disease. J Hepatol 54, 795–809, doi:10.1016/j.jhep.2010.11.005 (2011).

97 Milacic, M. et al. The Reactome Pathway Knowledgebase 2024. Nucleic Acids Res 52, D672–D678, doi:10.1093/nar/gkad1025 (2024).

98 Hsiang, C. Y. et al. Glycyrrhizin, silymarin, and ursodeoxycholic acid regulate a common hepatoprotective pathway in HepG2 cells. Phytomedicine 22, 768–777, doi:10.1016/j.phymed.2015.05.053 (2015).

99 Lovelace, E. S. et al. Silymarin Suppresses Cellular Inflammation By Inducing Reparative Stress Signaling. J Nat Prod 78, 1990–2000, doi:10.1021/acs.jnatprod.5b00288 (2015).

100 DebRoy, S. et al. Hepatitis C virus dynamics and cellular gene expression in uPA-SCID chimeric mice with humanized livers during intravenous silibinin monotherapy. J Viral Hepat 23, 708–717, doi:10.1111/jvh.12551 (2016).

101 Badawi, A. F., Cavalieri, E. L. & Rogan, E. G. Role of human cytochrome P450 1A1, 1A2, 1B1, and 3A4 in the 2-, 4-, and 16alpha-hydroxylation of 17beta-estradiol. Metabolism 50, 1001–1003, doi:10.1053/meta.2001.25592 (2001).

102 Kisselev, P., Schunck, W. H., Roots, I. & Schwarz, D. Association of CYP1A1 polymorphisms with differential metabolic activation of 17beta-estradiol and estrone. Cancer Res 65, 2972–2978, doi:10.1158/0008-5472.CAN-04-3543 (2005).

103 Schwarz, D. et al. Arachidonic and eicosapentaenoic acid metabolism by human CYP1A1: highly stereoselective formation of 17(R),18(S)-epoxyeicosatetraenoic acid. Biochem Pharmacol 67, 1445–1457, doi:10.1016/j.bcp.2003.12.023 (2004).

104 Fer, M. et al. Cytochromes P450 from family 4 are the main omega hydroxylating enzymes in humans: CYP4F3B is the prominent player in PUFA metabolism. J Lipid Res 49, 2379–2389, doi:10.1194/jlr.M800199-JLR200 (2008).

105 Lucas, D. et al. Stereoselective epoxidation of the last double bond of polyunsaturated fatty acids by human cytochromes P450. J Lipid Res 51, 1125–1133, doi:10.1194/jlr.M003061 (2010).

106 Filali-Mouncef, Y. et al. The menage a trois of autophagy, lipid droplets and liver disease. Autophagy 18, 50–72, doi:10.1080/15548627.2021.1895658 (2022).

107 Gluchowski, N. L., Becuwe, M., Walther, T. C. & Farese, R. V., Jr. Lipid droplets and liver disease: from basic biology to clinical implications. Nat Rev Gastroenterol Hepatol 14, 343–355, doi:10.1038/nrgastro.2017.32 (2017).

108 Shi, H. et al. PPAR gamma Regulates Genes Involved in Triacylglycerol Synthesis and Secretion in Mammary Gland Epithelial Cells of Dairy Goats. PPAR Res 2013, 310948, doi:10.1155/2013/310948 (2013).

109 Petan, T. Lipid Droplets in Cancer. Rev Physiol Biochem Pharmacol, doi:10.1007/112_2020_51 (2020).

110 Tang, Y., Zhou, J., Hooi, S. C., Jiang, Y. M. & Lu, G. D. Fatty acid activation in carcinogenesis and cancer development: Essential roles of long-chain acyl-CoA synthetases. Oncol Lett 16, 1390–1396, doi:10.3892/ol.2018.8843 (2018).

111 Tong, L. Acetyl-coenzyme A carboxylase: crucial metabolic enzyme and attractive target for drug discovery. Cell Mol Life Sci 62, 1784–1803, doi:10.1007/s00018-005-5121-4 (2005).

112 Yamashita, A. et al. Acyltransferases and transacylases that determine the fatty acid composition of glycerolipids and the metabolism of bioactive lipid mediators in mammalian cells and model organisms. Prog Lipid Res 53, 18–81, doi:10.1016/j.plipres.2013.10.001 (2014).

113 Batchuluun, B., Pinkosky, S. L. & Steinberg, G. R. Lipogenesis inhibitors: therapeutic opportunities and challenges. Nat Rev Drug Discov, doi:10.1038/s41573-021-00367-2 (2022).

114 Kleiboeker, B. & Lodhi, I. J. Peroxisomal regulation of energy homeostasis: Effect on obesity and related metabolic disorders. Mol Metab 65, 101577, doi:10.1016/j.molmet.2022.101577 (2022).

115 Liang, D., Minikes, A. M. & Jiang, X. Ferroptosis at the intersection of lipid metabolism and cellular signaling. Mol Cell, doi:10.1016/j.molcel.2022.03.022 (2022).

116 Gan, B. Mitochondrial regulation of ferroptosis. J Cell Biol 220, doi:10.1083/jcb.202105043 (2021).

117 Araya, J. et al. Increase in long-chain polyunsaturated fatty acid n - 6/n - 3 ratio in relation to hepatic steatosis in patients with non-alcoholic fatty liver disease. Clin Sci (Lond) 106, 635–643, doi:10.1042/CS20030326 (2004).

118 Gomez-Lechon, M. J. et al. A human hepatocellular in vitro model to investigate steatosis. Chem Biol Interact 165, 106–116, doi:10.1016/j.cbi.2006.11.004 (2007).

119 Guguen-Guillouzo, C. & Guillouzo, A. General review on in vitro hepatocyte models and their applications. Methods Mol Biol 640, 1–40, doi:10.1007/978-1-60761-688-7_1 (2010).

120 da Silva, D. C., Valentao, P., Andrade, P. B. & Pereira, D. M. Endoplasmic reticulum stress signaling in cancer and neurodegenerative disorders: Tools and strategies to understand its complexity. Pharmacol Res 155, 104702, doi:10.1016/j.phrs.2020.104702 (2020).

121 Koeberle, A., Loser, K. & Thurmer, M. Stearoyl-CoA desaturase-1 and adaptive stress signaling. Biochim Biophys Acta 1861, 1719–1726, doi:10.1016/j.bbalip.2016.08.009 (2016).

122 Dallio, M. et al. PNPLA3, TM6SF2, and MBOAT7 Influence on Nutraceutical Therapy Response for Non-alcoholic Fatty Liver Disease: A Randomized Controlled Trial. Front Med (Lausanne) 8, 734847, doi:10.3389/fmed.2021.734847 (2021).

123 Federico, A. et al. Evaluation of the Effect Derived from Silybin with Vitamin D and Vitamin E Administration on Clinical, Metabolic, Endothelial Dysfunction, Oxidative Stress Parameters, and Serological Worsening Markers in Nonalcoholic Fatty Liver Disease Patients. Oxid Med Cell Longev 2019, 8742075, doi:10.1155/2019/8742075 (2019).

124 Federico, A. et al. The Bisphenol A Induced Oxidative Stress in Non-Alcoholic Fatty Liver Disease Male Patients: A Clinical Strategy to Antagonize the Progression of the Disease. Int J Environ Res Public Health 17, doi:10.3390/ijerph17103369 (2020).

125 Wah Kheong, C., Nik Mustapha, N. R. & Mahadeva, S. A Randomized Trial of Silymarin for the Treatment of Nonalcoholic Steatohepatitis. Clin Gastroenterol Hepatol 15, 1940–1949 e1948, doi:10.1016/j.cgh.2017.04.016 (2017).

126 Loguercio, C. et al. Silybin combined with phosphatidylcholine and vitamin E in patients with nonalcoholic fatty liver disease: a randomized controlled trial. Free Radic Biol Med 52, 1658–1665, doi:10.1016/j.freeradbiomed.2012.02.008 (2012).

127 Delmas, D., Xiao, J., Vejux, A. & Aires, V. Silymarin and Cancer: A Dual Strategy in Both in Chemoprevention and Chemosensitivity. Molecules 25, doi:10.3390/molecules25092009 (2020).

128 Moon, Y. J., Wang, X. & Morris, M. E. Dietary flavonoids: effects on xenobiotic and carcinogen metabolism. Toxicol In Vitro 20, 187–210, doi:10.1016/j.tiv.2005.06.048 (2006).

129 Kiruthiga, P. V., Karthikeyan, K., Archunan, G., Pandian, S. K. & Devi, K. P. Silymarin prevents benzo(a)pyrene-induced toxicity in Wistar rats by modulating xenobiotic-metabolizing enzymes. Toxicol Ind Health 31, 523–541, doi:10.1177/0748233713475524 (2015).

130 Kawaguchi-Suzuki, M. et al. The effects of milk thistle (Silybum marianum) on human cytochrome P450 activity. Drug Metab Dispos 42, 1611–1616, doi:10.1124/dmd.114.057232 (2014).

131 Upadhyay, G., Kumar, A. & Singh, M. P. Effect of silymarin on pyrogallol- and rifampicin-induced hepatotoxicity in mouse. Eur J Pharmacol 565, 190–201, doi:10.1016/j.ejphar.2007.03.004 (2007).

132 Zuber, R. et al. Effect of silybin and its congeners on human liver microsomal cytochrome P450 activities. Phytother Res 16, 632–638, doi:10.1002/ptr.1000 (2002).

133 Baer-Dubowska, W., Szaefer, H. & Krajka-Kuzniak, V. Inhibition of murine hepatic cytochrome P450 activities by natural and synthetic phenolic compounds. Xenobiotica 28, 735–743, doi:10.1080/004982598239155 (1998).

134 Sridar, C., Goosen, T. C., Kent, U. M., Williams, J. A. & Hollenberg, P. F. Silybin inactivates cytochromes P450 3A4 and 2C9 and inhibits major hepatic glucuronosyltransferases. Drug Metab Dispos 32, 587–594, doi:10.1124/dmd.32.6.587 (2004).

135 Dvorak, Z., Vrzal, R. & Ulrichova, J. Silybin and dehydrosilybin inhibit cytochrome P450 1A1 catalytic activity: a study in human keratinocytes and human hepatoma cells. Cell Biol Toxicol 22, 81–90, doi:10.1007/s10565-006-0017-0 (2006).

136 Beckmann-Knopp, S. et al. Inhibitory effects of silibinin on cytochrome P-450 enzymes in human liver microsomes. Pharmacol Toxicol 86, 250–256, doi:10.1111/j.0901-9928.2000.860602.x (2000).

137 Ahn, K., Szczesna-Skorupa, E. & Kemper, B. The amino-terminal 29 amino acids of cytochrome P450 2C1 are sufficient for retention in the endoplasmic reticulum. J Biol Chem 268, 18726–18733, doi:10.1016/S0021-9258(17)46690-7 (1993).

138 Galal, A. M., Walker, L. A. & Khan, I. A. Induction of GST and related events by dietary phytochemicals: sources, chemistry, and possible contribution to chemoprevention. Curr Top Med Chem 14, 2802–2821, doi:10.2174/1568026615666141208110721 (2015).

139 Maiorino, M., Conrad, M. & Ursini, F. GPx4, Lipid Peroxidation, and Cell Death: Discoveries, Rediscoveries, and Open Issues. Antioxid Redox Signal 29, 61–74, doi:10.1089/ars.2017.7115 (2018).

140 Bomgning, C. L. K. et al. Hepatoprotective effects of extracts, fractions and compounds from the stem bark of Pentaclethra macrophylla Benth: Evidence from in vitro and in vivo studies. Biomed Pharmacother 136, 111242, doi:10.1016/j.biopha.2021.111242 (2021).

141 Grasselli, E. et al. Excess fructose and fatty acids trigger a model of nonalcoholic fatty liver disease progression in vitro: Protective effect of the flavonoid silybin. Int J Mol Med 44, 705–712, doi:10.3892/ijmm.2019.4234 (2019).

142 Raza, H. Dual localization of glutathione S-transferase in the cytosol and mitochondria: implications in oxidative stress, toxicity and disease. FEBS J 278, 4243–4251, doi:10.1111/j.1742-4658.2011.08358.x (2011).

143 Polachi, N. et al. Modulatory effects of silibinin in various cell signaling pathways against liver disorders and cancer - A comprehensive review. Eur J Med Chem 123, 577–595, doi:10.1016/j.ejmech.2016.07.070 (2016).

144 Quiroga, A. D. & Lehner, R. Pharmacological intervention of liver triacylglycerol lipolysis: The good, the bad and the ugly. Biochem Pharmacol 155, 233–241, doi:10.1016/j.bcp.2018.07.005 (2018).

145 Reid, B. N. et al. Hepatic overexpression of hormone-sensitive lipase and adipose triglyceride lipase promotes fatty acid oxidation, stimulates direct release of free fatty acids, and ameliorates steatosis. J Biol Chem 283, 13087–13099, doi:10.1074/jbc.M800533200 (2008).

146 Kato, M., Higuchi, N. & Enjoji, M. Reduced hepatic expression of adipose tissue triglyceride lipase and CGI-58 may contribute to the development of non-alcoholic fatty liver disease in patients with insulin resistance. Scand J Gastroenterol 43, 1018–1019, doi:10.1080/00365520802008140 (2008).

147 Wu, J. W. et al. Deficiency of liver adipose triglyceride lipase in mice causes progressive hepatic steatosis. Hepatology 54, 122–132, doi:10.1002/hep.24338 (2011).

148 Grabner, G. F., Xie, H., Schweiger, M. & Zechner, R. Lipolysis: cellular mechanisms for lipid mobilization from fat stores. Nat Metab 3, 1445–1465, doi:10.1038/s42255-021-00493-6 (2021).

149 Gluchowski, N. L. et al. Hepatocyte Deletion of Triglyceride-Synthesis Enzyme Acyl CoA: Diacylglycerol Acyltransferase 2 Reduces Steatosis Without Increasing Inflammation or Fibrosis in Mice. Hepatology 70, 1972–1985, doi:10.1002/hep.30765 (2019).

150 Amin, N. B. et al. Targeting diacylglycerol acyltransferase 2 for the treatment of nonalcoholic steatohepatitis. Sci Transl Med 11, doi:10.1126/scitranslmed.aav9701 (2019).

151 Zhang, Z. et al. CGI1746 targets sigma(1)R to modulate ferroptosis through mitochondria-associated membranes. Nat Chem Biol, doi:10.1038/s41589-023-01512-1 (2024).

152 Romeo, S., Sanyal, A. & Valenti, L. Leveraging Human Genetics to Identify Potential New Treatments for Fatty Liver Disease. Cell Metab 31, 35–45, doi:10.1016/j.cmet.2019.12.002 (2020).

153 Liu, Z. et al. Association of Lipoprotein-Associated Phospholipase A2 with the Prevalence of Nonalcoholic Fatty Liver Disease: A Result from the APAC Study. Sci Rep 8, 10127, doi:10.1038/s41598-018-28494-8 (2018).

154 Sookoian, S. & Pirola, C. J. Systematic review with meta-analysis: risk factors for non-alcoholic fatty liver disease suggest a shared altered metabolic and cardiovascular profile between lean and obese patients. Aliment Pharmacol Ther 46, 85–95, doi:10.1111/apt.14112 (2017).

155 Luukkonen, P. K. et al. Distinct contributions of metabolic dysfunction and genetic risk factors in the pathogenesis of non-alcoholic fatty liver disease. J Hepatol 76, 526–535, doi:10.1016/j.jhep.2021.10.013 (2022).

156 Barbagallo, I. et al. Silibinin Regulates Lipid Metabolism and Differentiation in Functional Human Adipocytes. Front Pharmacol 6, 309, doi:10.3389/fphar.2015.00309 (2015).

157 Zhu, S. Y. et al. Silybum marianum oil attenuates hepatic steatosis and oxidative stress in high fat diet-fed mice. Biomed Pharmacother 100, 191–197, doi:10.1016/j.biopha.2018.01.144 (2018).

158 Schriewer, H. [Increase in rat liver synthesis of fatty acids and glycerol phospholipids following parenteral application of silybin-dihemisuccinate]. Arzneimittelforschung 28, 51–53 (1978).

159 Schriewer, H., Kramer, U., Rutkowski, G. & Borgis, K. J. [Influence of Silybin-dihemisuccinate on fatty acid synthesis in rat liver (author’s transl)]. Arzneimittelforschung 29, 524–526 (1979).

160 Cui, S. et al. Silybin alleviates hepatic lipid accumulation in methionine-choline deficient diet-induced nonalcoholic fatty liver disease in mice via peroxisome proliferator-activated receptor alpha. Chin J Nat Med 19, 401–411, doi:10.1016/S1875-5364(21)60039-0 (2021).

161 Kheiripour, N. et al. Silymarin prevents lipid accumulation in the liver of rats with type 2 diabetes via sirtuin1 and SREBP-1c. J Basic Clin Physiol Pharmacol 29, 301–308, doi:10.1515/jbcpp-2017-0122 (2018).

162 Pferschy-Wenzig, E. M. et al. Identification of isosilybin a from milk thistle seeds as an agonist of peroxisome proliferator-activated receptor gamma. J Nat Prod 77, 842–847, doi:10.1021/np400943b (2014).

163 Prakash, P. et al. Silymarin ameliorates fructose induced insulin resistance syndrome by reducing de novo hepatic lipogenesis in the rat. Eur J Pharmacol 727, 15–28, doi:10.1016/j.ejphar.2014.01.038 (2014).

164 Liu, Y., Yu, Q. & Chen, Y. Effect of silibinin on CFLAR-JNK pathway in oleic acid-treated HepG2 cells. Biomed Pharmacother 108, 716–723, doi:10.1016/j.biopha.2018.09.089 (2018).

165 Cui, C. X. et al. Silibinin Capsules improves high fat diet-induced nonalcoholic fatty liver disease in hamsters through modifying hepatic de novo lipogenesis and fatty acid oxidation. J Ethnopharmacol 208, 24–35, doi:10.1016/j.jep.2017.06.030 (2017).

166 Ka, S. O., Kim, K. A., Kwon, K. B., Park, J. W. & Park, B. H. Silibinin attenuates adipogenesis in 3T3-L1 preadipocytes through a potential upregulation of the insig pathway. Int J Mol Med 23, 633–637, doi:10.3892/ijmm_00000174 (2009).

167 Silva, C. M., Ferrari, G. D., Alberici, L. C., Malaspina, O. & Moraes, K. C. M. Cellular and molecular effects of silymarin on the transdifferentiation processes of LX-2 cells and its connection with lipid metabolism. Mol Cell Biochem 468, 129–142, doi:10.1007/s11010-020-03717-7 (2020).

168 Salamone, F. et al. Silibinin modulates lipid homeostasis and inhibits nuclear factor kappa B activation in experimental nonalcoholic steatohepatitis. Transl Res 159, 477–486, doi:10.1016/j.trsl.2011.12.003 (2012).

169 Vecchione, G. et al. Silybin counteracts lipid excess and oxidative stress in cultured steatotic hepatic cells. World J Gastroenterol 22, 6016–6026, doi:10.3748/wjg.v22.i26.6016 (2016).

170 Xie, Y. et al. Suppression of up-regulated LXRalpha by silybin ameliorates experimental rheumatoid arthritis and abnormal lipid metabolism. Phytomedicine 80, 153339, doi:10.1016/j.phymed.2020.153339 (2021).

171 Mourelle, M. & Franco, M. T. Erythrocyte defects precede the onset of CCl4-induced liver cirrhosis. Protection by silymarin. Life Sci 48, 1083–1090, doi:10.1016/0024-3205(91)90510-i (1991).

172 Schriewer, H. & Lohmann, J. [Disturbances in the regulation of phospholipid metabolism of the whole liver, mitochondria and microsomes in acute thioacetamide poisoning and the influcence of silymarin]. Arzneimittelforschung 26, 65–69 (1976).

173 Harayama, T. & Riezman, H. Understanding the diversity of membrane lipid composition. Nat Rev Mol Cell Biol 19, 281–296, doi:10.1038/nrm.2017.138 (2018).

174 Mateen, S., Tyagi, A., Agarwal, C., Singh, R. P. & Agarwal, R. Silibinin inhibits human nonsmall cell lung cancer cell growth through cell-cycle arrest by modulating expression and function of key cell-cycle regulators. Mol Carcinog 49, 247–258, doi:10.1002/mc.20595 (2010).

175 Kaur, M. et al. Silibinin suppresses growth and induces apoptotic death of human colorectal carcinoma LoVo cells in culture and tumor xenograft. Mol Cancer Ther 8, 2366–2374, doi:10.1158/1535-7163.MCT-09-0304 (2009).

176 Kauntz, H., Bousserouel, S., Gosse, F., Marescaux, J. & Raul, F. Silibinin, a natural flavonoid, modulates the early expression of chemoprevention biomarkers in a preclinical model of colon carcinogenesis. Int J Oncol 41, 849–854, doi:10.3892/ijo.2012.1526 (2012).

177 Valentine, W. J. et al. Update and nomenclature proposal for mammalian lysophospholipid acyltransferases, which create membrane phospholipid diversity. J Biol Chem 298, 101470, doi:10.1016/j.jbc.2021.101470 (2021).

178 Ramanathan, R. & Sivanesan, K. Evaluation of ameliorative ability of Silibinin against zidovudine and isoniazid-induced hepatotoxicity and hyperlipidaemia in rats: Role of Silibinin in Phase I and II drug metabolism. Chem Biol Interact 273, 142–153, doi:10.1016/j.cbi.2017.06.008 (2017).

179 Bernasconi, C. et al. Validation of in vitro methods for human cytochrome P450 enzyme induction: Outcome of a multi-laboratory study. Toxicol In Vitro 60, 212–228, doi:10.1016/j.tiv.2019.05.019 (2019).

180 Korobkova, E. A. Effect of Natural Polyphenols on CYP Metabolism: Implications for Diseases. Chem Res Toxicol 28, 1359–1390, doi:10.1021/acs.chemrestox.5b00121 (2015).

181 Villeneuve, J. P. & Pichette, V. Cytochrome P450 and liver diseases. Curr Drug Metab 5, 273–282, doi:10.2174/1389200043335531 (2004).

182 Pirmohamed, M., Madden, S. & Park, B. K. Idiosyncratic drug reactions. Metabolic bioactivation as a pathogenic mechanism. Clin Pharmacokinet 31, 215–230, doi:10.2165/00003088-199631030-00005 (1996).

183 Li, C., Lee, M. Y. & Choi, J. S. Effects of silybinin, CYP3A4 and P-glycoprotein inhibitor in vitro, on the bioavailability of loratadine in rats. Pharmazie 65, 510–514 (2010).

184 Tunca, R. et al. Pyridine induction of cytochrome P450 1A1, iNOS and metallothionein in Syrian hamsters and protective effects of silymarin. Exp Toxicol Pathol 61, 243–255, doi:10.1016/j.etp.2008.05.011 (2009).

185 Leber, H. W. & Knauff, S. Influence of silymarin on drug metabolizing enzymes in rat and man. Arzneimittelforschung 26, 1603–1605 (1976).

186 Zhu, H. J. et al. An assessment of pharmacokinetics and antioxidant activity of free silymarin flavonolignans in healthy volunteers: a dose escalation study. Drug Metab Dispos 41, 1679–1685, doi:10.1124/dmd.113.052423 (2013).

187 Wanwimolruk, S. & Prachayasittikul, V. Cytochrome P450 enzyme mediated herbal drug interactions (Part 1). EXCLI J 13, 347–391, doi:10.17877/DE290R-15628 (2014).

188 Rajnarayana, K., Reddy, M. S., Vidyasagar, J. & Krishna, D. R. Study on the influence of silymarin pretreatment on metabolism and disposition of metronidazole. Arzneimittelforschung 54, 109–113, doi:10.1055/s-0031-1296944 (2004).

189 Tighe, S. P., Akhtar, D., Iqbal, U. & Ahmed, A. Chronic Liver Disease and Silymarin: A Biochemical and Clinical Review. J Clin Transl Hepatol 8, 454–458, doi:10.14218/JCTH.2020.00012 (2020).

190 Xiao, F., Gao, F., Zhou, S. & Wang, L. The therapeutic effects of silymarin for patients with glucose/lipid metabolic dysfunction: A meta-analysis. Medicine (Baltimore) 99, e22249, doi:10.1097/MD.0000000000022249 (2020).

191 Pein, H. et al. Endogenous metabolites of vitamin E limit inflammation by targeting 5-lipoxygenase. Nat Commun 9, 3834, doi:10.1038/s41467-018-06158-5 (2018).

192 Koeberle, A. et al. SAR studies on curcumin’s pro-inflammatory targets: discovery of prenylated pyrazolocurcuminoids as potent and selective novel inhibitors of 5-lipoxygenase. J Med Chem 57, 5638–5648, doi:10.1021/jm500308c (2014).

193 Koeberle, A., Shindou, H., Harayama, T. & Shimizu, T. Role of lysophosphatidic acid acyltransferase 3 for the supply of highly polyunsaturated fatty acids in TM4 Sertoli cells. FASEB J 24, 4929–4938, doi:10.1096/fj.10-162818 (2010).

194 Koeberle, A., Shindou, H., Harayama, T., Yuki, K. & Shimizu, T. Polyunsaturated fatty acids are incorporated into maturating male mouse germ cells by lysophosphatidic acid acyltransferase 3. FASEB J 26, 169–180, doi:10.1096/fj.11-184879 (2012).

195 Espada, L. et al. Loss of metabolic plasticity underlies metformin toxicity in aged Caenorhabditis elegans. Nat Metab 2, 1316–1331, doi:10.1038/s42255-020-00307-1 (2020).

196 Koeberle, A. et al. Arachidonoyl-phosphatidylcholine oscillates during the cell cycle and counteracts proliferation by suppressing Akt membrane binding. Proc Natl Acad Sci U S A 110, 2546–2551, doi:10.1073/pnas.1216182110 (2013).

197 Koeberle, A. et al. Role of p38 mitogen-activated protein kinase in linking stearoyl-CoA desaturase-1 activity with endoplasmic reticulum homeostasis. FASEB J 29, 2439–2449, doi:10.1096/fj.14-268474 (2015).

198 Thürmer, M. et al. PI(18:1/18:1) is a SCD1-derived lipokine that limits stress signaling. Nat Commun 13, 2982, doi:10.1038/s41467-022-30374-9 (2022).

199 Liao, S. et al. alpha-Tocopherol-13’-Carboxychromanol Induces Cell Cycle Arrest and Cell Death by Inhibiting the SREBP1-SCD1 Axis and Causing Imbalance in Lipid Desaturation. Int J Mol Sci 24, doi:10.3390/ijms24119229 (2023).

200 van Pijkeren, A. et al. Proteome Coverage after Simultaneous Proteo-Metabolome Liquid-Liquid Extraction. J Proteome Res 22, 951–966, doi:10.1021/acs.jproteome.2c00758 (2023).

201 Glatzel, D. K. et al. Acetyl-CoA carboxylase 1 regulates endothelial cell migration by shifting the phospholipid composition. J Lipid Res 59, 298–311, doi:10.1194/jlr.M080101 (2018).

202 Ewels, P., Magnusson, M., Lundin, S. & Kaller, M. MultiQC: summarize analysis results for multiple tools and samples in a single report. Bioinformatics 32, 3047–3048, doi:10.1093/bioinformatics/btw354 (2016).

203 Dobin, A. et al. STAR: ultrafast universal RNA-seq aligner. Bioinformatics 29, 15–21, doi:10.1093/bioinformatics/bts635 (2013).

204 Anders, S., Pyl, P. T. & Huber, W. HTSeq--a Python framework to work with high-throughput sequencing data. Bioinformatics 31, 166–169, doi:10.1093/bioinformatics/btu638 (2015).

205 Love, M. I., Huber, W. & Anders, S. Moderated estimation of fold change and dispersion for RNA-seq data with DESeq2. Genome Biol 15, 550, doi:10.1186/s13059-014-0550-8 (2014).

206 Barrett, T. et al. NCBI GEO: archive for functional genomics data sets--update. Nucleic Acids Res 41, D991–995, doi:10.1093/nar/gks1193 (2013).

207 Lupp, A., Nagel, F. & Schulz, S. Reevaluation of sst(1) somatostatin receptor expression in human normal and neoplastic tissues using the novel rabbit monoclonal antibody UMB-7. Regul Pept 183, 1–6, doi:10.1016/j.regpep.2013.02.001 (2013).

208 Ellman, G. L. Tissue sulfhydryl groups. Arch Biochem Biophys 82, 70–77, doi:10.1016/0003-9861(59)90090-6 (1959).

209 Hissin, P. J. & Hilf, R. A fluorometric method for determination of oxidized and reduced glutathione in tissues. Anal Biochem 74, 214–226, doi:10.1016/0003-2697(76)90326-2 (1976).

210 Yagi, T. & Day, R. S., 3rd. Differential sensitivities of transformed and untransformed murine cell lines to DNA cross-linking agents relative to repair of O6-methylguanine. Mutat Res 184, 223–227, doi:10.1016/0167-8817(87)90020-4 (1987).

211 Klinger, W. & Muller, D. The influence of age on the protein concentration in serum, liver and kidney of rats determined by various methods. Z Versuchstierkd 16, 149–153 (1974).

212 Lubet, R. A. et al. Dealkylation of pentoxyresorufin: a rapid and sensitive assay for measuring induction of cytochrome(s) P-450 by phenobarbital and other xenobiotics in the rat. Arch Biochem Biophys 238, 43–48, doi:10.1016/0003-9861(85)90138-9 (1985).

213 Aitio, A. A simple and sensitive assay of 7-ethoxycoumarin deethylation. Anal Biochem 85, 488–491, (1978). doi:10.1016/0003-2697(78)90245-2

214 Pohl, R. J. & Fouts, J. R. A rapid method for assaying the metabolism of 7-ethoxyresorufin by microsomal subcellular fractions. Anal Biochem 107, 150–155, doi:10.1016/0003-2697(80)90505-9 (1980).

215 Kleeberg, U. & Klinger, W. Sensitive formaldehyde determination with Nash’s reagent and a ’tryptophan reaction’. J Pharmacol Methods 8, 19–31, doi:10.1016/0160-5402(82)90004-3 (1982).

216 Habig, W. H., Pabst, M. J. & Jakoby, W. B. Glutathione S-transferases. The first enzymatic step in mercapturic acid formation. J Biol Chem 249, 7130–7139 (1974).

217 Lilienblum, W., Walli, A. K. & Bock, K. W. Differential induction of rat liver microsomal UDP-glucuronosyltransferase activites by various inducing agents. Biochem Pharmacol 31, 907–913, doi:10.1016/0006-2952(82)90319-7 (1982).

218 Bock, K. W. et al. UDP-glucuronosyltransferase activities. Guidelines for consistent interim terminology and assay conditions. Biochem Pharmacol 32, 953–955, doi:10.1016/0006-2952(83)90610-x (1983).

