## Supplementary Information for "Silybin A from *Silybum marianum* reprograms lipid metabolism to induce a cell fate-dependent class switch from triglycerides to phospholipids"

### ORIGINAL ARTICLE

### Supplementary Figures

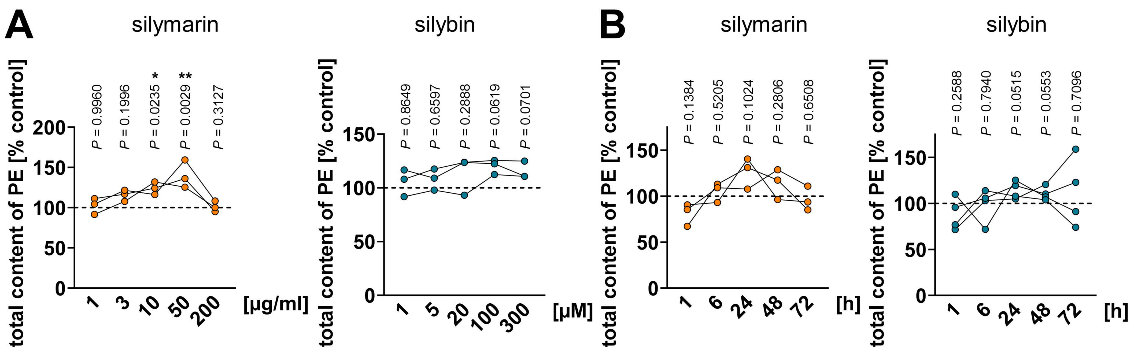


Figure S1. Concentration- and time-dependent effects of silymarin and silybin on cellular PE levels. (A, B) HepG2 cells were treated with silymarin, silybin or vehicle (ethanol for silymarin, DMSO for silybin) at the indicated concentrations for 24 h (A) or with silymarin (10 µg/ml), silybin (20 µM) or vehicle (ethanol for silymarin, DMSO for silybin) for the indicated incubation times (B). Independent datasets connected by lines; n = 3 (A, B, silymarin) or n = 4 (B, silybin). *$P < 0.05$, **$P < 0.01$ vs. vehicle control for the respective time point; two-tailed paired Student's *t*-test.

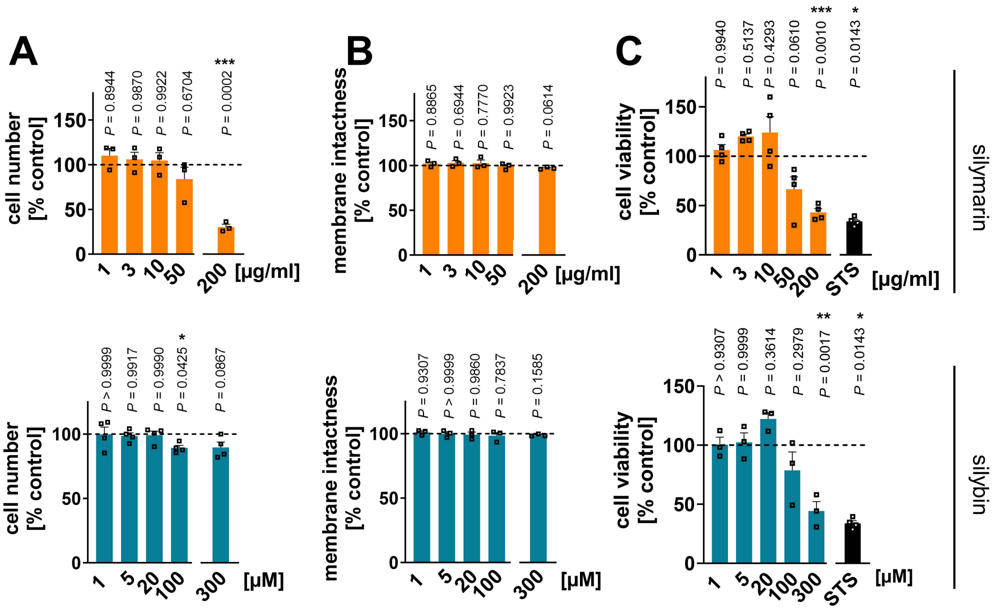


Figure S2. Effects of silymarin and silybin on cell number, membrane integrity and cell viability. HepG2 cells were treated with silymarin and silybin (at the indicated concentrations), staurosporine (STS, 1 µM) (C), or vehicle (ethanol for silymarin, DMSO for silybin and STS) for 24 h. (A) Cell numbers. Individual values and mean + SEM; n = 3 (silymarin), or n = 4 (silybin). (B) Membrane intactness measured by trypan blue staining. Individual values and mean + SEM; n = 3; effects of silymarin (200 µg/ml) and silybin (300 µM) were assessed in independent experiments (A, B). (C) Cell viability determined by MTT assay. Individual values and mean + SEM; n = 3 (silybin) or n = 4 (silymarin and STS). *$P < 0.05$, **$P < 0.01$, ***$P < 0.001$ vs. vehicle control; ordinary one-way ANOVA + Tukey HSD *post hoc* tests (C) and two-tailed paired Student's *t*-test (A and B, 200 µg/ml silymarin and 300 µM silybin, C, STS).

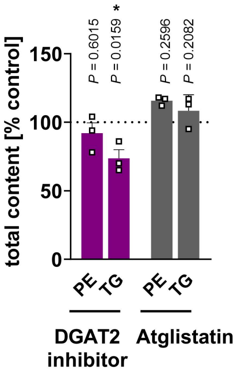


Figure S3. Influence of DGAT and ATGL inhibition on the phospholipid and TG content of hepatocytes. HepG2 cells were treated with the DGAT inhibitor PF 06424439 or the ATGL inhibitor atglistatin for 24-30 h. Total amounts of PE and TG were determined by UPLC-MS/MS. Individual values and mean + SEM; n = 3. *$P < 0.05$ vs. vehicle control; two-tailed unpaired Student's *t*-test.

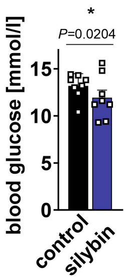


Figure S4. Blood glucose levels. Mice received silybin hemisuccinate (‘silybin’; 200 mg/kg, i.p.) or vehicle (0.9% NaCl) trice at 0, 12, and 24 h and were sacrificed after 37 h. Individual values and mean from n = 8 mice/group, *$P$ < 0.05 vs. vehicle control; two-tailed unpaired Student’s $t$-test.

# A

### monocytes

### PC

| | control | silymarin | control | silybin |
|---|---|---|---|---|
| PC(16:0/16:0) | 0.139 ± 0.045 | 0.170 ± 0.039 | 0.972 ± 0.060 | 1.053 ± 0.082 |
| PC(16:0/18:1) | 0.078 ± 0.029 | 0.125 ± 0.032 | 0.456 ± 0.015 | 0.524 ± 0.019 |
| PC(18:0/18:1) | 0.035 ± 0.011 | 0.052 ± 0.007 | 1.804 ± 0.069 | 2.064 ± 0.116 |
| PC(18:1/18:1) | 0.038 ± 0.015 | 0.059 ± 0.017 | 0.374 ± 0.035 | 0.440 ± 0.035 |
| PC(16:0/18:2) | 0.011 ± 0.003 | 0.024 ± 0.004 | 0.178 ± 0.007 | 0.232 ± 0.010 |
| PC(18:0/18:2) | 0.012 ± 0.004 | 0.022 ± 0.002 | 0.510 ± 0.010 | 0.566 ± 0.029 |
| PC(18:1/20:1) | 0.006 ± 0.003 | 0.009 ± 0.002 | 0.183 ± 0.020 | 0.182 ± 0.017 |
| PC(16:0/20:3) | 0.005 ± 0.002 | 0.007 ± 0.003 | 0.038 ± 0.004 | 0.046 ± 0.004 |
| PC(18:0/20:3) | 0.002 ± 0.001 | 0.003 ± 0.000 | 0.081 ± 0.016 | 0.101 ± 0.016 |
| PC(16:0/20:4) | 0.011 ± 0.004 | 0.019 ± 0.004 | 0.232 ± 0.008 | 0.248 ± 0.006 |
| PC(18:0/20:4) | 0.015 ± 0.005 | 0.023 ± 0.005 | 0.093 ± 0.003 | 0.105 ± 0.004 |
| PC(18:1/20:4) | 0.004 ± 0.001 | 0.007 ± 0.002 | 0.255 ± 0.013 | 0.257 ± 0.009 |
| PC(16:0/22:5) | 0.001 ± 0.000 | 0.002 ± 0.001 | 0.019 ± 0.002 | 0.048 ± 0.021 |
| PC(18:0/22:6) | 0.001 ± 0.000 | 0.004 ± 0.000 | 0.028 ± 0.001 | 0.024 ± 0.000 |
| PC(O-16:0/16:0) | 0.033 ± 0.009 | 0.040 ± 0.008 | 0.213 ± 0.025 | 0.226 ± 0.022 |
| PC(O-16:0/18:2) | 0.007 ± 0.002 | 0.009 ± 0.001 | 0.021 ± 0.002 | 0.021 ± 0.000 |

### PE

| | control | silymarin | control | silybin |
|---|---|---|---|---|
| PE(16:0/18:1) | 0.001 ± 0.000 | 0.002 ± 0.001 | 0.022 ± 0.002 | 0.037 ± 0.004 |
| PE(18:0/18:1) | 0.012 ± 0.005 | 0.017 ± 0.005 | 0.190 ± 0.007 | 0.204 ± 0.007 |
| PE(18:1/18:1) | 0.003 ± 0.001 | 0.004 ± 0.001 | 0.031 ± 0.003 | 0.049 ± 0.006 |
| PE(18:1/20:1) | 0.001 ± 0.000 | 0.001 ± 0.001 | 0.004 ± 0.000 | 0.005 ± 0.001 |
| PE(18:0/20:4) | 0.008 ± 0.003 | 0.013 ± 0.003 | 0.269 ± 0.013 | 0.328 ± 0.018 |
| PE(18:0/22:5) | 0.001 ± 0.000 | 0.001 ± 0.000 | 0.019 ± 0.001 | 0.022 ± 0.001 |
| PE(O-18:1/16:0) | 0.003 ± 0.001 | 0.003 ± 0.001 | 0.037 ± 0.004 | 0.043 ± 0.003 |

### PS

| | control | silymarin | control | silybin |
|---|---|---|---|---|
| PS(18:0/18:1) | 0.039 ± 0.012 | 0.061 ± 0.017 | 0.636 ± 0.036 | 0.726 ± 0.052 |
| PS(18:0/18:2) | 0.004 ± 0.001 | 0.006 ± 0.002 | 0.086 ± 0.006 | 0.103 ± 0.005 |
| PS(18:0/20:3) | 0.013 ± 0.005 | 0.020 ± 0.007 | 0.081 ± 0.008 | 0.093 ± 0.012 |
| PS(18:0/20:4) | 0.006 ± 0.001 | 0.012 ± 0.003 | 0.338 ± 0.044 | 0.442 ± 0.068 |

### PI

| | control | silymarin | control | silybin |
|---|---|---|---|---|
| PI(18:0/18:1) | 0.007 ± 0.002 | 0.011 ± 0.003 | 0.032 ± 0.002 | 0.032 ± 0.001 |
| PI(18:0/18:2) | 0.004 ± 0.001 | 0.008 ± 0.002 | 0.024 ± 0.000 | 0.024 ± 0.001 |
| PI(18:0/20:3) | 0.003 ± 0.001 | 0.005 ± 0.002 | 0.045 ± 0.003 | 0.052 ± 0.003 |
| PI(18:0/20:4) | 0.055 ± 0.016 | 0.094 ± 0.031 | 0.293 ± 0.011 | 0.334 ± 0.013 |

### PG

| | control | silymarin | control | silybin |
|---|---|---|---|---|
| PG(14:0/18:1) | 0.005 ± 0.003 | 0.006 ± 0.003 | 0.041 ± 0.006 | 0.088 ± 0.016 |
| PG(16:0/18:1) | 0.001 ± 0.000 | 0.004 ± 0.001 | 0.025 ± 0.003 | 0.021 ± 0.003 |
| PG(18:0/18:1) | 0.002 ± 0.000 | 0.005 ± 0.001 | 0.008 ± 0.001 | 0.010 ± ± 0.001 |

### SM

| | control | silymarin | control | silybin |
|---|---|---|---|---|
| 14:0 SM | 0.004 ± 0.001 | 0.007 ± 0.001 | 0.022 ± 0.002 | 0.025 ± 0.001 |
| 16:0 SM | 0.133 ± 0.044 | 0.191 ± 0.046 | 0.602 ± 0.019 | 0.630 ± 0.023 |
| 18:0 SM | 0.007 ± 0.002 | 0.009 ± 0.002 | 0.033 ± 0.002 | 0.041 ± 0.003 |

### TG

| | control | silymarin | control | silybin |
|---|---|---|---|---|
| 16:0/16:0/18:1 | 0.002 ± 0.000 | 0.001 ± 0.000 | 0.001 ± 0.000 | 0.001 ± 0.000 |
| 16:0/18:1/18:1 | 0.008 ± 0.002 | 0.005 ± 0.001 | 0.005 ± 0.001 | 0.005 ± 0.001 |
| 18:0/18:1/18:1 | 0.007 ± 0.001 | 0.005 ± 0.001 | 0.005 ± 0.001 | 0.005 ± 0.001 |
| 16:0/18:0/20:4 | 0.0005 ± 0.0001 | 0.0002 ± 0.000 | 0.0002 ± 0.0001 | 0.0001 ± 0.0000 |

# B

### HepG2

### PC

| | control | silymarin | control | silybin |
|---|---|---|---|---|
| PC(16:0/16:0) | 0.762 ± 0.058 | 0.943 ± 0.057 | 0.770 ± 0.055 | 0.799 ± 0.026 |
| PC(16:0/16:1) | 1.369 ± 0.125 | 2.123 ± 0.056 | 1.539 ± 0.049 | 1.765 ± 0.021 |
| PC(16:0/18:1) | 3.377 ± 0.246 | 4.940 ± 0.123 | 4.002 ± 0.149 | 4.654 ± 0.071 |
| PC(18:1/18:1) | 2.149 ± 0.170 | 3.275 ± 0.048 | 2.406 ± 0.109 | 3.047 ± 0.078 |
| PC(16:0/18:2) | 0.451 ± 0.044 | 0.698 ± 0.024 | 0.492 ± 0.048 | 0.499 ± 0.032 |

### PE

| | control | silymarin | control | silybin |
|---|---|---|---|---|
| PE(16:0/16:1) | 0.025 ± 0.002 | 0.041 ± 0.006 | 0.027 ± 0.002 | 0.032 ± 0.001 |
| PE(16:0/18:1) | 0.108 ± 0.011 | 0.182 ± 0.029 | 0.124 ± 0.006 | 0.156 ± 0.004 |
| PE(18:0/18:1) | 0.166 ± 0.021 | 0.279 ± 0.050 | 0.962 ± 0.077 | 1.389 ± 0.149 |
| PE(18:1/18:1) | 0.165 ± 0.014 | 0.256 ± 0.040 | 0.221 ± 0.006 | 0.272 ± 0.005 |
| PE(18:0/20:4) | 0.119 ± 0.014 | 0.167 ± 0.024 | 0.143 ± 0.007 | 0.171 ± 0.003 |

### PS

| | control | silymarin | control | silybin |
|---|---|---|---|---|
| PS(16:0/18:1) | 0.132 ± 0.014 | 0.187 ± 0.008 | 0.121 ± 0.007 | 0.129 ± 0.002 |
| PS(18:0/18:1) | 0.806 ± 0.092 | 1.056 ± 0.053 | 0.875 ± 0.012 | 1.047 ± 0.102 |
| PS(18:1/18:1) | 0.116 ± 0.015 | 0.147 ± 0.010 | 0.159 ± 0.007 | 0.151 ± 0.004 |
| PS(18:0/18:2) | 0.090 ± 0.010 | 0.122 ± 0.009 | 0.141 ± 0.013 | 0.137 ± 0.005 |

### PI

| | control | silymarin | control | silybin |
|---|---|---|---|---|
| PI(16:0/18:1) | 0.206 ± 0.013 | 0.316 ± 0.016 | 0.139 ± 0.001 | 0.141 ± 0.011 |
| PI(18:1/18:1) | 0.228 ± 0.014 | 0.383 ± 0.011 | 0.136 ± 0.006 | 0.164 ± 0.003 |
| PI(16:0/20:4) | 0.046 ± 0.004 | 0.059 ± 0.002 | 0.030 ± 0.005 | 0.027 ± 0.001 |
| PI(18:0/20:4) | 0.310 ± 0.034 | 0.415 ± 0.009 | 0.149 ± 0.028 | 0.140 ± 0.015 |

### PG

| | control | silymarin | control | silybin |
|---|---|---|---|---|
| PG(16:0/18:1) | 0.133 ± 0.006 | 0.149 ± 0.006 | 0.145 ± 0.022 | 0.112 ± 0.018 |
| PG(18:0/18:1) | 0.055 ± 0.002 | 0.081 ± 0.001 | 0.057 ± 0.006 | 0.071 ± 0.006 |
| PG(18:1/18:1) | 0.118 ± 0.001 | 0.176 ± 0.015 | 0.126 ± 0.016 | 0.136 ± 0.012 |

### SM

| | control | silymarin | control | silybin |
|---|---|---|---|---|
| 14:0 SM | 0.020 ± 0.002 | 0.020 ± 0.003 | 0.027 ± 0.003 | 0.030 ± 0.001 |
| 16:0 SM | 0.447 ± 0.035 | 0.650 ± 0.032 | 0.607 ± 0.050 | 0.661 ± 0.036 |
| 20:0 SM | 0.492 ± 0.036 | 0.788 ± 0.036 | 0.600 ± 0.049 | 0.629 ± 0.024 |
| 22:0 SM | 0.716 ± 0.055 | 1.300 ± 0.092 | 0.738 ± 0.028 | 0.909 ± 0.039 |

### TG

| | control | silymarin | control | silybin |
|---|---|---|---|---|
| 16:0/16:0/16:1 | 0.666 ± 0.122 | 0.712 ± 0.121 | 0.683 ± 0.122 | 0.401 ± 0.046 |
| 16:0/16:0/18:1 | 1.532 ± 0.265 | 1.649 ± 0.274 | 1.407 ± 0.275 | 0.962 ± 0.144 |
| 16:0/16:1/18:0 | 0.297 ± 0.054 | 0.325 ± 0.062 | 0.321 ± 0.096 | 0.173 ± 0.038 |
| 16:0/18:0/18:1 | 0.907 ± 0.163 | 1.051 ± 0.181 | 0.852 ± 0.252 | 0.469 ± 0.106 |
| 16:0/16:1/16:1 | 0.461 ± 0.080 | 0.497 ± 0.069 | 0.227 ± 0.033 | 0.130 ± 0.006 |
| 16:0/16:1/18:1 | 1.153 ± 0.186 | 1.244 ± 0.184 | 1.218 ± 0.144 | 0.743 ± 0.026 |
| 18:0/16:1/16:1 | 0.093 ± 0.017 | 0.101 ± 0.017 | 0.108 ± 0.023 | 0.055 ± 0.005 |
| 18:1/16:1/16:1 | 0.294 ± 0.057 | 0.298 ± 0.050 | 0.361 ± 0.057 | 0.209 ± 0.012 |
| 16:0/18:0/18:2 | 2.010 ± 0.311 | 2.229 ± 0.312 | 2.481 ± 0.199 | 1.567 ± 0.072 |
| 16:0/16:1/18:2 | 0.120 ± 0.028 | 0.114 ± 0.022 | 0.138 ± 0.030 | 0.080 ± 0.006 |
| 18:1/16:1/18:2 | 0.203 ± 0.051 | 0.188 ± 0.039 | 0.261 ± 0.051 | 0.155 ± 0.009 |
| 16:0/18:1/18:2 | 0.387 ± 0.081 | 0.396 ± 0.074 | 1.277 ± 0.333 | 0.817 ± 0.219 |
| 16:0/16:1/20:3 | 0.141 ± 0.034 | 0.129 ± 0.026 | 0.177 ± 0.039 | 0.105 ± 0.007 |

50 100 200 [%]

Figure S5. Phospholipid and TG profiles of human monocytes and hepatocytes upon treatment with silymarin or silybin. (A, B) Human primary monocytes (A) and HepG2 cells (B) were treated as described in Figure 1. Heatmap showing the absolute abundance of phospholipid and TG species normalized to the cell number (in relative units) as mean ± SEM. The color indicates percentage changes relative to control. Data and the number of experiments are identical to Figure 1.

**mouse liver**

**PC**

| | control | silybin |
|---|---|---|
| PC(16:0/16:0) | 4.376 ± 0.381 | 4.843 ± 0.135 |
| PC(16:0/18:1) | 11.962 ± 1.354 | 9.787 ± 0.730 |
| PC(18:0/18:1) | 5.659 ± 0.563 | 4.663 ± 0.228 |
| PC(18:1/18:1) | 2.663 ± 0.310 | 1.332 ± 0.085 |
| PC(16:0/18:2) | 18.908 ± 2.139 | 22.594 ± 1.350 |
| PC(18:0/18:2) | 9.555 ± 1.094 | 11.341 ± 0.520 |
| PC(16:0/20:3) | 5.633 ± 0.646 | 4.043 ± 0.350 |
| PC(18:0/20:3) | 1.118 ± 0.134 | 0.962 ± 0.059 |
| PC(16:0/20:4) | 9.452 ± 1.378 | 9.309 ± 0.586 |
| PC(18:0/20:4) | 6.412 ± 0.649 | 6.698 ± 0.324 |
| PC(18:1/20:4) | 1.683 ± 0.169 | 1.141 ± 0.079 |
| PC(16:0/22:5) | 0.889 ± 0.121 | 0.680 ± 0.046 |
| PC(18:0/22:6) | 3.222 ± 0.309 | 4.059 ± 0.108 |

**PE**

| | control | silybin |
|---|---|---|
| PE(16:0/18:1) | 4.624 ± 0.497 | 3.640 ± 0.317 |
| PE(18:0/18:1) | 0.817 ± 0.060 | 0.771 ± 0.033 |
| PE(18:1/18:1) | 1.625 ± 0.137 | 1.022 ± 0.056 |
| PE(16:0/18:2) | 8.006 ± 0.563 | 10.795 ± 0.745 |
| PE(18:0/18:2) | 5.675 ± 0.404 | 7.617 ± 0.359 |
| PE(18:1/18:2) | 4.152 ± 0.327 | 3.964 ± 0.179 |
| PE(16:0/20:3) | 2.062 ± 0.168 | 1.860 ± 0.167 |
| PE(16:0/20:4) | 14.876 ± 1.059 | 15.831 ± 0.973 |
| PE(18:0/20:4) | 34.085 ± 2.353 | 41.784 ± 1.493 |

**PS**

| | control | silybin |
|---|---|---|
| PS(16:0/20:4) | 5.386 ± 0.349 | 5.532 ± 0.339 |
| PS(18:0/20:4) | 34.183 ± 1.866 | 43.432 ± 1.849 |
| PS(18:0/22:6) | 21.078 ± 1.251 | 26.887 ± 1.185 |

**PI**

| | control | silybin |
|---|---|---|
| PI(16:0/18:1) | 100.794 ± 2.273 | 124.104 ± 6.368 |
| PI(16:0/20:3) | 1.332 ± 0.065 | 1.114 ± 0.122 |
| PI(16:0/20:4) | 16.510 ± 0.508 | 18.077 ± 1.512 |

**PG**

| | control | silybin |
|---|---|---|
| PG(16:0/18:1) | 8.349 ± 0.215 | 7.850 ± 0.308 |
| PG(18:1/18:1) | 0.961 ± 0.066 | 0.955 ± 0.079 |
| PG(16:0/18:2) | 2.789 ± 0.126 | 3.284 ± 0.180 |
| PG(18:1/18:2) | 0.605 ± 0.019 | 0.786 ± 0.044 |

**CE**

| | control | silybin |
|---|---|---|
| CE(16:0) | 0.0018 ± 0.0002 | 0.0027 ± 0.0003 |
| CE(18:0) | 0.0009 ± 0.0001 | 0.0014 ± 0.0002 |
| CE(16:1) | 0.0027 ± 0.0002 | 0.0029 ± 0.0004 |
| CE(18:1) | 0.0111 ± 0.0007 | 0.0133 ± 0.0023 |
| CE(18:2) | 0.0228 ± 0.0019 | 0.0303 ± 0.0025 |
| CE(18:3) | 0.0015 ± 0.0001 | 0.0017 ± 0.0002 |
| CE(20:4) | 0.0172 ± 0.0004 | 0.0328 ± 0.0041 |
| CE(22:6) | 0.0063 ± 0.0004 | 0.0146 ± 0.0019 |

**SM**

| | control | silybin |
|---|---|---|
| 16:0 SM | 2.239 ± 0.238 | 3.384 ± 0.233 |
| 20:0 SM | 47.372 ± 4.870 | 59.904 ± 3.841 |
| 22:0 SM | 20.677 ± 1.507 | 25.802 ± 1.851 |
| 22:1 SM | 11.541 ± 0.914 | 9.146 ± 0.632 |
| 22:2 SM | 27.720 ± 2.856 | 27.830 ± 1.704 |

**TG**

| | control | silybin |
|---|---|---|
| 16:0/16:0/16:0 | 0.099 ± 0.006 | 0.111 ± 0.006 |
| 16:0/16:0/16:1 | 0.196 ± 0.016 | 0.120 ± 0.005 |
| 16:0/16:0/18:1 | 1.068 ± 0.067 | 0.886 ± 0.054 |
| 16:0/16:1/18:0 | 0.025 ± 0.002 | 0.023 ± 0.002 |
| 16:0/16:1/16:1 | 0.117 ± 0.009 | 0.053 ± 0.008 |
| 16:0/16:1/18:1 | 0.782 ± 0.046 | 0.303 ± 0.024 |
| 16:0/18:1/18:1 | 5.118 ± 0.203 | 3.086 ± 0.163 |
| 18:1/16:1/16:1 | 0.088 ± 0.012 | 0.054 ± 0.012 |
| 16:0/16:0/18:2 | 1.614 ± 0.071 | 1.039 ± 0.107 |
| 16:0/18:0/18:2 | 3.419 ± 0.139 | 2.111 ± 0.107 |
| 16:0/16:1/18:2 | 0.562 ± 0.077 | 0.260 ± 0.059 |
| 16:0/18:1/18:2 | 3.364 ± 0.383 | 2.260 ± 0.376 |
| 18:0/18:1/18:2 | 0.310 ± 0.032 | 0.296 ± 0.033 |
| 18:1/16:1/18:2 | 0.305 ± 0.051 | 0.252 ± 0.064 |
| 16:0/18:2/18:2 | 1.797 ± 0.390 | 1.587 ± 0.387 |
| 16:0/18:0/20:3 | 0.216 ± 0.014 | 0.106 ± 0.009 |
| 16:0/16:1/20:3 | 0.199 ± 0.038 | 0.198 ± 0.056 |
| 16:0/16:0/20:4 | 1.273 ± 0.274 | 1.033 ± 0.237 |

50 100 150 [%]

Figure S6. Phospholipid, TG and CE profile of mouse liver upon gavage of silybin. Mice were treated as described in Figure 1. Heatmap showing the absolute abundance of phospholipid, TG and CE species normalized to the cell number (in relative units) as mean ± SEM. The color indicates percentage changes relative to control. Data and the number of experiments are identical to Figure 1.

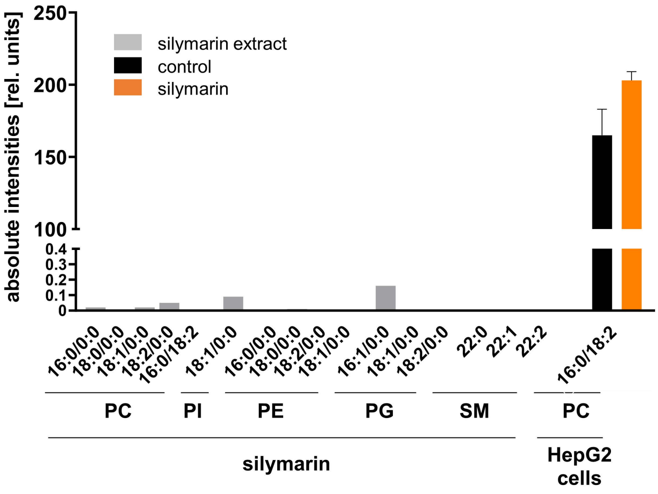


Figure S7. (Lyso)phospholipid content of silymarin in comparison to hepatocytes. Absolute abundance of (lyso)phospholipids in 10 µg silymarin or $3\times10^5$ HepG2 cells, which corresponds to the treatment of hepatocytes with 10 µg/ml under our experimental conditions. Lipid analysis of HepG2 cells focused on those species from Figure S5 and S6 that are present in silymarin, i.e., PC(16:0/18:2). Single value or mean + SEM; n = 1 (silymarin) or n = 3 (HepG2 cells).

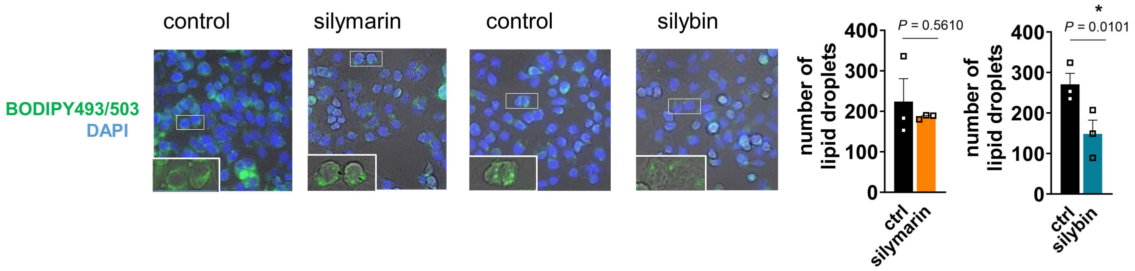


Figure S8. Impact of silymarin/silybin on lipid droplets. HepG2 cells were incubated with silymarin (10 µg/ml), silybin (20 µM) or vehicle control (ethanol for silymarin, DMSO for silybin) for 24 h. Immunofluorescence staining of lipid droplets with BODIPY493/503 (green); nuclei were visualized with DAPI (blue). Lipid droplets were counted in three independent experiments for 100 cells (each). Images are representative out of 3 independent experiments; magnified BODIPY493/503-stained section: 40×20 µm. Individual values and mean + SEM; n = 3. *$P < 0.05$ vs. vehicle control. Two-tailed paired Student's *t*-test.

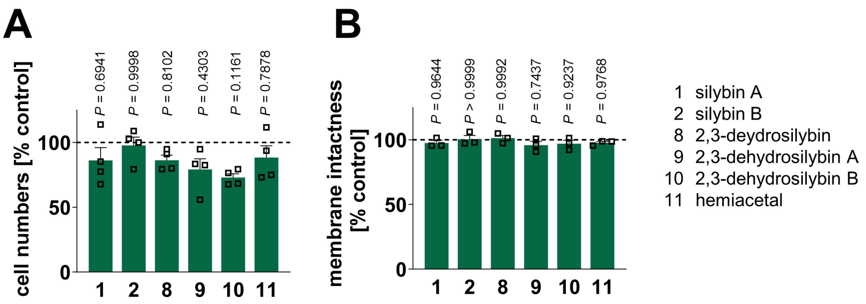


Figure S9. Effect of silybin derivatives on cell number and cell membrane intactness. HepG2 cells were treated with the indicated compounds (20 µM) or vehicle (DMSO) for 24 h. (A) Cell numbers; (B) membrane intactness measured by trypan blue staining. Individual values and mean + SEM; n = 3 (B) or n = 4 (A). *P* values vs. vehicle control; repeated measures one-way ANOVA + Tukey HSD *post hoc* tests.

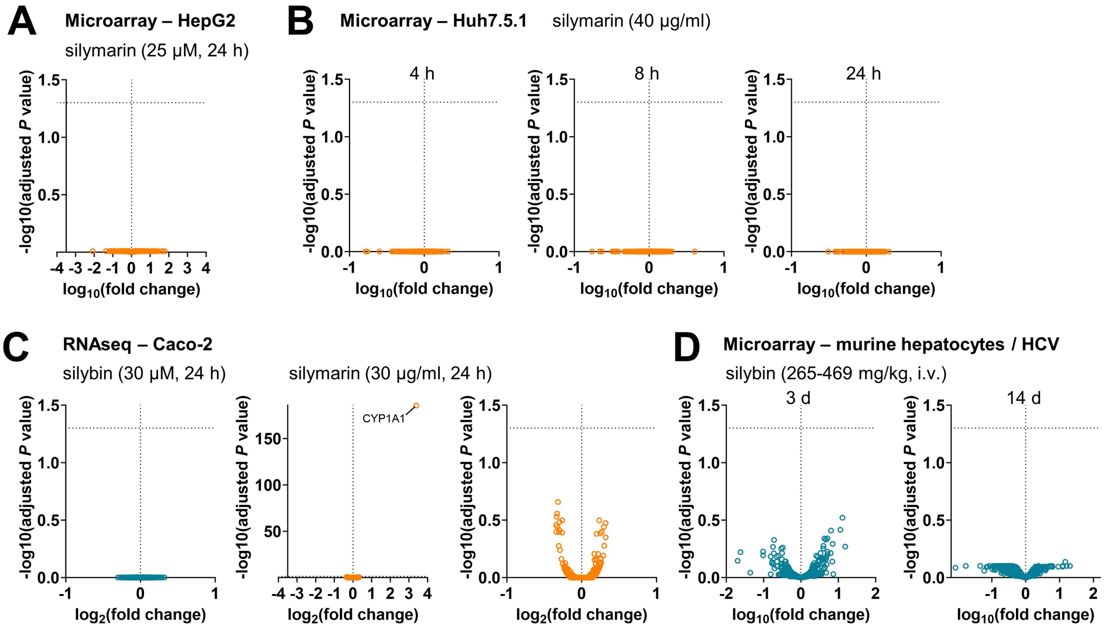


Figure S10. Volcano plots showing the data from Fig. 6A-D adjusted for multiple comparisons. Statistical calculations were performed by pairwise comparison of treatment and control groups using the GEO2R interactive webtool (https://www.ncbi.nlm.nih.gov/geo/geo2r/)[1].

Adjusted *P* values were calculated by multiple *t*-tests, with correction for multiple comparisons according to Benjamini and Hochberg (false discovery rate 5%) and autodetection for log-transformation. The dashed line indicates an adjusted *P* value of 0.05.

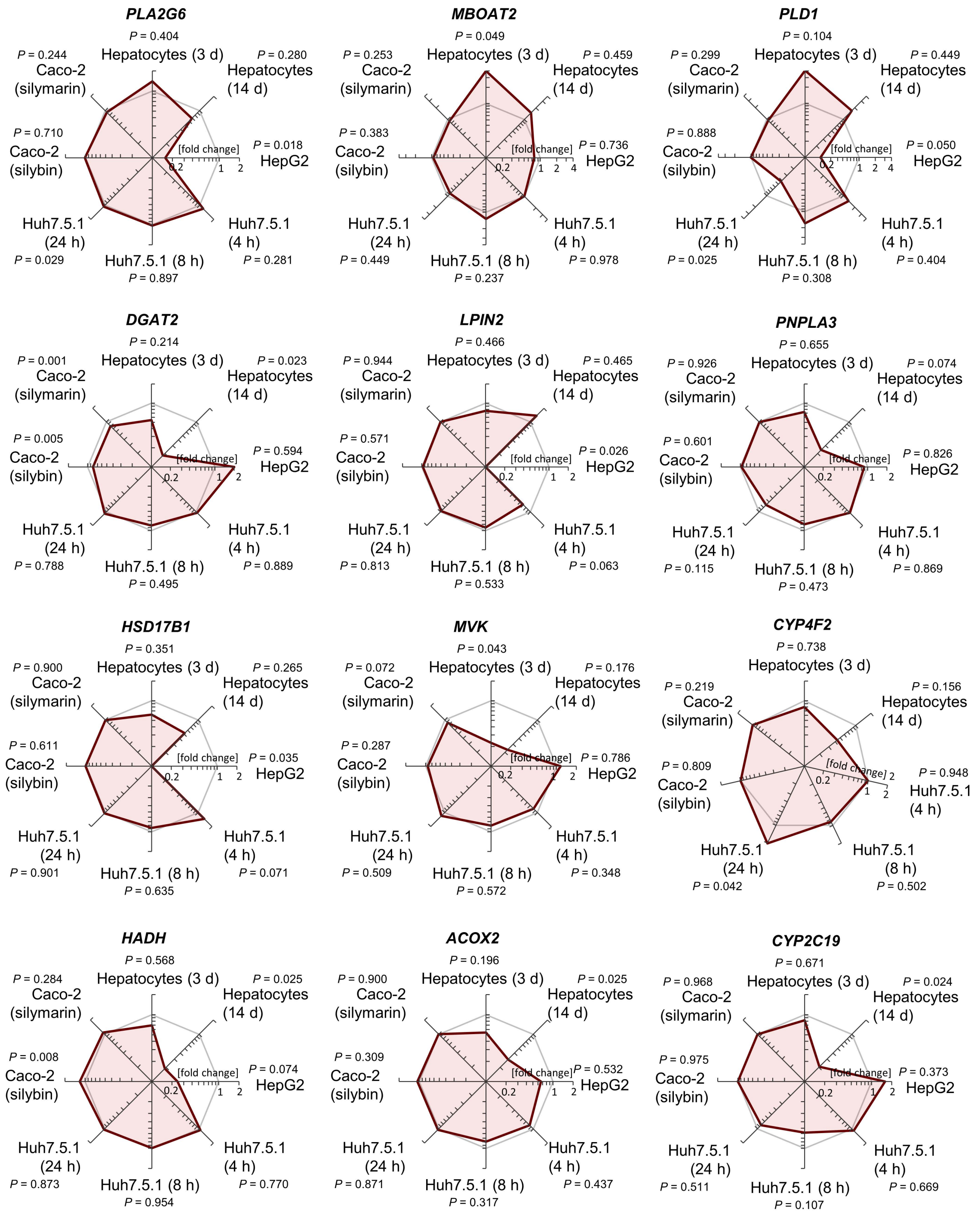


Figure S11. Lipid metabolic enzymes differentially regulated by silymarin/silybin. Comparative analysis of transcriptome data from silymarin-treated HepG2 and Huh7.5.1 hepatocarcinoma cells, silybin-and silymarin-treated CaCo-2 colon carcinoma cells, and hepatocytes derived from HCV-infected mice receiving silybin. Radar plots indicating the fold change in *PLA2G6*, *MBOAT2, PLD1, DGAT2, LPIN2, PNPLA3, HSH17B1, MVK, CYP4F2, HADH, ACOX2,* and *CYP2C19* expression by silymarin (HepG2, Huh7.5.1, CaCo-2) or silybin (hepatocytes, CaCo-2) relative to vehicle control. Non-adjusted *P* values given vs. vehicle control; multiple two-tailed unpaired Student's *t*-tests. Data are identical to Figure 6.

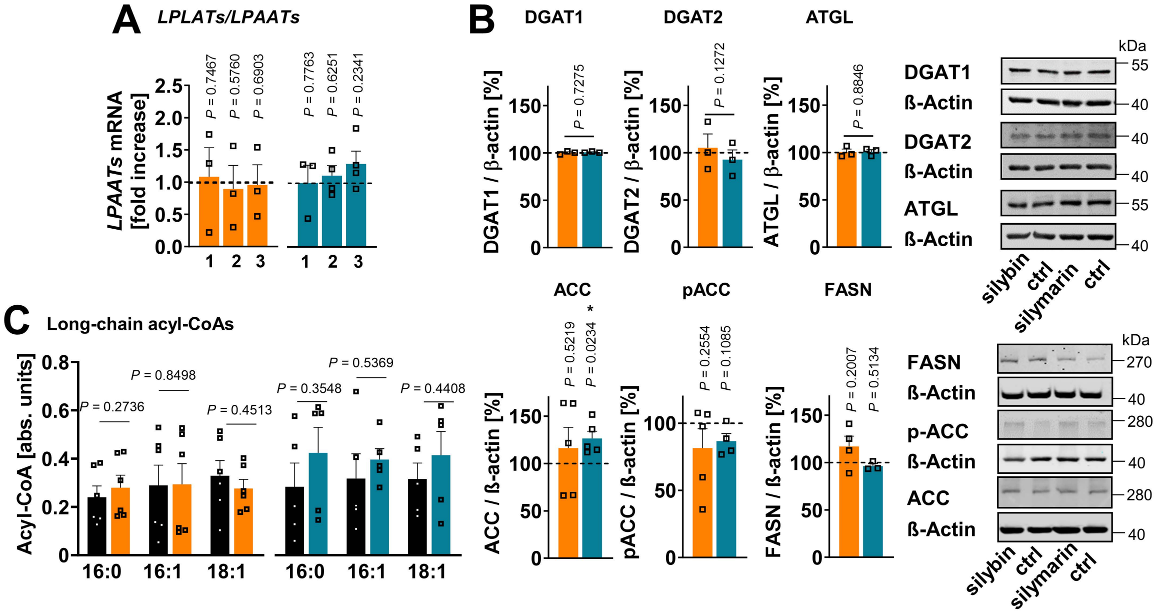


Figure S12. Remodeling of *de novo* phospholipid biosynthesis and TG metabolism. (A-C) HepG2 cells were incubated with silymarin (10 µg/ml), silybin (20 µM) or vehicle (ethanol for silymarin, DMSO for silybin) for 24 h; (A) mRNA levels of *LPLAT/LPAAT1-3* normalized to β-actin (*LPLATs/LPAATs* silybin) or *GAPDH* (*LPLATs/LPAATs* silymarin). Individual values and mean + SEM as fold-change of control; n = 3 (*LPLATs/LPAATs* silymarin, *LPLAT1/LPAAT1* silybin) and n = 4 (*LPLATs/LPAATs* silybin except *LPLAT1/LPAAT1*). (B) Protein expression of DGAT1, DGAT2, ATGL/PNPLA2, ACC1/2 and FASN, phosphorylation of ACC1/2. Individual values and mean + SEM; n = 3 (DGAT1, DGAT2, ATGL, FASN silybin), n = 4 (pACC silybin, FASN silymarin), and n = 5 (ACC, pACC silymarin). Individual values and mean + SEM; n = 3. Representative Western blots are shown. (C) Effects of silymarin and silybin on the cellular ratio of long-chain acyl-CoAs, normalized to the internal standard $[^{13}C_3]$-malonyl-CoA. Individual values and mean + SEM; n = 5 (silybin) and n = 6 (silymarin). *$P < 0.05$ vs. vehicle controls; two-tailed paired Student's t-tests.

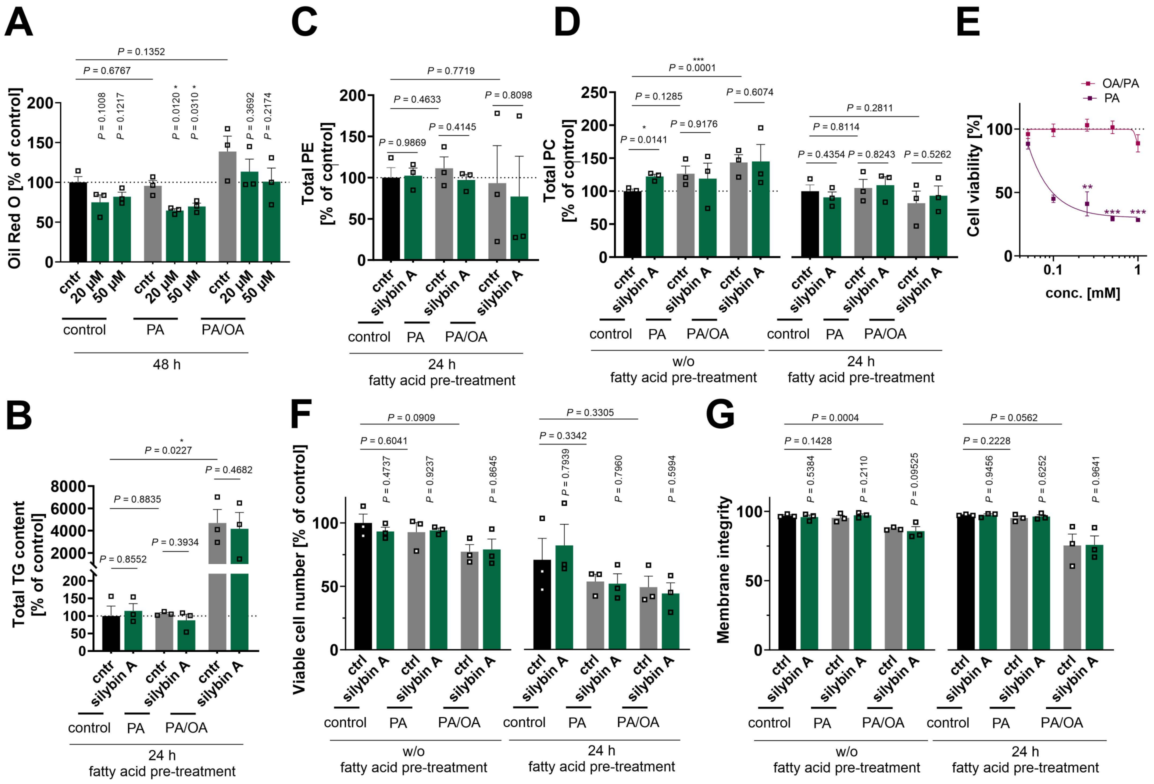


Figure S13. Comparison of the silybin A effects in non-challenged hepatocytes and cell-based disease models of MAFLD and acute lipotoxicity. (A) HepaRG cells were treated with 0.1 mM palmitate (PA) or a mixture of PA/oleate (OA) in a 1:2 ratio (in total 1 mM) together with vehicle (DMSO, 0.5%) or compounds for 48 h. Relative lipid droplet content was determined by Oil Red O staining. (B-D) HepaRG cells were either pre-treated with 0.1 mM palmitate (PA) or a mixture of PA/oleate (OA) in a 1:2 ratio (in total 1 mM) for 24 h followed by vehicle (DMSO, 0.5%) or silybin a (20 µM) treatment or cells were directly co-treated with vehicle (DMSO, 0.5%) or silybin A (20 µM), and the incubation was prolonged for a further 24 h. Total levels of TG (B), PE (C), and PC (D) determined by UPLC-MS/MS. (E) Cell viability measured by MTT assay. (F) Viable cell numbers. (G) Cell membrane integrity determined by trypan blue staining. Individual values and mean + SEM or ± SEM, n = 2 (E, 0.1 mM PA) or n = 3 (A-G, except D, 0.1 mM PA). *$P < 0.05$, ***$P < 0.001$ vs. control; two-tailed unpaired Student's *t*-test.

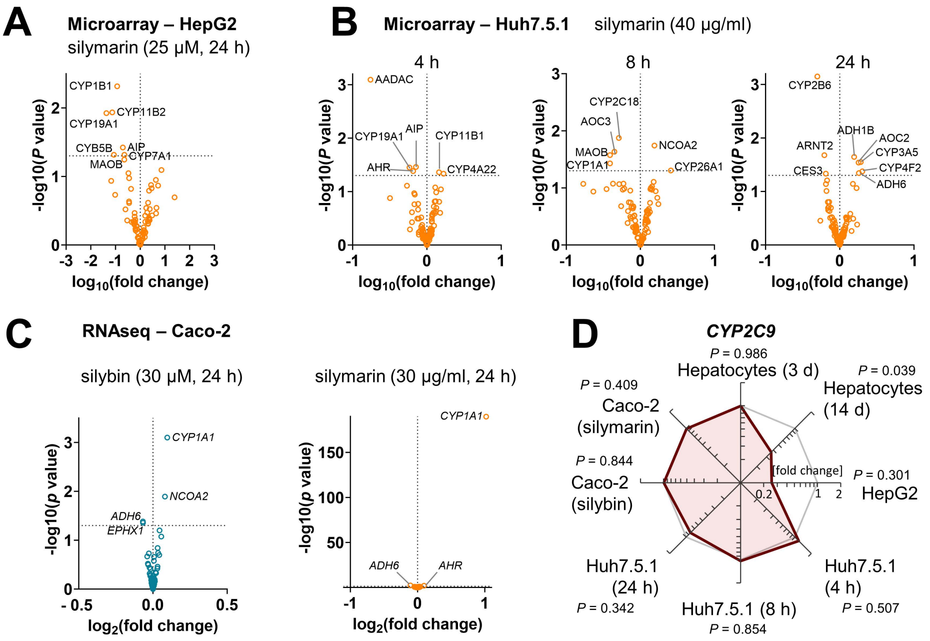


Figure S14. Silymarin/silybin kinetically controls the mRNA levels of genes involved in drug metabolism. Comparative analysis of transcriptome data from silymarin-treated HepG2 and Huh7.5.1 hepatocarcinoma cells, silybin- and silymarin-treated CaCo-2 colon carcinoma cells, and hepatocytes derived from HCV-infected mice receiving silybin. (A-C) Volcano plots compare the expression of proteins involved in drug metabolism upon silymarin (A-C) or silybin (C) treatment vs. vehicle control. Statistical calculations were performed by pairwise comparison of treatment and control groups using the GEO2R interactive webtool (https://www.ncbi.nlm.nih.gov/geo/geo2r/)[1]. *P*-values were calculated by multiple *t*-tests without correction for multiple comparisons. The dashed line indicates a *P*-value of 0.05. (D) Radar plots indicating the fold change in *CYP2C9* by silymarin (HepG2, Huh7.5.1, CaCo-2) or silybin (hepatocytes, CaCo-2) relative to vehicle control. Non-adjusted *P* values are given vs. vehicle control; multiple two-tailed unpaired Student's *t*-tests.

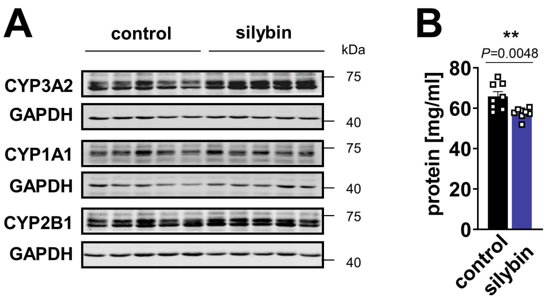


Figure S15. Impact of silybin on CYP expression in mouse liver. (A) Western blots for the densitometric data shown in Figure 8C are representative of n = 5 mice/group; (B) Protein concentration of the 9,000×g supernatant of the liver homogenate. Individual values and mean + SEM; n = 8 mice/group. $^{**}P < 0.01$ vs. vehicle control; two-tailed unpaired Student's *t*-test.

### Supplementary Notes

#### Supplementary Note 1: Cholesterol and CE metabolism

The effects of silybin/silymarin on CEs, the second major neutral lipid in lipid droplets after TGs, are less consistent. In some settings, silybin/silymarin stimulates cholesterol biosynthesis by upregulating the transcription factor *SREBP1* (Figure 6A) and repressing its endoplasmic reticulum anchor *INSIG1* (Figure 6D and E), thereby favoring the transfer of *SREBP1* to the Golgi for proteolytic procession to the mature form[2-4]. On the contrary, several SREBP-regulated target genes were downregulated (*ACACA, ELOVL6, MVK*, Figure 6A, E, Figure S11), except for FASN, which was upregulated as expected (Figure 6C). In other settings, silybin/silymarin reduces the expression of cholesterol biosynthetic enzymes (*MVK, TM7SF2*, Figure 6C, D, and E, and Figure 11). Such counter-regulation could be explained by initially increased levels of sterols, which then bind to the cholesterol sensor *INSIG1* (Figure S6 D and E) and suppress SREBP1 signaling (along with target protein expression) without necessarily interfering with SREBP1 expression[2]. In addition to canonical cholesterol biosynthesis, increased lipoprotein and sterol uptake (*LRP2*) (Figure 6B and C) and possibly endosomal cholesterol transport (*STARD3NL*) (Figure 6C) may further add to the accumulation of intracellular CE. In strong support of this hypothesis, silymarin administration to mice substantially elevated the hepatic CE content (Figure 1D). However, the increase in esterified cholesterol levels did not seem to be translated into enhanced cholesterol metabolism, as the expression of various sterol-metabolizing enzymes (*CYP1A1, CYP1B1, CYP2C8, CYP3A4, CYP7A1, CYP11B2, CYP17A1, CYP19A1, CYP27A1, ABCB11, SLC10A1, SLC27A5, HSD3B2, HSD17B1, HSD17B3, AKR1D1, AKR1C3, EBP, BAAT, STS*) is largely repressed (Figure 6A, B, C, D and E, and Figure S11), with a few exceptions (*CYP1A1, CYP1B1, CYP11B1, AKR1C3*, Figure 6B-D). Together, silymarin/silybin consistently downregulate anabolic and catabolic sterol metabolism, while exerting complex, partially opposite effects on cholesterol biosynthesis.

#### Supplementary Note 2: Vitamin A metabolism

In addition, silymarin/silybin differentially regulates a significant number of genes related to vitamin A metabolism, including retinoic acid biosynthesis (*CYP1B1, CYP3A4, ADH1B, ADH6,* Figure 8 A, Figure S14A-C), degradation (*CYP2C18, CYP3A5, CYP26A1, CYP2C8*, Figure 8A and B, Figure S14B), and vitamin A storage (*DGAT1, PNPLA3 and ATGL/ PNPLA2,* Figure 6A, B, E, and F, Figure S11).

Vitamin A is stored as retinyl esters in lipid droplets in the liver and shares common metabolic pathways with TG. This involves genetic risk factors for MASLD, such as *DGAT1, PNPLA3*, and *ATGL/PNPLA2*, which participate in retinol ester synthesis and degradation[5,6]. Vitamin A plays a central role in the regulation of hepatic lipid metabolism, including lipogenesis, lipid transport, and lipid catabolism[6], and disturbed vitamin A metabolism has been associated with MASLD[5,6]. It is therefore remarkable that our comparative transcriptome analysis revealed that numerous genes involved in vitamin A metabolism (including *CYP1B1, CYP3A4, ADH1B, ADH6, CYP2C18, CYP3A5, CYP26A1, CYP2C8, DGAT1, PNPLA2/ATGL* and possibly *PNPLA3*) are subject to regulation by silybin/silymarin. Further studies are needed to explore whether the interference with vitamin A metabolism by silymarin/silybin affects hepatic lipid metabolism and influences the development and/or progression of MASLD.

### References


1 Barrett, T. *et al.* NCBI GEO: archive for functional genomics data sets--update. *Nucleic Acids Res* **41**, D991-995, doi:10.1093/nar/gks1193 (2013).

2 Yang, T. *et al.* Crucial step in cholesterol homeostasis: sterols promote binding of SCAP to INSIG-1, a membrane protein that facilitates retention of SREBPs in ER. *Cell* **110**, 489-500, doi:10.1016/s0092-8674(02)00872-3 (2002).

3 Engelking, L. J., Cantoria, M. J., Xu, Y. & Liang, G. Developmental and extrahepatic physiological functions of SREBP pathway genes in mice. *Semin Cell Dev Biol* **81**, 98-109, doi:10.1016/j.semcdb.2017.07.011 (2018).

4 Chen, Y. *et al.* Maturation and activity of sterol regulatory element binding protein 1 is inhibited by acyl-CoA binding domain containing 3. *PLoS One* **7**, e49906, doi:10.1371/journal.pone.0049906 (2012).

5 Saeed, A. *et al.* Impaired Hepatic Vitamin A Metabolism in NAFLD Mice Leading to Vitamin A Accumulation in Hepatocytes. *Cell Mol Gastroenterol Hepatol* **11**, 309-325 e303, doi:10.1016/j.jcmgh.2020.07.006 (2021).

6 Saeed, A., Dullaart, R. P. F., Schreuder, T., Blokzijl, H. & Faber, K. N. Disturbed Vitamin A Metabolism in Non-Alcoholic Fatty Liver Disease (NAFLD). *Nutrients* **10**, doi:10.3390/nu10010029 (2017).
